# X-Plat: a polynomial regression–based tool for cross-platform transformation of expression and methylation data

**DOI:** 10.64898/2026.02.22.707273

**Authors:** Neeraja M. Krishnan, Sarah I Rahman, Lars Røn Olsen, Binay Panda

## Abstract

Many biological studies could benefit from combining data from legacy microarray and high-throughput sequencing platforms, especially in clinical domains where collecting additional samples is not possible. However, incompatibility between platforms makes legacy data difficult to integrate, owing to differences in platform design, target preparation, and dependence on prior annotations. Here, we describe X-Plat, a cross-platform data transformation tool for both expression and methylation assays that inter-converts data between microarray and sequencing platforms using per-gene second-degree polynomial regression. X-Plat learns cross-platform conversion rules from paired microarray–sequencing datasets spanning multiple conditions, sample sources, organisms, and platforms, and evaluates performance using cross-validated root mean square error (RMSE) per gene. In rat, *Arabidopsis*, and human datasets, X-Plat achieved lower cross-validated RMSE than TDM, HARMONY, and HARMONY2 for the vast majority of genes (≥95% in all sequencing-to-array transformations and most array-to-sequencing transformations, with ∼82% in the *Arabidopsis* array-to-sequencing setting), and these findings were confirmed using RMSE on held-out test samples from the first cross-validation fold. X-Plat also achieved low RMSE (≤0.2) for the majority of CpG regions in paired human array–sequencing methylation datasets. Using X-Plat, users can transform data between microarray and high-throughput sequencing platforms, enabling cross-platform comparison and reuse of legacy cohorts.

**GRAPHICAL ABSTRACT:** 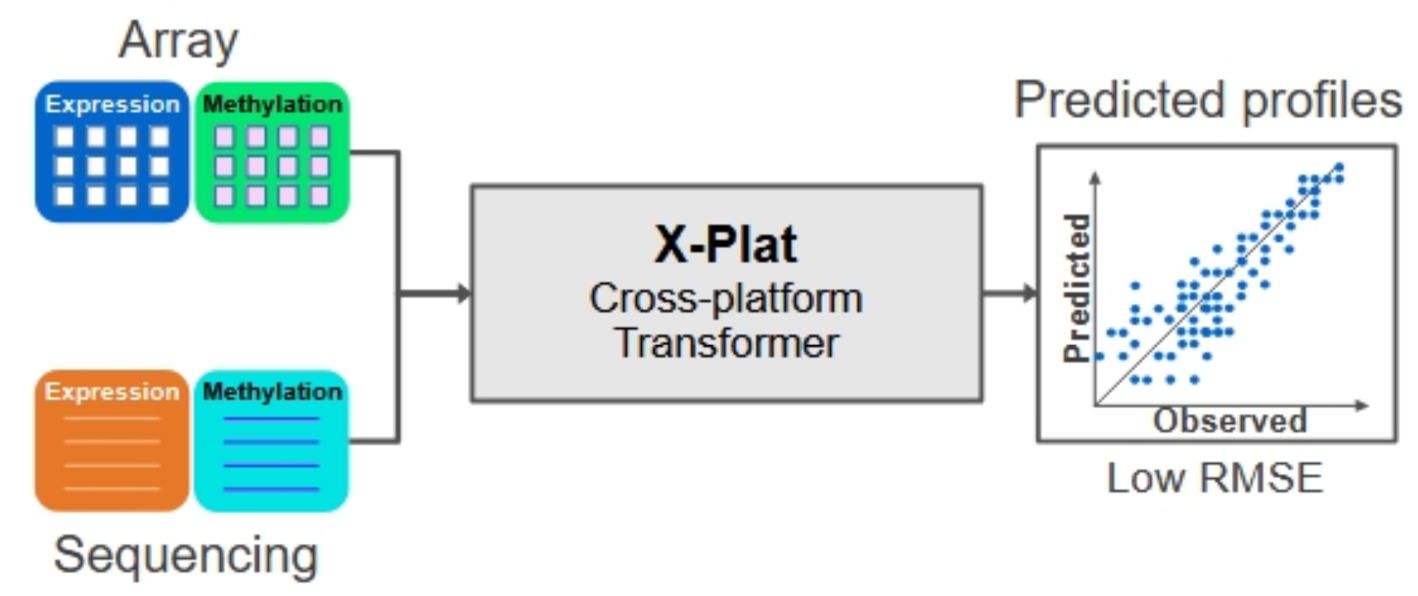

## INTRODUCTION

DNA microarrays revolutionized high-throughput data generation and were the primary mode of measuring genome-wide gene expression, genetic polymorphisms, and promoter methylation for the better part of the last three decades. Over the last decade, next-generation DNA sequencing technology has increasingly replaced microarrays for genome-wide measurement of DNA, RNA, and epigenetic changes and is favored over array-based platforms because of its unbiased coverage and reduced dependence on prior sequencing and annotation information (1, 2). However, a full transition from array-based methods to sequencing-based ones, although imminent, faces hurdles due to higher cost, limited access to instruments and assay know-how and the need for a skilled workforce trained to perform data analysis. Even today, many laboratories rely on microarrays to generate data while gradually increasing their use of sequencing-based assays (3, 4).

Legacy datasets generated using microarrays and deposited in public databases could be immensely useful for validating results obtained with sequencing-based assays, especially in clinical domains where sample numbers and accessibility are major constraints. For this reason, the ability to convert data from microarray-based formats to sequencing-based formats, and vice versa, is not just beneficial but increasingly necessary to continue using legacy data (5, 6).

Past efforts have produced cross-platform normalization and transformation tools to leverage legacy data, including methods for batch correction, probe-level alignment, transcriptomic distribution matching and methylation remapping (7–10). In one prior work, a two-stage Bayesian mixture modeling strategy was applied to assimilate and analyze four independent microarray studies to derive an inter-study-validated meta-signature that unified disparate gene expression data on a common probability scale, enabling robust prognostic signatures (11). ComBat is a widely used batch correction tool for array data that integrates data from multiple experiments and platforms (7). ComBat can also be used for cross-platform integration by treating data from different platforms as different batches; however, it merges mean and variance across batches (inter-batch) while ignoring within-batch mean and variance, and an extension that accounts for single-batch mean and variance was subsequently proposed (12). Probe mapping across microarray platforms has been used as a crucial step for data integration (13). The methyLiftover method maps DNA methylation data from bisulfite sequencing approaches to CpG sites measured with Illumina methylation bead-array platforms, thereby enabling cross-platform DNA methylation data integration (9). A comparison of microarray platform integration approaches revealed that methods relying on actual probe-set signal intensities are superior to those considering only biological characteristics of probe sequences and that cross-platform integration improves correlation with RNA-seq data (14). Methods that transform RNA-seq data to make it compatible with microarray-based models have also been proposed (8, 15, 16), and bidirectional conversion is possible and has been demonstrated (5). Other cross-platform normalization tools, such as MatchMixeR, model platform differences using gene-specific linear mixed-effects regression estimated from paired profiles, and can effectively reduce cross-platform discrepancies for expression data (17). In contrast, our goal here is to evaluate a simple per-gene polynomial mapping that supports explicit bidirectional conversion between platforms and is readily extensible to non-expression modalities such as DNA methylation. However, neither explicit, reusable gene-specific conversion rules nor a universal conversion rule between platforms has been developed for routine forward and reverse transformation of both expression and methylation profiles. Solving this key problem and providing a platform to integrate legacy data with modern high-throughput data would greatly benefit the community.

Here, we describe a cross-platform data transformation tool, X-Plat, for bidirectional conversion of microarray and sequencing data for both expression and methylation assays. X-Plat builds per-gene predictive models by fitting second-degree polynomial regressions to paired array–sequencing values and evaluates these models using 10-fold cross-validated RMSE. It derives cross-platform conversion rules from paired mixed-source microarray–sequencing datasets spanning multiple conditions, sample sources and organisms, and multiple array and sequencing platforms.

## MATERIAL AND METHODS

### Collecting data

We selected datasets from NCBI Gene Expression Omnibus (https://www.ncbi.nlm.nih.gov/geo) for rat, *Arabidopsis*, and human expression studies and for human methylation studies. Datasets with at least 50 paired samples, where the same sample was assayed using both the same microarray platform and the same high-throughput sequencing platform, were chosen to build the model. The sample and gene numbers, including platform details are summarized in Table 1, and sample-level details are provided in Supplementary Table S1.

**Table 1:** Description of paired microarray–sequencing data used to build and test X-Plat.

| Assay | Organism | Array platform | Seq platform | No. of paired samples | No. of probes/regions | No. of genes |
| --- | --- | --- | --- | --- | --- | --- |
| Expression | Rat<br>(*Rattus norvegicus*) | Affymetrix Rat Genome 230 2.0 Array | Illumina HiScanSQ | 148 | 21 887 | 11 012 |
|  |  |  | Illumina HiSeq 2000 |  |  |  |
|  |  |  | Illumina HiSeq 500 |  |  |  |
|  | *Arabidopsis*<br>(*Arabidopsis thaliana*) | Affymetrix Arabidopsis ATH1 Genome Array | Illumina NextSeq 500 | 111 | 22 812 | 8 785 |
|  | Human<br>(*Homo sapiens*) | Affymetrix HG-U133 Plus 2.0 Array | Illumina HiSeq 2000 | 62 | 54 675 | 14 323 |
| Methylation | Human<br>(*Homo sapiens*) | Illumina HumanMethylation450 BeadChip Array | Illumina HiSeq 2500 | 99 | 119 535 | 19 607 |

### Data pre-processing

The pipelines used for pre-processing the expression and methylation data, including quality assessment using the Galaxy framework (stable release v22.05), are illustrated in Figure 1 (18).

**Figure 1:**
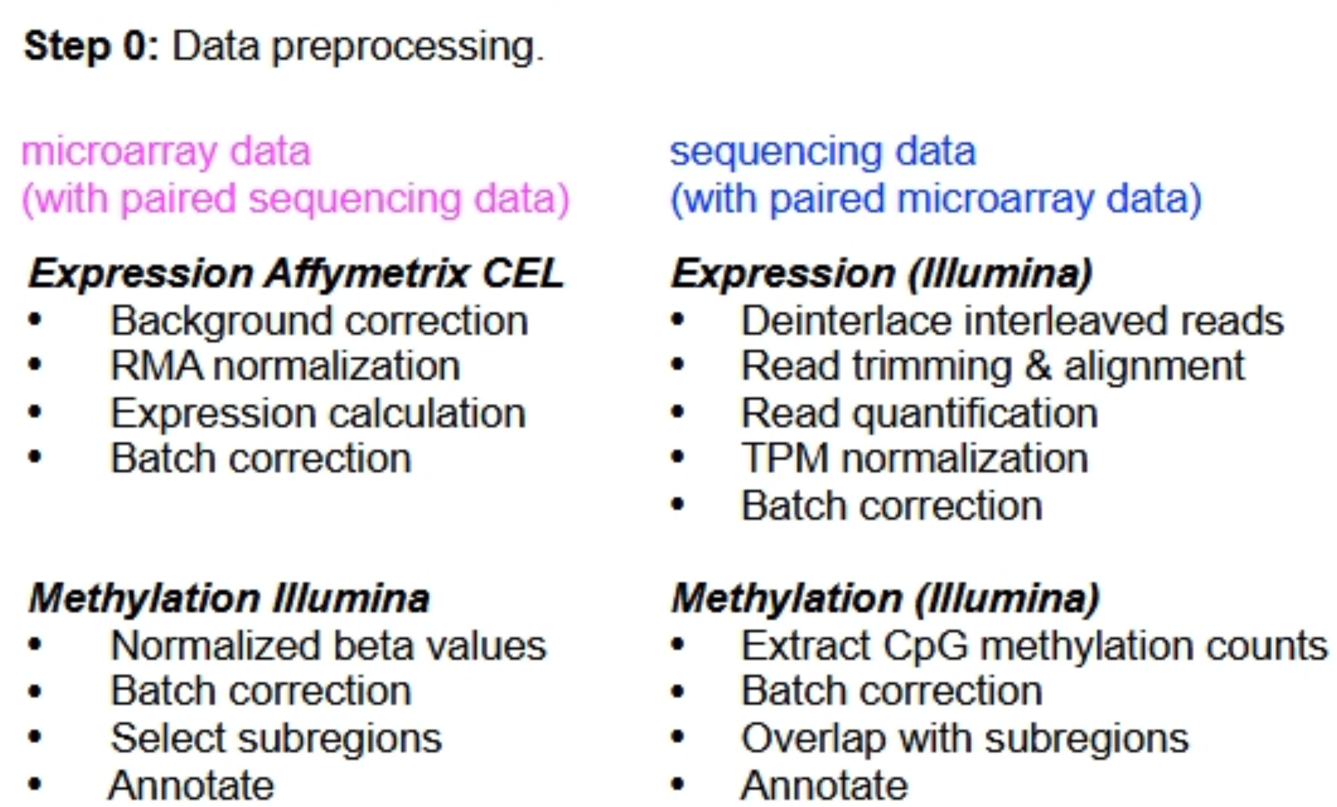
Pre-processing of expression and methylation data. The schematic summarizes platform-specific preprocessing steps for paired microarray and sequencing assays, including normalization, batch correction, and region-level annotation for both expression and methylation data.

#### Expression data

We analyzed the Affymetrix microarray expression data (.CEL) using the R/Bioconductor package affy (19). We first normalized the data using the robust multi-array average (RMA) procedure to correct for systematic, non-biological variation in three steps: (a) background correction, (b) adjustment for hybridization effects unrelated to the probe–target interactions, and (c) normalization. This was followed by summarization, in which multiple probe intensities within a probe set were combined to yield a single value per gene representing the RNA transcript expression level. At this stage, we corrected for batch effects using ComBat (7) and then performed probe-to-gene annotation.

We implemented the pre-processing pipeline for RNA-seq data on the Galaxy server (18). First, we extracted reads using the NCBI SRA Toolkit fastq-dump utility (v2.11.0) as single interleaved FASTQ files. We then used the Galaxy FASTQ de-interlacer tool (v1.1.5) to split interleaved reads into two FASTQ files for the paired reads. Quality control was performed using FastQC (v0.73) (20), followed by adaptor and Illumina-specific sequence trimming using Trimmomatic v0.38 (21) using the ILLUMINACLIP and SLIDINGWINDOW parameters. Next, we aligned reads to the reference genomes (mRatBN7.2 for rat, TAIR10 for *Arabidopsis*, and T2T-CHM13v2.0 for human) using HISAT2 (v2.2.1) (22). To quantify the number of reads mapped to each gene, we used featureCounts (v2.0.1) on the aligned BAM files together with the corresponding gene annotation for each genome. We then adjusted the counts for batch effects using the R package Combat-Seq (23) to eliminate batch-related technical variation, before normalizing to transcripts per kilobase million (TPM) using the R package DGEobj.utils (v1.0.6) (24).

#### Methylation data

We downloaded Illumina HumanMethylation450 (450K) microarray methylation data from NCBI GEO as normalized β-values and batch-corrected these using ComBat (7). We downloaded all bisulfite-seq methylation data as CpG-level methylation counts from NCBI GEO and batch-corrected these using ComBat as well. The raw files (.IDAT for microarrays and .fastq for sequencing) were not available for all methylation studies; therefore, full preprocessing from raw intensities or reads was not possible.

### X-Plat algorithm workflow

The workflow of the X-Plat algorithm is illustrated in Figure 2. We prepared data for each platform as matrices, where the rows represented samples and columns represented genes. We ensured a one-to-one correspondence between the microarray (array) and sequencing (seq) matrices in terms of both the order and exact identity of samples and genes before proceeding with the analysis. For brevity, we refer to these as “array” and “seq” and denote the two prediction directions as array→seq and seq→array. For each gene, we considered all available paired array–seq samples and worked with their log-transformed values. The array and sequencing data were transformed using Python’s NumPy function log1p (natural logarithm of one plus the input value) element-wise to handle small values (25).

**Figure 2:**
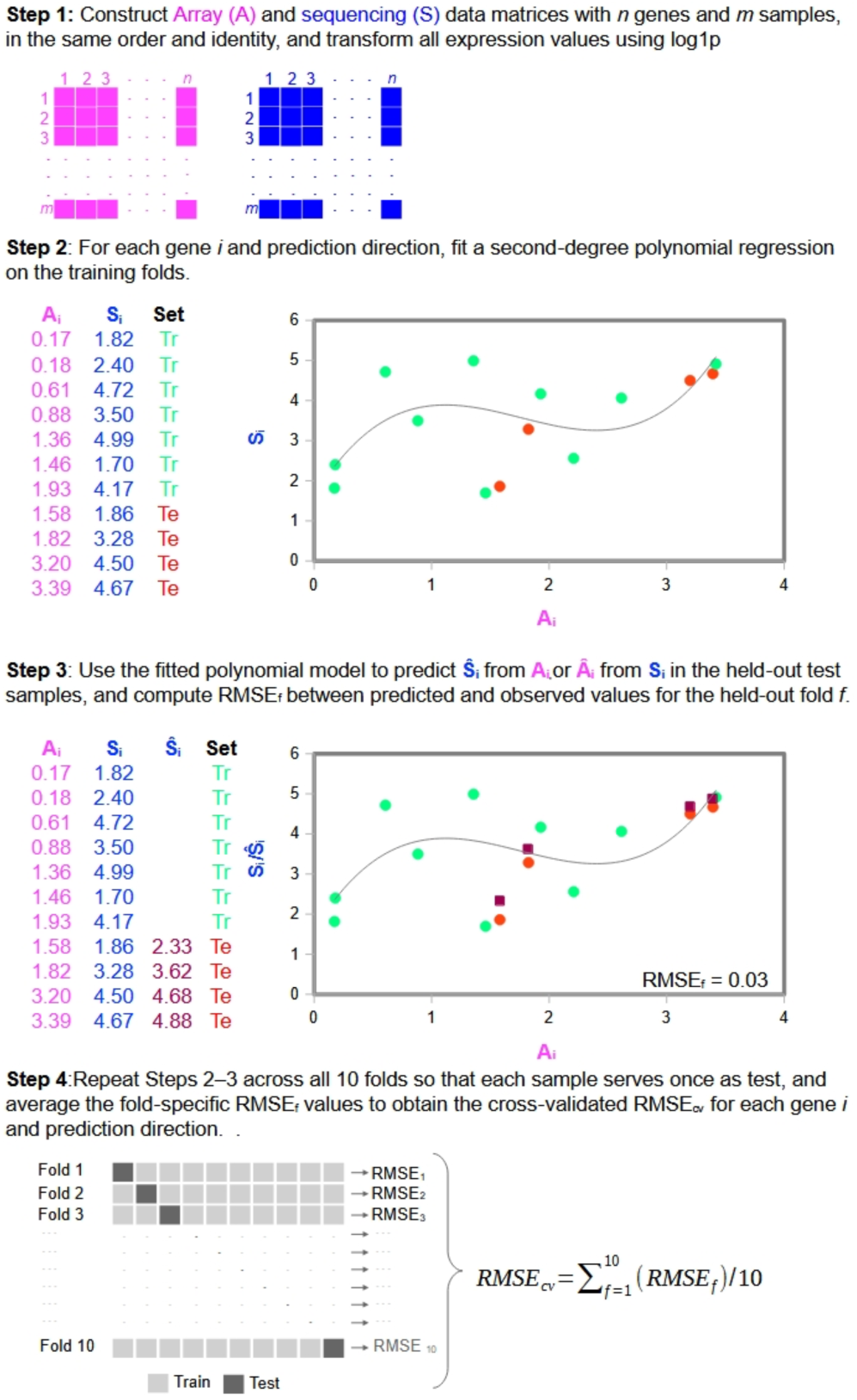
X-Plat algorithm workflow. The schematic illustrates the construction of paired array (A) and sequencing (S) matrices with matched genes and samples, log1p transformation, fitting per-gene second-degree polynomial regression models on training samples for each prediction direction, using the fitted models to predict expression values in the held out test samples for each fold, and computation of cross-validated RMSE across folds for both array→seq and seq→array prediction directions.

For array-to-sequencing (array → seq) conversion, a second-degree polynomial regression model is fitted per gene g to the paired log-transformed values (*array_ig_*, *seq_ig_*), and the polynomial coefficients are estimated for each gene. The fitted model is then used to predict log-transformed sequencing values *seq_jg_* for held-out samples j from their corresponding log-transformed array values, *array_jg_*. The prediction errors e*_jg_* between predicted and observed sequencing values are computed for each held-out sample, and a root mean square error (RMSE) is calculated over the held-out set for that gene as

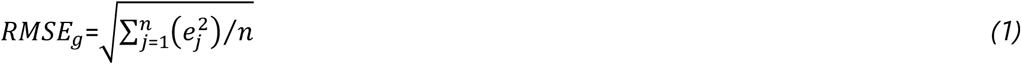

where *n* is the number of held-out samples for gene *g*.

For sequencing-to-array (seq → array) conversion, the roles of the array and seq axes are reversed in the above procedure, and a second-degree polynomial is fitted to predict log(1 + array) from log(1 + seq).

We evaluated prediction performance using 10-fold cross-validation, performed independently for each prediction direction across genes. All paired samples were randomly partitioned into 10 approximately equal folds with shuffling and a fixed random seed. In each iteration, a polynomial regression model was trained on 9 folds and evaluated on the remaining fold, and we computed the RMSE between predicted and observed log-transformed values on the held-out fold. This process was repeated so that each fold served once as the test set. For each gene and prediction direction, we thus obtained 10 RMSE values {*RMSE_1_*,…,*RMSE_10_*} and computed their mean as

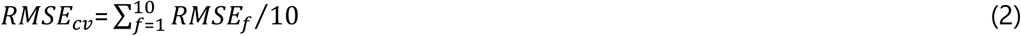

where *RMSE_cv_* denotes the cross-validated RMSE and *f* indexes the fold.

These mean 10-fold RMSE values were used to summarize gene-wise performance distributions and for all figures that report X-Plat RMSE alone. For direct comparisons between X-Plat and other cross-platform normalization tools, we used the RMSE from the first cross-validation fold (i.e., the RMSE on the test set of that fold) so that all methods were evaluated on the same held-out samples.

To assess robustness to training sample size, we performed down-sampling analyses on the rat expression dataset, randomly reducing the original training set to 90%, 70%, 50%, or 30% of its size while keeping the same held-out test set and computing cross-validated RMSE as described above.

### Comparison with other tools

We compared the performance of X-Plat to three other tools, Training Distribution Matching (TDM; v0.9.9; 8), Shambhala as implemented in HARMONY, and Shambhala-2 as implemented in HARMONY2 (26,27), using the held-out test set samples from the first cross-validation fold so that all methods were evaluated on the same samples. As in X-Plat, we applied the log1p transform to both microarray and sequencing data for the test samples using Python’s NumPy implementation prior to running the tools.

TDM is an R package that adjusts the distribution of RNA-seq data to make it more compatible with models trained on microarray data while preserving between-sample relationships, thereby improving recognition of features learned from microarrays. It uses the tdm_transform function on the matrix of RNA-seq values (set as the target) and the microarray expression values (set as the reference) to produce transformed RNA-seq values, which we used for RMSE estimation.

Shambhala is a cross-platform data harmonization tool that compares multiple gene expression datasets by transforming them into a common, platform-independent representation. It harmonizes a target dataset to a reference definitive dataset *Q* using an auxiliary calibration dataset and makes the target more comparable to the reference platform (26). Shambhala is implemented in the R package HARMONY, which is available at https://github.com/oncobox-admin/harmony. We used the harmony function with paired microarray and sequencing data from the test set and extracted harmonized sequencing values for RMSE estimation.

Shambhala-2 (HARMONY2) is an updated harmonizer of gene expression profiles obtained from both microarray hybridization and next-generation sequencing. It converts each profile independently to a predefined “definitive” shape defined by a reference dataset Q, using an auxiliary calibration dataset P to make the conversion more robust (27). Briefly, each profile to be harmonized is first quantile-normalized to P, then normalized using the CuBlock method, and finally rescaled so that for each gene the mean and standard deviation match those of the definitive dataset Q. In our analyses, we used the HARMONY2 R package (https://github.com/oncobox-admin/harmony2) with the Shambhala-2 implementation. For sequencing-to-microarray conversion, we supplied log1p-transformed test RNA-seq profiles as inputs, with the log1p-transformed training microarray data serving as both the calibration dataset P and the definitive dataset Q. Conversely, for microarray-to-sequencing conversion, we supplied log1p-transformed test microarray profiles as inputs, with the log1p-transformed training RNA-seq data serving as both P and Q. The resulting harmonized profiles were used for RMSE and prediction-error comparisons alongside X-Plat, TDM, and HARMONY.

In addition to RMSE, we compared sample-wise prediction errors across X-Plat, TDM, HARMONY and HARMONY2. For all three organisms (rat, *Arabidopsis* and human), and for each gene, we computed the difference (Δ) in prediction errors (*E*) between tools for each test sample, separately for array (a) and sequencing (s) expression values. Specifically, for each test sample we calculated *E_a_*(X-Plat) − *E_a_*(TDM), *E_a_*(X-Plat) − *E_a_*(HARMONY), *E_a_*(X-Plat) − *E_a_*(HARMONY2), *E_s_*(X-Plat) − *E_s_*(TDM), *E_s_*(X-Plat) − *E_s_*(HARMONY) and *E_s_*(X-Plat) − *E_s_*(HARMONY2) which we refer to as Δ*E_a_*(XT), Δ*E_a_*(XH), Δ*E_a_*(XH2), Δ*E_s_*(XT), Δ*E_s_*(XH) and Δ*E_s_*(XH2), respectively. A negative Δ*E_a_*(XT) for a given sample–gene combination indicates that TDM has a smaller prediction error than X-Plat, whereas a positive value indicates the reverse; analogous interpretations apply for Δ*E_a_*(XH), Δ*E_a_*(XH2), Δ*E_s_*(XT), Δ*E_s_*(XH) and Δ*E_s_(*XH2).

For each gene, we recorded the maximum negative and maximum positive Δ*E* values across samples (max Δ) for each of the Δ*E_a_*(XT), Δ*E_a_*(XH), Δ*E_a_*(XH2), Δ*E_s_*(XT), Δ*E_s_*(XH) and Δ*E_s_*(XH2) categories. In each category, we then selected the five genes with the largest positive and the five with the largest negative max Δ values and examined their expression profiles in detail.

## RESULTS

### Performance trends of X-Plat across platforms, assays, and organisms

The performance of X-Plat in predicting sequencing values from microarray values and vice versa was primarily measured using cross-validated RMSE (see Materials and Methods). For ease of discussion, we refer to the RMSE for predicted microarray values as array RMSE, and the RMSE for predicted sequencing values as seq RMSE.

#### Expression data

X-Plat achieved array RMSE ranges of 0–0.25, 0.01–0.18, and 0.01–0.35, and seq RMSE ranges of 0–1.21, 0.04–3.51, and 0.04–1.87 for expression data across 11 012, 8 785, and 14 323 genes from rat, *Arabidopsis* and human, respectively (Figure 3). The corresponding median array RMSE values were 0.03, 0.06, and 0.04, and the median seq RMSE values were 0.17, 1.25, and 0.34, respectively. These ranges correspond to relatively small errors on the log scale compared with the overall expression dynamic range, indicating that most genes are predicted with modest absolute deviation. Overall, the RMSE range was consistently smaller for the microarray platform than for the sequencing platform.

**Figure 3:**
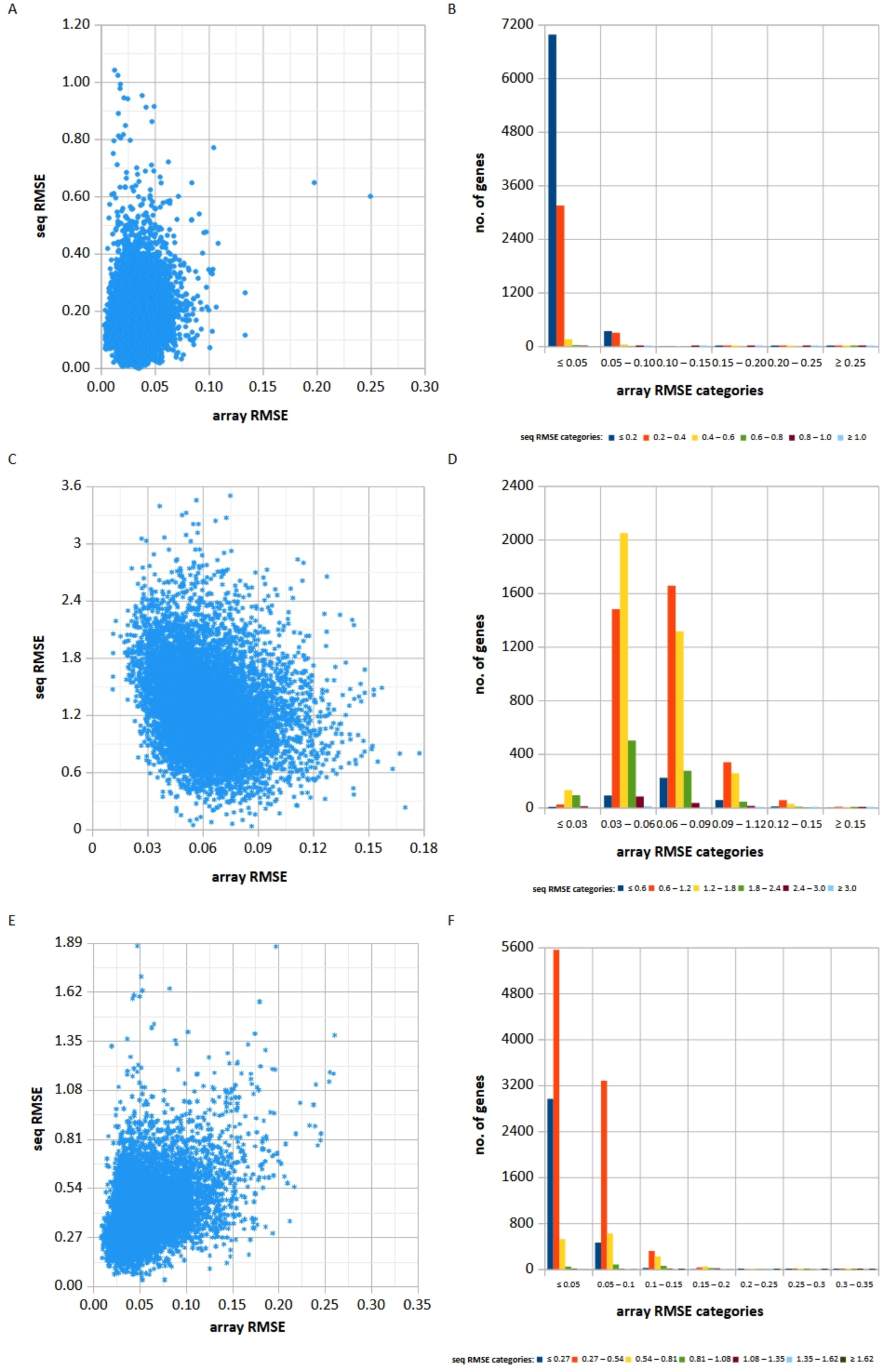
Root mean square errors (RMSE) for X-Plat-predicted microarray and RNA-seq expression values (array RMSE and seq RMSE) in rat (A–B), *Arabidopsis* (C–D), and human (E–F). Scatterplots of array RMSE vs seq RMSE (A, C, E) and histograms of gene counts within predefined array RMSE and seq RMSE categories (B, D, F) are shown.

X-Plat’s prediction trends varied across organisms. For rat (Figure 3A–B), array RMSE was ≤0.05 for most genes (10 576/11 012) and 0.05–0.1 for the remainder. Among these 10 576 genes, seq RMSE was ≤0.2 for 7 354 genes. Only one gene had a high array RMSE of 0.25, and 18 genes had relatively high seq RMSE values ≥0.8.

For *Arabidopsis* (Figure 3C–D), array RMSE was in an intermediate-low range of 0.03–0.09 for most genes (7 645/8 785). Within this subset, seq RMSE was 0.6–1.2 for 3 235 genes and 1.2–1.8 for 3 256 genes; 18 genes had higher array RMSE (≥0.15) and 16 genes had higher seq RMSE (≥0.24).

For human (Figure 3E–F), array RMSE was ≤0.05 and 0.05–0.10 for most genes (9 100 and 5 223 of 14 323 genes, respectively). Among these 9 100 and 5 223 genes, seq RMSE was 0.27–0.54 for 5 557 and 3 628 genes, respectively, and ≤0.27 for 2 963 and 487 genes, respectively. Only one gene, *SCGB2A2*, had a higher array RMSE (0.35), and 6 genes had higher seq RMSE (>1.62).

#### Methylation data

For the human methylation data (Figure 4A), there were 127 031 CpG regions after annotation, pairing between microarray and sequencing data, filtering out regions for which more than 3% of samples had zero values, and restricting to chosen sub-regions for which X-Plat was applied. Of these, 15 895 regions were within TSS200 of genes, 27 401 within TSS1500, 71 837 within the 5′ UTR, 21 932 within the 3′ UTR, and 43 064 within the first exon. Overall, array RMSE values were in the low range (≤0.1) for most (120 707 out of 127 031) filtered CpG regions. Among these, 120 681 regions had low seq RMSE (≤0.2), 23 regions had seq RMSE between 0.2 and 0.4, and only two regions had seq RMSE ≥0.4: cg09872233 in the first exon of *ALOX15* (array RMSE 0.015, seq RMSE 0.61) and cg24043887 in the TSS200 region of *TMEM80* (array RMSE 0.014, seq RMSE 1.12) (Figure 4B). The highest array RMSE (0.49) was observed for cg03884082 in the 5′ UTR of *NBL1* (Figure 4B), followed by cg13449778 (array RMSE 0.48) in the first exon of *FAM163A* (Figure 4B).

**Figure 4:**
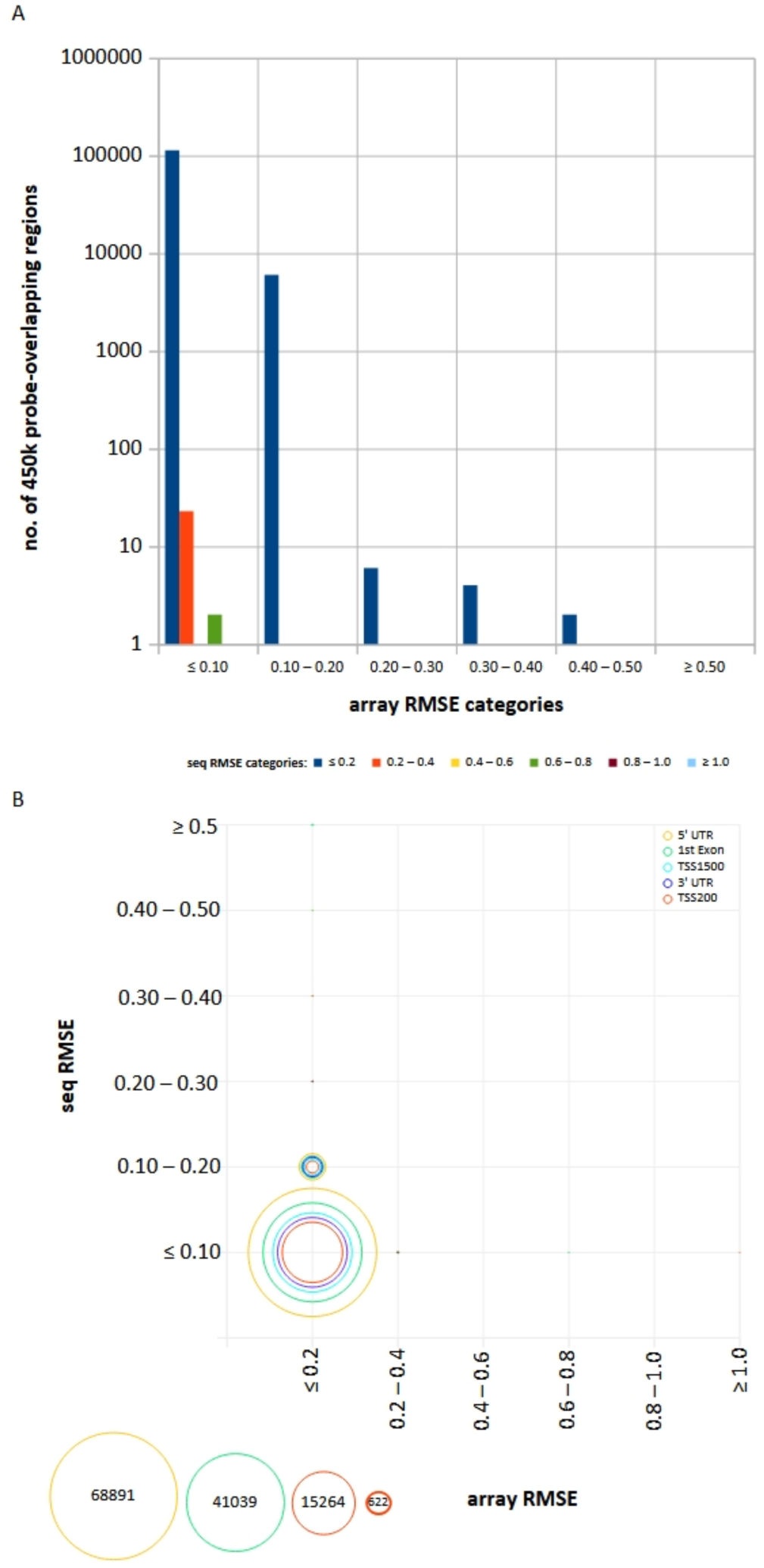
Root mean square error (RMSE) values for X-Plat-predicted methylation microarray and sequencing values (array RMSE and seq RMSE) in human breast cancer samples for probes overlapping various sub-regions (TSS200, TSS1500, 5′ UTR, 3′ UTR, and first exon) (A), and for each sub-region individually (B). Histograms show the number of overlapping regions within predefined array RMSE and seq RMSE categories; bubble sizes represent individual sub-regions within each region category.

Considering individual sub-regions, the dominant pattern of most CpGs falling within the combined ≤0.2 array RMSE and ≤0.2 seq RMSE range was preserved across TSS200, TSS1500, 5′ UTR, 3′ UTR, and first exon (Figure 4B). The 3′ UTR was a slight exception, with a modest decrease (to 92.71%) in the proportion of CpGs within the 0–0.1 array RMSE and 0–0.2 seq RMSE combined range, accompanied by an increase (to 7.26%) in the 0.1–0.2 array RMSE and 0–0.2 seq RMSE combined range (Figure 4B).

### Factors affecting X-Plat RMSE for expression data

We explored whether factors such as expression value range and transcript length could explain variability in X-Plat RMSE across genes, and observed that the expression data range correlated directly with RMSE for both array and sequencing transformations.

#### Data Range

We observed a strong positive correlation between array RMSE and the range of array expression values, and between seq RMSE and the range of sequencing expression values (Figure 5). The Pearson correlation coefficients (*r*) were 0.51 (array) and 0.50 (seq) for rat, 0.91 (array) and 0.91 (seq) for *Arabidopsis*, and 0.92 (array) and 0.85 (seq) for human, indicating a stronger trend in *Arabidopsis* and human than in rat.

**Figure 5:**
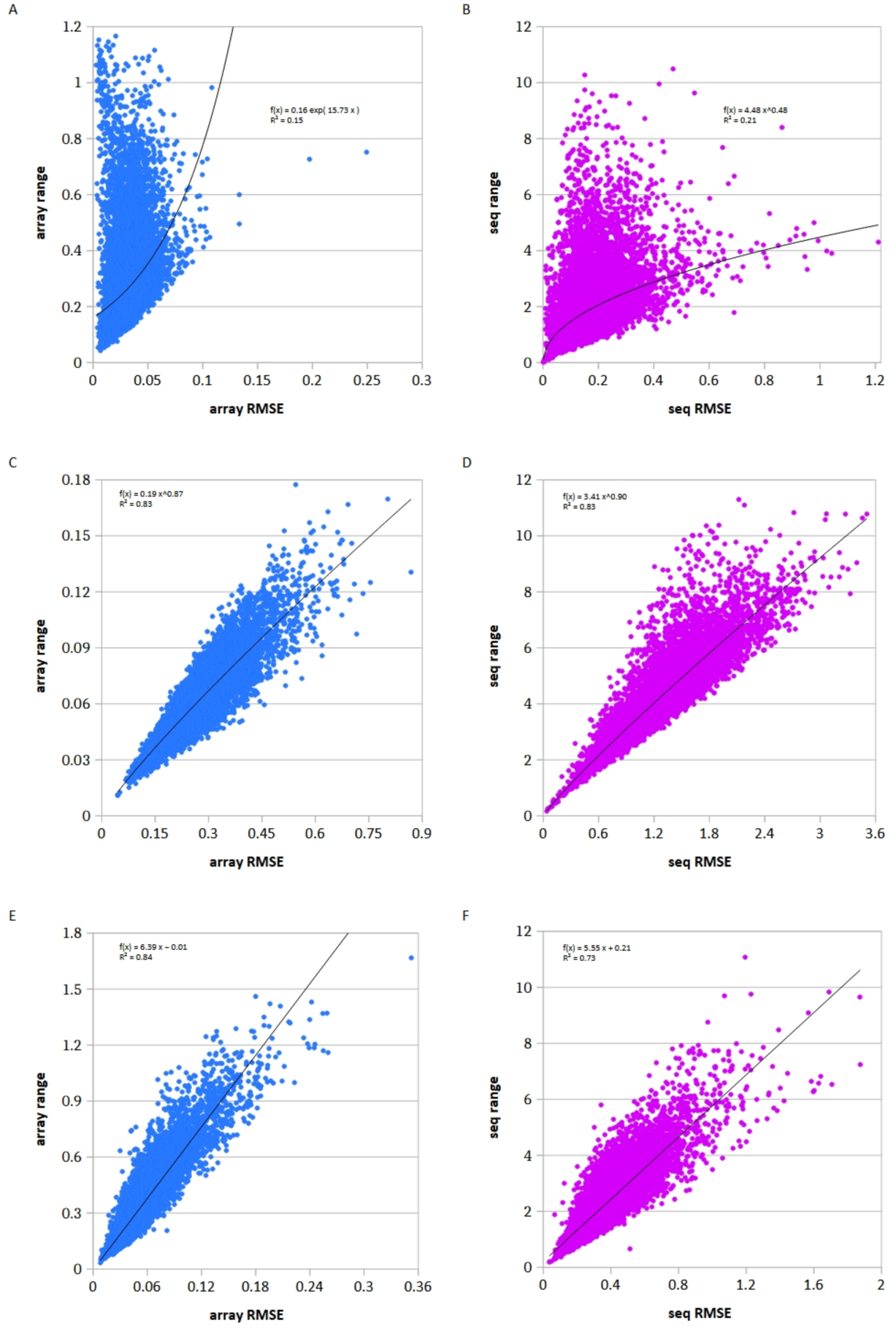
Root mean square errors (RMSE) for X-Plat-predicted microarray and RNA-seq expression values (array RMSE and seq RMSE) in rat (A–B), *Arabidopsis* (C–D), and human (E–F). Scatterplots of array RMSE vs array range (A, C, E) and seq RMSE vs seq range (B, D, F), where “range” denotes the difference between maximum and minimum expression values, are shown.

#### Relationship with transcript length

The RMSE values bear no direct correlation with transcript length for either array or sequencing data in any of the organisms studied (Supplementary Figure S1).

### Comparing X-Plat with other transformation tools

We applied TDM, HARMONY and HARMONY2 to the expression datasets from all three organisms and compared the resulting RMSE values on the held-out test samples from the first cross-validation fold with those obtained using X-Plat (Figure 6).

**Figure 6:**
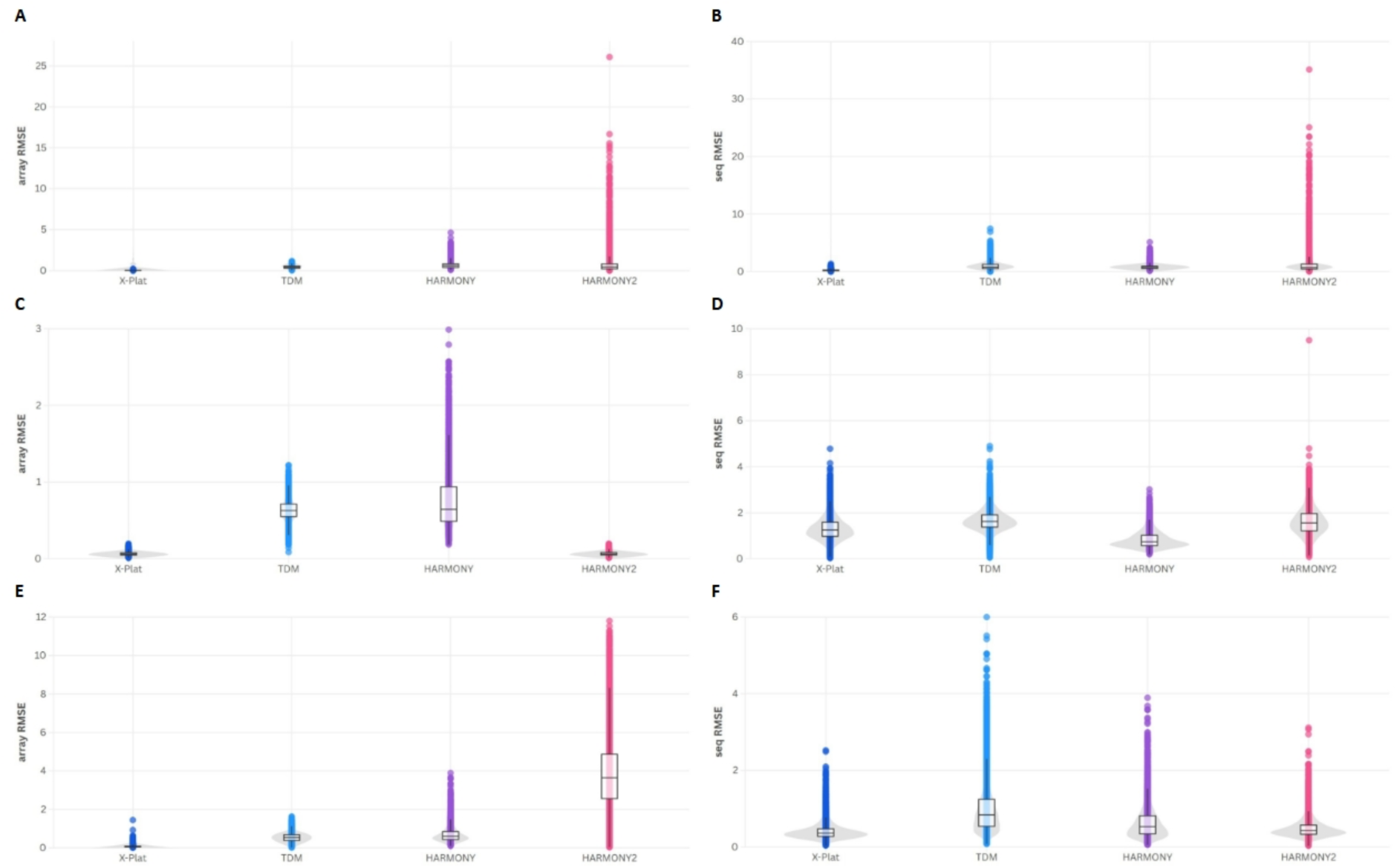
Root mean square error (RMSE) values for X-Plat, TDM, HARMONY, and HARMONY2 for array RMSE (A, C, E) and seq RMSE (B, D, F) across rat, *Arabidopsis*, and human.

#### Comparison using RMSE

In terms of mean RMSE per gene, X-Plat outperformed TDM, HARMONY and HARMONY2 for the vast majority of genes across organisms and directions. Specifically, for the sequencing-to-array transformation, X-Plat achieved lower average RMSE than all three tools for 100%, 100%, and 99.92% of genes in rat, *Arabidopsis*, and human, respectively, and for the array-to-sequencing transformation, for 99.65%, 82.53%, and 95.80% of genes in these organisms. The array RMSE spread for X-Plat was ≤0.5 across all three organisms. TDM showed a broader array RMSE spread (≤1.5), HARMONY had an even broader spread, with array RMSE extending beyond 2 in all three organisms; and HARMONY2 had the largest array RMSE spread in rat and human, but a tighter array RMSE spread comparable to X-Plat in Arabidopsis (Figure 6A, C, E). In terms of sequencing RMSE, X-Plat values were ≤2 in rat and human and substantially lower than those of TDM, HARMONY and HARMONY2, whereas in *Arabidopsis* they were generally ≥3; comparable to TDM and HARMONY, while still outperforming HARMONY2 (Figure 6B, D, F), with X-Plat performing best overall, followed by HARMONY or HARMONY2 and then TDM.

#### Comparison using per-sample prediction error

For rat array predictions, X-Plat had the lowest prediction error in a substantial fraction of samples for nearly all genes (Figure 7A). X-Plat was the best-performing tool in at least 80% of samples for 673 of 11 012 genes, including 21 genes for which it had the lowest error in all samples, and in a further 4,676 and 5,228 genes it was best in 60–80% and 40–60% of samples, respectively. For every gene, X-Plat was the best tool in at least 21% of the samples, with *Mrpl13* and *Psmd11* representing the lower end of this range, where X-Plat was optimal in fewer than 25% of samples. In contrast, TDM had the lowest array prediction error in more than 20% of samples for only 30 genes (with *Rictor* reaching 35.13%), and HARMONY did so for 15 genes, with *Bcat2*, *Sgf29*, and *Pde8a* exceeding 25%. HARMONY2 performed particularly well for a small subset of genes, achieving the lowest array prediction error for *Lamb3*, *Sh3glb2*, and *Stx4* in at least 70% of samples, but for 931 genes it was best in at most 20% of samples.

**Figure 7:**
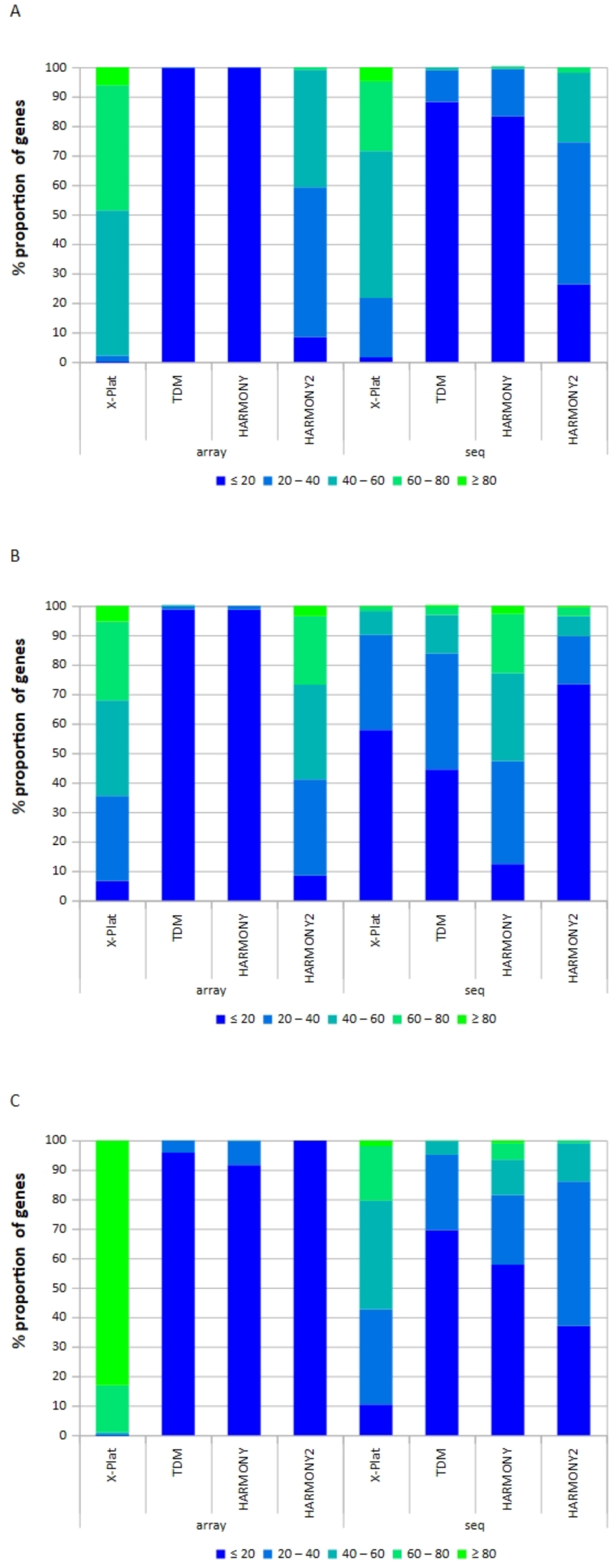
Comparison of results obtained using X-Plat, TDM, HARMONY, and HARMONY2. Prediction errors and sample frequencies in five threshold categories (≤20%, 20–40%, 40–60%, 60–80%, ≥80%) are shown as histograms of the proportion of genes, separately for array and seq predictions, for rat (A), *Arabidopsis* (B), and human (C).

For the array → seq transformation in rat, X-Plat had the lowest prediction error in all samples for 21 genes, namely, *Alb*, *Aldob*, *Ano9*, *Azgp1*, *Bhmt2*, *C3*, *Cd6*, *Cfi*, *Cpb2*, *Hpd*, *Itih1*, *LOC100911093*, *LOC120093688*, *RGD1561916*, *Rpl19*, *Serpina1*, *Serpinc1*, *Tat*, *Tdo2*, *Thsd7a* and *Ttn*, in ≥80% of samples for 510 more genes and in <20% of samples for 171 genes, including *Fcho1*, *Kcnab2*, and *Unc80*, for which X-Plat had the lowest seq prediction error in only 2.7% of samples (Figure 7B). Among the remaining genes, X-Plat had the lowest seq prediction error in 60–80% of samples for 2 601 genes (23.62%), in 40–60% of samples for 5 487 genes, and in 20–40% of samples for 2 222 genes. In contrast, TDM, HARMONY, and HARMONY2 had the lowest seq prediction error in ≥80 % of the samples for 6, 0, and 5 genes, respectively and in ≤20 % of the samples for 9 720, 9 170, and 2 900 genes, respectively.

For Arabidopsis, X-Plat had the lowest prediction error for the seq → array transformation in ≥80% of samples for 473 genes, including in all samples for 10 genes, namely *at2g40090*, *at3g01290*, *at3g48500*, *at3g52170*, *at3g56940*, *at3g61410*, *at4g32070*, *at5g15390*, *at5g17640*, and *at5g53900* (Figure 7C). For the remaining genes, X-Plat still had the lowest array prediction error in 60–80% of samples for 26.68% of genes, 40–60% of samples for 32.48% of genes, 20–40% of samples for 28.88% of genes, and ≤20% of samples for the remaining 6.6% of genes. TDM and HARMONY had the lowest array prediction errors predominantly (98.83% and 98.68% of genes, respectively) only in the ≤20% sample-frequency range, with the remainder in 20–40% of samples. HARMONY2 had the lowest array prediction error in all samples for two genes, namely *at3g55610* and *at5g54540*, in ≥80% of samples for 309 genes, and in ≤20% of samples for 745 genes.

For the array → seq transformation in *Arabidopsis*, X-Plat had the lowest prediction error in ≥80% of samples for 21 genes (Figure 7D). Among the remaining genes, X-Plat had the lowest seq prediction error in 60–80% of samples for 138 genes (1.57%), in 40–60% of samples for 705 genes, in 20–40% of samples for 2 839 genes, and in ≤20 % of samples for the remaining 57.85 % of genes. TDM performed with low prediction error for 83.85% of genes in ≤40% of samples. HARMONY had the lowest prediction error for comparatively more genes in higher sample frequency ranges (29.78%, 20.14%, and 2.73% of genes in the 40–60%, 60–80%, and ≥80% sample frequency ranges, respectively). HARMONY2, on the other hand, had the lowest seq prediction error in ≤20% of samples for 73.43% of genes.

For human array predictions, X-Plat overwhelmingly had the lowest array prediction error across samples (Figure 7E). X-Plat was the best-performing tool in ≥80% of samples for 83.09% of genes and in 60–80% of samples for a further 15.94% of genes, with only 0.96% and 0.01% of genes falling in the 40–60% and 20–40% sample-frequency ranges, respectively, and none in the ≤20% range. In contrast, TDM and HARMONY almost never dominated at high sample frequencies and instead had the lowest array prediction error mainly at very low sample frequencies: 95.85% and 91.65% of genes, respectively, were best for these tools in ≤20% of samples, with only small fractions (4.12% and 8.20%) in the 20–40% range and virtually none above 40%. HARMONY2 did not achieve the lowest array prediction error in more than 20% of samples for any gene, with all genes falling in the ≤20% sample-frequency range.

For human seq predictions, X-Plat most often had the lowest seq prediction error at intermediate sample-frequency ranges (Figure 7F). X-Plat was the best-performing tool in 40–60% of samples for 36.77% of genes and in 60–80% of samples for a further 18.54% of genes, with only 10.38% of genes falling in the ≤20% sample-frequency range. In contrast, TDM and HARMONY had the lowest seq prediction error predominantly at low sample frequencies, with 69.59% and 57.84% of genes, respectively, showing TDM or HARMONY as best in ≤20% of samples and only 4.67% and 12.02% of genes in the 40–60% range; genes for which these tools were best in ≥80% of samples were rare or absent (0% for TDM and 0.88% for HARMONY). HARMONY2 showed a mixed pattern, with the lowest seq prediction error in ≤20% of samples for 37.08% of genes but in 20–40% and 40–60% of samples for 49.01% and 13.06% of genes, respectively, and only 0.03% of genes in the ≥80% range.

#### Factors affecting differences in per-sample prediction error

We next explored factors that could explain differences in performance between the four tools, i.e. why X-Plat had the lowest predicted seq error for most genes at high sample-frequency ranges, but not for all genes, and to identify cases in which TDM, HARMONY, or HARMONY2 instead had the lowest prediction error.

For each organism and each comparison category (Δ*E*_a_(XT), Δ*E*_a_(XH), Δ*E*_a_(XH2), Δ*E_s_*(XT), Δ*E_s_*(XH), and Δ*E_s_*(XH2)), we selected the top five genes with the largest positive Δ values and the top five with the largest negative Δ values (10 genes per category) for detailed inspection. These genes represent cases where X-Plat or the comparator tool most strongly outperformed the other, allowing us to investigate expression-related factors associated with extreme differences in prediction error.

For the seq → array direction, array expression values showed a much tighter spread with higher median expression (2.6–2.7 in rat, 2.5–2.62 in *Arabidopsis*, and 2.55–2.6 in human; Figure 8A, C, E) for genes with the top five maximum positive Δ array values (Δ*E*_a_(XT), where X-Plat outperformed TDM in at least one sample) than for genes with the top five minimum negative Δ array values, where TDM outperformed X-Plat in at least one sample (1.7–2.5 in rat, 1.5–2.2 in *Arabidopsis*, and 1–2.2 in human; Figure 8A, C, E). This pattern suggests better performance of X-Plat for more highly expressed genes across organisms. A similar trend was observed when comparing X-Plat with HARMONY for predicted array expression in rat and *Arabidopsis*, but to a lesser extent in human, and when comparing X-Plat with HARMONY2 for predicted array expression in rat, but not in *Arabidopsis* or human.

**Figure 8:**
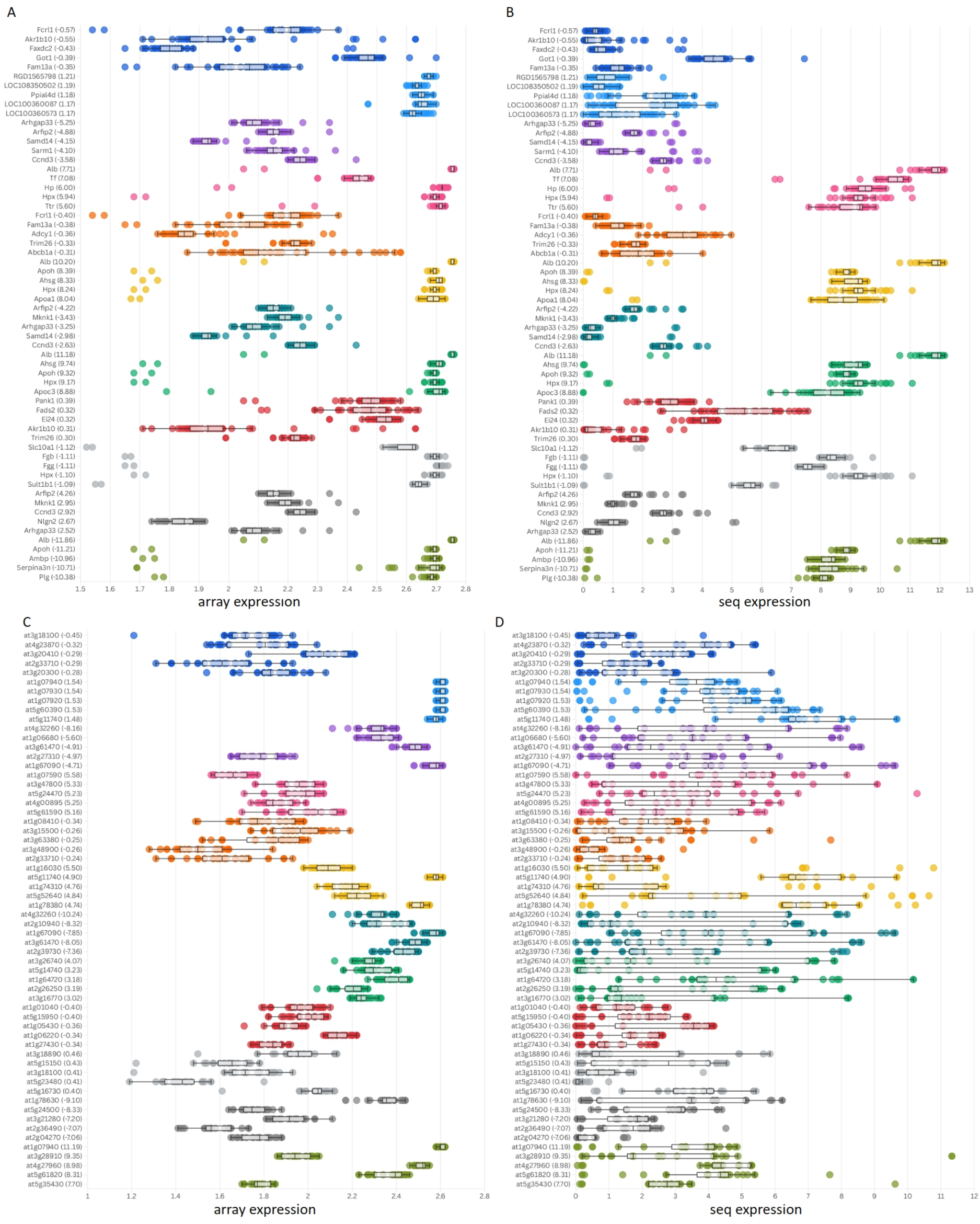
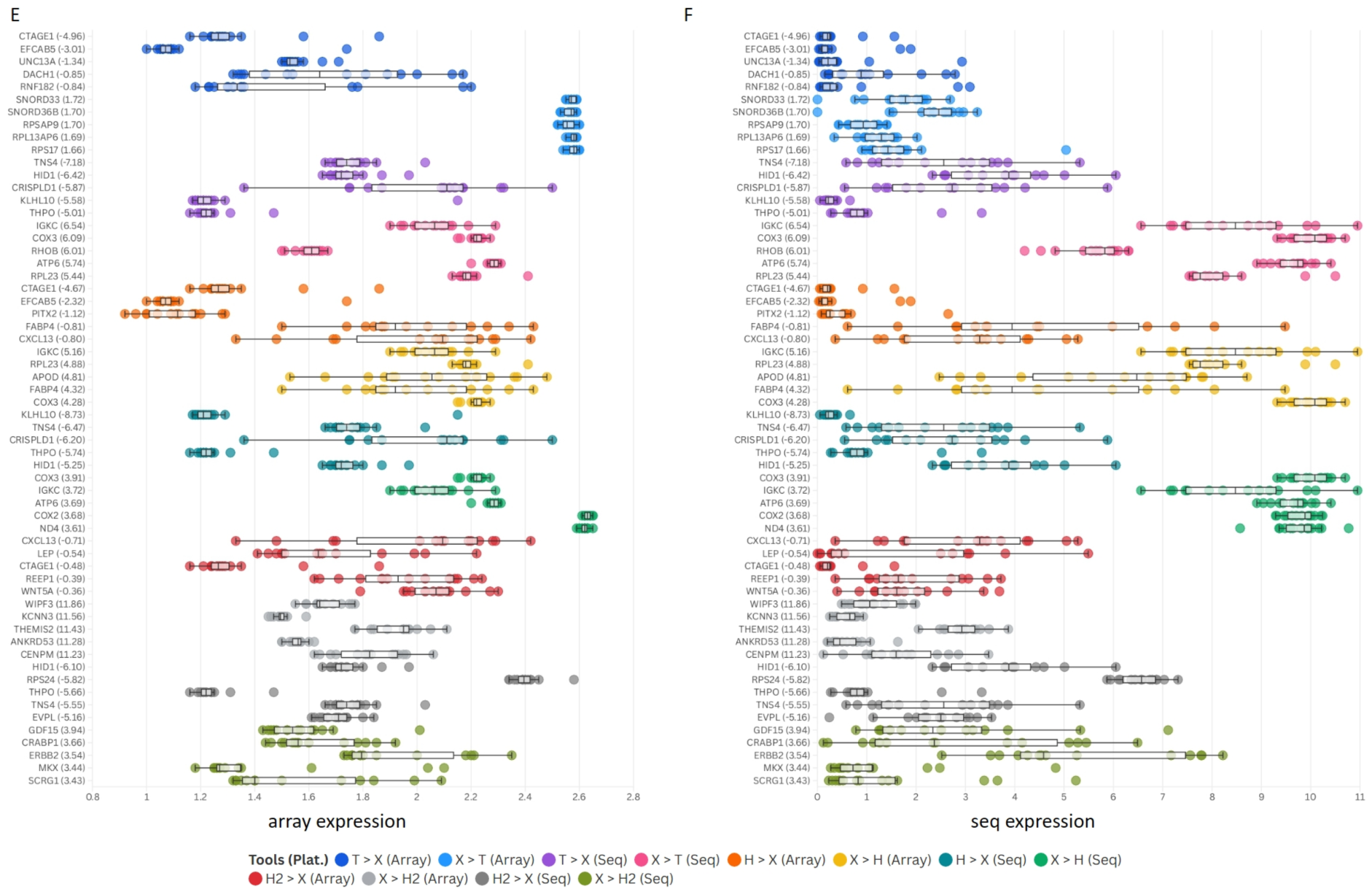
Array (A, C, E) and sequencing (B, D, F) expression values for the genes with five largest positive and negative Δ values in each of Δ*E_a_*(XT), Δ*E_a_*(XH), Δ*E_a_*(XH2), Δ*E_s_*(XT), Δ*E_s_*(XH), and Δ*E_s_*(XH2) categories for rat (A–B), *Arabidopsis* (C–D), and human (E–F) datasets (see Methods for details on estimation of Δ*E*). T > X indicates cases where TDM predictions outperform those of X-Plat; other labels (X > T, H > X, X > H, H2 > X and X > H2) are interpreted analogously for the corresponding tool pairs. The y-axes list gene names with the corresponding Δ value in parentheses.

For the array → seq direction, sequencing expression values also showed a relatively higher spread (∼8.5–12 in rat and ∼5.5–∼10.3 in human; Figure 8B, F) for genes with the top five maximum positive Δ seq values (Δ*E*_s_(XT), where X-Plat outperformed TDM in at least one sample) than for genes with the top five minimum negative Δ seq values, where TDM outperformed X-Plat (0–∼3 in rat and ∼0–4.5 in human; Figure 8B, F). This again suggests that X-Plat performs better for genes with higher sequencing expression. In *Arabidopsis*, this trend was not seen with Δ*E*_s_(XT) versus sequencing expression levels; instead, an inverse relationship was observed between Δ*E*_s_(XT) and array expression (except for *atg27310*), suggesting that in this organism X-Plat can perform relatively better for less highly expressed genes (1.6–∼2.2; Figure 8D). Similar expression-level trends were observed for the top five genes with maximum positive and negative Δ*E*_s_(XH) values when comparing X-Plat and HARMONY in rat and human, but not in *Arabidopsis*, and when comparing X-Plat with HARMONY2 for predicted seq expression in rat, but not in *Arabidopsis* or human.

We found that TDM predictions for seq expression values contained many zeros. This apparent artefact was observed across all three organisms, with the proportion of samples receiving zero predictions ranging from 0% to 100% in a gene-dependent manner (Figure 9). The sample frequencies of zero seq value predictions by TDM were inversely correlated with the median array expression value in rat, *Arabidopsis*, and human, indicating that lower-expression genes were more likely to be driven to zero by TDM.

**Figure 9:**
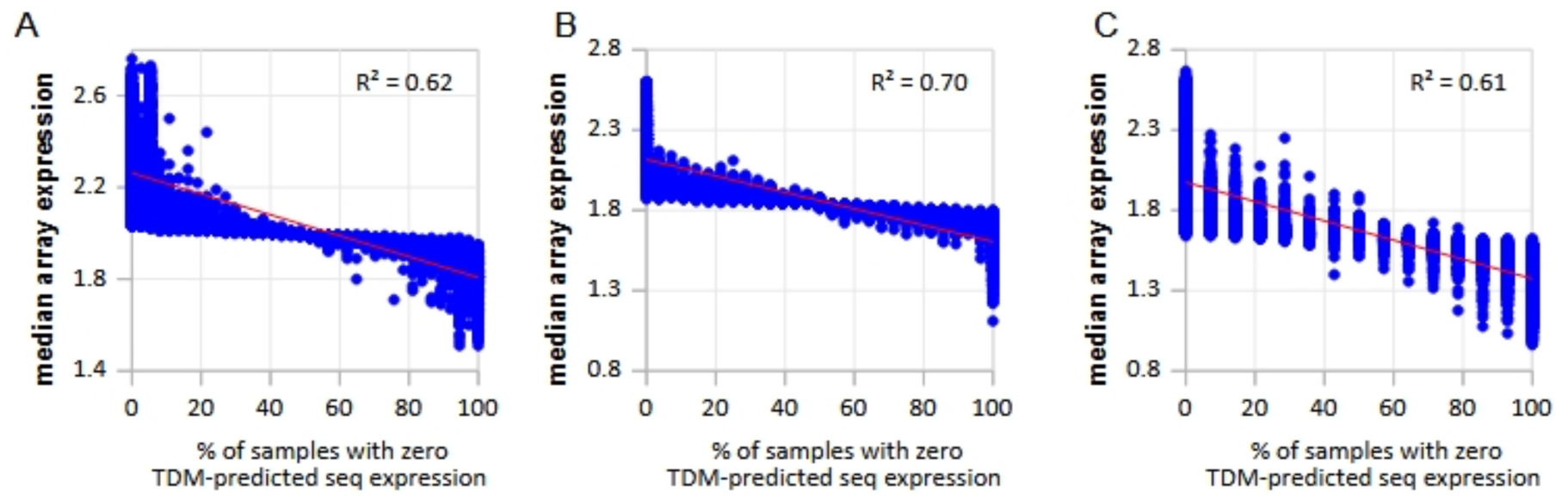
Percentage of samples with zero predicted seq expression values from TDM for rat (A), *Arabidopsis* (B), and human (C) datasets.

## DISCUSSION

We developed X-Plat as a cross-platform conversion tool that enables effective use of legacy gene expression and methylation data for model building, thereby expanding the available sample pool for training and validation. While the transition to next-generation sequencing–based assays is inevitable as microarray platforms become obsolete and reagent supplies are discontinued, X-Plat ensures that the vast amount of data generated using microarrays and single-gene experiments remains valuable and usable. Although several tools have been proposed to transform data between platforms (5, 8, 14–16), to our knowledge, no existing method provides a practically applicable, gene-specific and bidirectional conversion framework that spans both expression and methylation data, and some explicitly restrict sequencing reads to regions where microarray probes were designed (15, 16, 27). By contrast, X-Plat estimates gene-wise mappings that can be applied to new samples without requiring probe-level sequencing read information, making it suitable for large-scale public compendia and routine model deployment.

Following the development of X-Plat, we compared its performance with three widely used conversion tools, TDM (8), HARMONY (26), and HARMONY2 (27). Across platforms and organisms, X-Plat generally outperformed TDM, HARMONY, and HARMONY2, particularly for more highly expressed genes and in the majority of genes within each dataset. In rat, *Arabidopsis*, and human expression datasets, X-Plat achieved lower average cross-validated RMSE than TDM, HARMONY, and HARMONY2 for ≥95% of genes in all sequencing-to-array transformations and in most array-to-sequencing transformations, with only the *Arabidopsis* array-to-sequencing setting showing a slightly lower proportion (∼82%). X-Plat also achieved low RMSE (≤0.2) for the majority of CpG regions in paired human array–sequencing methylation datasets. These findings are consistent with prior work showing that careful cross-platform normalization can enable robust model transfer between microarray and RNA-seq data, but indicate that a gene-wise, bidirectional mapping such as X-Plat can further reduce prediction error across diverse organisms and assays (8, 29). We observed discrepancies in RMSE values for rat compared with *Arabidopsis* and human; this is likely contributed by larger annotation gaps and less mature genome builds for rat, whereas *Arabidopsis* and human benefit from extensive, community-driven and long-term annotation refinement, respectively, which improves mapping and quantification and thus supports more accurate cross-platform modeling (30, 31).

We further investigated factors underlying the observed performance differences among the four tools and found that X-Plat performed particularly well for highly expressed genes across organisms, for both microarray- and sequencing-based data. Expression-range analyses showed that genes with larger dynamic range and higher median expression tended to have lower RMSE under X-Plat than under TDM, HARMONY or HARMONY2, whereas lowly expressed or low-range genes were more likely to show smaller or reversed differences, consistent with the fact that RMSE is dominated by large-magnitude errors. Down-sampling of the training data (Supplementary Figure S2) resulted in only a marginal increase in RMSE for X-Plat predictions of both array and sequencing expression values, indicating that the method is relatively robust to reductions in training sample size. Although TDM occasionally exhibited lower sequencing RMSE than X-Plat, this improvement often—but not always—coincided with its tendency to predict zero expression values for sequencing data; in such cases, the apparent superior performance of TDM reflects an artefact of zero-inflation rather than a genuine gain in predictive accuracy (32). In practical terms, X-Plat provides more stable performance across genes and organisms, with fewer extreme failure modes, whereas TDM, HARMONY, and HARMONY2 tend to underperform except in a small subset of genes where their assumptions happen to align with the data (29).

In summary, our study contributes to broader efforts in cross-platform conversion and integration, addressing a key barrier to reusing legacy datasets (6, 8, 29). Because it is difficult to draw robust conclusions from data generated in any single laboratory—often due to limited sample sizes—it is increasingly important to correlate and integrate data across experiments, laboratories, platforms, assays, and technologies (33, 34). A platform that can comparably analyze data across assays and technologies, such as X-Plat, will help integrate useful legacy data with data generated using modern high-throughput sequencing instruments (29). This can enhance basic hypothesis generation and strengthen conclusions across domains, including disease research, where leveraging both historical microarray cohorts and contemporary sequencing datasets is particularly valuable (6, 35).

## AUTHOR CONTRIBUTIONS

Neeraja M Krishnan: Conceptualization, Formal analysis, Methodology, Visualization, Validation, Writing—original draft, review & editing. Sarah I Rahman: Formal analysis. Lars Røn Olsen: Data generation. Binay Panda: Conceptualization, Methodology, Writing—review & editing.

## SUPPLEMENTARY DATA

Supplementary Data are provided in a separate file.

## DATA AVAILABILITY

Data derived from sources in the public domain: NCBI Gene Expression Omnibus and European Genome-phenome Archive (details on Supplementary Table S1).

