## Supplementary File for "X-Plat: a polynomial regression–based tool for cross-platform transformation of expression and methylation data"

### SUPPLEMENTARY DATA

**Table S1:** Sample-level details for paired microarray–sequencing datasets used to build and test X-Plat. Table S1 is divided into four parts. Parts A–C list expression assay pairs for rat (A), *Arabidopsis* (B), and human (C), respectively, and Part D lists methylation assay pairs for human. For each organism and assay type, the table reports the GEO superseries and series identifiers (superseries, series), microarray and sequencing platforms (platform, platform description), individual sample identifiers and descriptions for microarray (array) and sequencing (seq), and the internal sample code used in this study.

| A superseries | array series | platform | platform desc. | sample | sample desc. | seq series | platform | platform desc. | sample | sample desc. | sample code |
|---|---|---|---|---|---|---|---|---|---|---|---|
| GSE47792 | GSE47875 | GPL1355 | [Rat230_2] Affymetrix Rat Genome 230 2.0 Array | GSM1161441 | Rat_Liver_Vehicle__1d_NN_IP_Rep1 | GSE55347 | GPL17116 | Illumina HiSeq 2000 (Rattus norvegicus) | GSM1336229 | In Vivo Rat Liver Carcinogen (COH_WANG_AB029JACXX_98681_L_CTL_NN_IP_CAGATC_s_4) | CVD040520-002 |
| GSE47792 | GSE47875 | GPL1355 | [Rat230_2] Affymetrix Rat Genome 230 2.0 Array | GSM1161438 | Rat_Liver_Vehicle__5d_NN_IP_Rep6 | GSE55347 | GPL17116 | Illumina HiSeq 2000 (Rattus norvegicus) | GSM1336227 | In Vivo Rat Liver Carcinogen (COH_WANG_AB029JACXX_98611_L_CTL_NN_IP_CAGATC_s_5) | CVD040520-006 |
| GSE47792 | GSE47875 | GPL1355 | [Rat230_2] Affymetrix Rat Genome 230 2.0 Array | GSM1161469 | Rat_Liver_N-NITROSODIMETHYLAMINE_10mg/kg_5d_NN_IP_Rep2 | GSE55347 | GPL17116 | Illumina HiSeq 2000 (Rattus norvegicus) | GSM1336208 | In Vivo Rat Liver Carcinogen (COH_WANG_AB0142ACXX_99746_L_NIT_NN_IP_TGACCA_s_1) | CVD040520-116 |
| GSE47792 | GSE47875 | GPL1355 | [Rat230_2] Affymetrix Rat Genome 230 2.0 Array | GSM1161479 | Rat_Liver_N-NITROSODIMETHYLAMINE_10mg/kg_5d_NN_IP_Rep3 | GSE55347 | GPL17116 | Illumina HiSeq 2000 (Rattus norvegicus) | GSM1336209 | In Vivo Rat Liver Carcinogen (COH_WANG_AB0142ACXX_99747_L_NIT_NN_IP_TGACCA_s_2) | CVD040520-117 |
| GSE47792 | GSE47875 | GPL1355 | [Rat230_2] Affymetrix Rat Genome 230 2.0 Array | GSM1161467 | Rat_Liver_N-NITROSODIMETHYLAMINE_10mg/kg_5d_NN_IP_Rep1 | GSE55347 | GPL17116 | Illumina HiSeq 2000 (Rattus norvegicus) | GSM1336207 | In Vivo Rat Liver Carcinogen (COH_WANG_AB0142ACXX_99697_L_NIT_NN_IP_TGACCA_s_3) | CVD040520-118 |
| GSE47792 | GSE47875 | GPL1355 | [Rat230_2] Affymetrix Rat Genome 230 2.0 Array | GSM1161437 | Rat_Liver_THIOACETAMIDE_200mg/kg_5d_NN_IP_Rep1 | GSE55347 | GPL17116 | Illumina HiSeq 2000 (Rattus norvegicus) | GSM1336189 | In Vivo Rat Liver Carcinogen (COH_WANG_AB0142ACXX_98573_L_THI_NN_IP_CGATGT_s_1) | CVD040520-130 |
| GSE47792 | GSE47875 | GPL1355 | [Rat230_2] Affymetrix Rat Genome 230 2.0 Array | GSM1161440 | Rat_Liver_THIOACETAMIDE_200mg/kg_5d_NN_IP_Rep2 | GSE55347 | GPL17116 | Illumina HiSeq 2000 (Rattus norvegicus) | GSM1336192 | In Vivo Rat Liver Carcinogen (COH_WANG_AB0142ACXX_98678_L_THI_NN_IP_CGATGT_s_2) | CVD040520-131 |
| GSE47792 | GSE47875 | GPL1355 | [Rat230_2] Affymetrix Rat Genome 230 2.0 Array | GSM1161442 | Rat_Liver_THIOACETAMIDE_200mg/kg_5d_NN_IP_Rep3 | GSE55347 | GPL17116 | Illumina HiSeq 2000 (Rattus norvegicus) | GSM1336193 | In Vivo Rat Liver Carcinogen (COH_WANG_AB0142ACXX_98726_L_THI_NN_IP_CGATGT_s_3) | CVD040520-132 |
| GSE47792 | GSE47875 | GPL1355 | [Rat230_2] | GSM1161464 | Rat_Liver_IFOSFA | GSE55347 | GPL17116 | Illumina HiSeq | GSM1336205 | In Vivo Rat Liver | I011106-043 |

| | | | | | | | | | | | |
|---|---|---|---|---|---|---|---|---|---|---|---|
| | | | Affymetrix Rat Genome 230 2.0 Array | | MIDE_143mg/kg_3d_NN_OG_Rep2 | | | 2000 (Rattus norvegicus) | | Carcinogen (COH_WANG_AB0142ACXX_99564_L_IFO_NN_OG_TGACCA_s_4) | |
| GSE47792 | GSE47875 | GPL1355 | [Rat230_2] Affymetrix Rat Genome 230 2.0 Array | GSM1161478 | Rat_Liver_IFOSFAMIDE_143mg/kg_3d_NN_OG_Rep3 | GSE55347 | GPL17116 | Illumina HiSeq 2000 (Rattus norvegicus) | GSM1336206 | In Vivo Rat Liver Carcinogen (COH_WANG_AB0142ACXX_99586_L_IFO_NN_OG_TGACCA_s_5) | I011106-044 |
| GSE47792 | GSE47875 | GPL1355 | [Rat230_2] Affymetrix Rat Genome 230 2.0 Array | GSM1161463 | Rat_Liver_IFOSFAMIDE_143mg/kg_3d_NN_OG_Rep1 | GSE55347 | GPL17116 | Illumina HiSeq 2000 (Rattus norvegicus) | GSM1336204 | In Vivo Rat Liver Carcinogen (COH_WANG_AB0142ACXX_99534_L_IFO_NN_OG_TGACCA_s_6) | I011106-045 |
| GSE47792 | GSE47875 | GPL1355 | [Rat230_2] Affymetrix Rat Genome 230 2.0 Array | GSM1161428 | Rat_Liver_Vehicle__5d_NN_OG_Rep2 | GSE55347 | GPL17116 | Illumina HiSeq 2000 (Rattus norvegicus) | GSM1336223 | In Vivo Rat Liver Carcinogen (COH_WANG_AB029JACXX_98432_L_CTL_NN_OG_TAGCTT_s_1) | R020514-019 |
| GSE47792 | GSE47875 | GPL1355 | [Rat230_2] Affymetrix Rat Genome 230 2.0 Array | GSM1161428 | Rat_Liver_Vehicle__5d_NN_OG_Rep2 | GSE55347 | GPL17116 | Illumina HiSeq 2000 (Rattus norvegicus) | GSM1336254 | In Vivo Rat Liver Carcinogen (COH_WANG_AC0HK2ACXX_542_CTTGTA_s_2) | R020514-019 |
| GSE47792 | GSE47875 | GPL1355 | [Rat230_2] Affymetrix Rat Genome 230 2.0 Array | GSM1161428 | Rat_Liver_Vehicle__5d_NN_OG_Rep2 | GSE55347 | GPL14844 | Illumina HiScanSQ (Rattus norvegicus) | GSM1336289 | In Vivo Rat Liver Carcinogen (COH_WANG_BC064YACXX_98432_L_CTL_NN_OG_TAGCTT_s_5) | R020514-019 |
| GSE47792 | GSE47875 | GPL1355 | [Rat230_2] Affymetrix Rat Genome 230 2.0 Array | GSM1161439 | Rat_Liver_Vehicle__5d_NN_OG_Rep4 | GSE55347 | GPL17116 | Illumina HiSeq 2000 (Rattus norvegicus) | GSM1336228 | In Vivo Rat Liver Carcinogen (COH_WANG_AB029JACXX_98619_L_CTL_NN_OG_TAGCTT_s_2) | R030409-001 |
| GSE47792 | GSE47875 | GPL1355 | [Rat230_2] Affymetrix Rat Genome 230 2.0 Array | GSM1161439 | Rat_Liver_Vehicle__5d_NN_OG_Rep4 | GSE55347 | GPL17116 | Illumina HiSeq 2000 (Rattus norvegicus) | GSM1336264 | In Vivo Rat Liver Carcinogen (COH_WANG_AC0HK2ACXX_558_GCCAAT_s_5) | R030409-001 |
| GSE47792 | GSE47875 | GPL1355 | [Rat230_2] Affymetrix Rat Genome 230 2.0 Array | GSM1161439 | Rat_Liver_Vehicle__5d_NN_OG_Rep4 | GSE55347 | GPL14844 | Illumina HiScanSQ (Rattus norvegicus) | GSM1336292 | In Vivo Rat Liver Carcinogen (COH_WANG_BC064YACXX_98619_L_CTL_NN_OG_TAGCTT_s_4) | R030409-001 |
| GSE47792 | GSE47875 | GPL1355 | [Rat230_2] Affymetrix Rat | GSM1161443 | Rat_Liver_Vehicle__5d_NN_OG_Rep5 | GSE55347 | GPL17116 | Illumina HiSeq 2000 (Rattus | GSM1336230 | In Vivo Rat Liver Carcinogen | R030409-003 |

| | | | | | | | | | | | |
|---|---|---|---|---|---|---|---|---|---|---|---|
| | | | Genome 230 2.0 Array | | | | | norvegicus) | | (COH_WANG_AB029JACXX_98741_L_CTL_NN_OG_GGCTAC_s_3) | |
| GSE47792 | GSE47875 | GPL1355 | [Rat230_2] Affymetrix Rat Genome 230 2.0 Array | GSM1161443 | Rat_Liver_Vehicle_5d_NN_OG_Rep5 | GSE55347 | GPL17116 | Illumina HiSeq 2000 (Rattus norvegicus) | GSM1336280 | In Vivo Rat Liver Carcinogen (COH_WANG_AC0HK2ACXX_574_TGACCA_s_8) | R030409-003 |
| GSE47792 | GSE47875 | GPL1355 | [Rat230_2] Affymetrix Rat Genome 230 2.0 Array | GSM1161443 | Rat_Liver_Vehicle_5d_NN_OG_Rep5 | GSE55347 | GPL14844 | Illumina HiScanSQ (Rattus norvegicus) | GSM1336293 | In Vivo Rat Liver Carcinogen (COH_WANG_BC064YACXX_98741_L_CTL_NN_OG_GGCTAC_s_5) | R030409-003 |
| GSE47792 | GSE47875 | GPL1355 | [Rat230_2] Affymetrix Rat Genome 230 2.0 Array | GSM1161435 | Rat_Liver_Vehicle_5d_NN_OG_Rep3 | GSE55347 | GPL14844 | Illumina HiScanSQ (Rattus norvegicus) | GSM1336291 | In Vivo Rat Liver Carcinogen (COH_WANG_BC064YACXX_98543_L_CTL_NN_OG_GGCTAC_s_2) | R030916-024 |
| GSE47792 | GSE47875 | GPL1355 | [Rat230_2] Affymetrix Rat Genome 230 2.0 Array | GSM1161453 | Rat_Liver_3-METHYLCHOLANTHRENE_300mg/kg_5d_NN_OG_Rep3 | GSE55347 | GPL17116 | Illumina HiSeq 2000 (Rattus norvegicus) | GSM1336225 | In Vivo Rat Liver Carcinogen (COH_WANG_AB029JACXX_98512_L_3ME_NN_OG_ATCACG_s_1) | R030916-046 |
| GSE47792 | GSE47875 | GPL1355 | [Rat230_2] Affymetrix Rat Genome 230 2.0 Array | GSM1161450 | Rat_Liver_3-METHYLCHOLANTHRENE_300mg/kg_5d_NN_OG_Rep1 | GSE55347 | GPL17116 | Illumina HiSeq 2000 (Rattus norvegicus) | GSM1336220 | In Vivo Rat Liver Carcinogen (COH_WANG_AB029JACXX_98326_L_3ME_NN_OG_ATCACG_s_2) | R030916-047 |
| GSE47792 | GSE47875 | GPL1355 | [Rat230_2] Affymetrix Rat Genome 230 2.0 Array | GSM1161451 | Rat_Liver_3-METHYLCHOLANTHRENE_300mg/kg_5d_NN_OG_Rep2 | GSE55347 | GPL17116 | Illumina HiSeq 2000 (Rattus norvegicus) | GSM1336222 | In Vivo Rat Liver Carcinogen (COH_WANG_AB029JACXX_98388_L_3ME_NN_OG_ATCACG_s_3) | R030916-048 |
| GSE47792 | GSE47875 | GPL1355 | [Rat230_2] Affymetrix Rat Genome 230 2.0 Array | GSM1161431 | Rat_Liver_BETA-NAPHTHOFLAVONE_1500mg/kg_5d_NN_OG_Rep1 | GSE55347 | GPL17116 | Illumina HiSeq 2000 (Rattus norvegicus) | GSM1336224 | In Vivo Rat Liver Carcinogen (COH_WANG_AB029JACXX_98449_L_NAP_NN_OG_CGATGT_s_1) | R030916-070 |
| GSE47792 | GSE47875 | GPL1355 | [Rat230_2] Affymetrix Rat Genome 230 2.0 Array | GSM1161434 | Rat_Liver_BETA-NAPHTHOFLAVONE_1500mg/kg_5d_NN_OG_Rep2 | GSE55347 | GPL17116 | Illumina HiSeq 2000 (Rattus norvegicus) | GSM1336226 | In Vivo Rat Liver Carcinogen (COH_WANG_AB029JACXX_98524_L_NAP_NN_OG_CGATGT_s_2) | R030916-071 |
| GSE47792 | GSE47875 | GPL1355 | [Rat230_2] Affymetrix Rat Genome 230 2.0 | GSM1161470 | Rat_Liver_BETA-NAPHTHOFLAVONE_1500mg/ | GSE55347 | GPL17116 | Illumina HiSeq 2000 (Rattus norvegicus) | GSM1336221 | In Vivo Rat Liver Carcinogen (COH_WANG_AB029JA | R030916-072 |

| | | | | | | | | | | | |
|---|---|---|---|---|---|---|---|---|---|---|---|
| | | | Array | | kg_5d_NN_OG_Rep3 | | | | | CXX_98377_L_NAP_NN_OG_CGATGT_s_3) | |
| GSE47792 | GSE47875 | GPL1355 | [Rat230_2] Affymetrix Rat Genome 230 2.0 Array | GSM1161452 | Rat_Liver_Vehicle__5d_NN_OG_Rep6 | GSE55347 | GPL14844 | Illumina HiScanSQ (Rattus norvegicus) | GSM1336290 | In Vivo Rat Liver Carcinogen (COH_WANG_BC064YACXX_98482_L_CTL_NN_OG_TGACCA_s_2) | R031014-023 |
| GSE47792 | GSE47875 | GPL1355 | [Rat230_2] Affymetrix Rat Genome 230 2.0 Array | GSM1161426 | Rat_Liver_Vehicle__1d_NU_OG_Rep1 | GSE55347 | GPL14844 | Illumina HiScanSQ (Rattus norvegicus) | GSM1336288 | In Vivo Rat Liver Carcinogen (COH_WANG_BC064YACXX_98348_L_CTL_NU_OG_CGATGT_s_3) | S010924-004 |
| GSE47792 | GSE47875 | GPL1355 | [Rat230_2] Affymetrix Rat Genome 230 2.0 Array | GSM1161430 | Rat_Liver_Vehicle__3d_NU_OG_Rep2 | GSE55347 | GPL14844 | Illumina HiScanSQ (Rattus norvegicus) | GSM1336287 | In Vivo Rat Liver Carcinogen (COH_WANG_BC064YACXX_98329_L_CTL_NU_OG_ACAGTG_s_4) | S010924-008 |
| GSE47792 | GSE47875 | GPL1355 | [Rat230_2] Affymetrix Rat Genome 230 2.0 Array | GSM1161511 | Rat_Liver_Vehicle__7d_NU_OG_Test_Rep6 | GSE55347 | GPL17116 | Illumina HiSeq 2000 (Rattus norvegicus) | GSM1336274 | In Vivo Rat Liver Carcinogen (COH_WANG_AC0HK2ACXX_568_TGACCA_s_7) | S010924-010 |
| GSE47792 | GSE47875 | GPL1355 | [Rat230_2] Affymetrix Rat Genome 230 2.0 Array | GSM1161444 | Rat_Liver_Vehicle__7d_NU_OG_Rep6 | GSE55347 | GPL17116 | Illumina HiSeq 2000 (Rattus norvegicus) | GSM1336196 | In Vivo Rat Liver Carcinogen (COH_WANG_AB0142ACXX_98848_L_CTL_NU_OG_GCCAAT_s_8) | S010924-011 |
| GSE47792 | GSE47875 | GPL1355 | [Rat230_2] Affymetrix Rat Genome 230 2.0 Array | GSM1161445 | Rat_Liver_CARBON-TETRACHLORIDE_1175mg/kg_7d_NU_OG_Rep1 | GSE55347 | GPL17116 | Illumina HiSeq 2000 (Rattus norvegicus) | GSM1336197 | In Vivo Rat Liver Carcinogen (COH_WANG_AB0142ACXX_98858_L_CAR_NU_OG_ACAGTG_s_3) | S010924-082 |
| GSE47792 | GSE47875 | GPL1355 | [Rat230_2] Affymetrix Rat Genome 230 2.0 Array | GSM1161449 | Rat_Liver_CARBON-TETRACHLORIDE_1175mg/kg_7d_NU_OG_Rep2 | GSE55347 | GPL17116 | Illumina HiSeq 2000 (Rattus norvegicus) | GSM1336199 | In Vivo Rat Liver Carcinogen (COH_WANG_AB0142ACXX_98873_L_CAR_NU_OG_ACAGTG_s_4) | S010924-083 |
| GSE47792 | GSE47875 | GPL1355 | [Rat230_2] Affymetrix Rat Genome 230 2.0 Array | GSM1161468 | Rat_Liver_Vehicle__1d_NN_IP_Rep4 | GSE55347 | GPL17116 | Illumina HiSeq 2000 (Rattus norvegicus) | GSM1336236 | In Vivo Rat Liver Carcinogen (COH_WANG_AB029JACXX_99728_L_CTL_NN_IP_ACTTGA_s_6) | S011002-030 |
| GSE47792 | GSE47875 | GPL1355 | [Rat230_2] Affymetrix Rat | GSM1161459 | Rat_Liver_Vehicle__3d_NN_IP_Rep5 | GSE55347 | GPL14844 | Illumina HiScanSQ | GSM1336294 | In Vivo Rat Liver Carcinogen | S011002-033 |

| | | | | | | | | | | |
|---|---|---|---|---|---|---|---|---|---|---|
| | | | Genome 230 2.0 Array | | | | | (Rattus norvegicus) | | (COH_WANG_BC064YACXX_98893_L_CTL_NN_IP_ACTTGA_s_2) | |
| GSE47792 | GSE47875 | GPL1355 | [Rat230_2] Affymetrix Rat Genome 230 2.0 Array | GSM1161448 | Rat_Liver_BEZAFIBRATE_617mg/kg_7d_NU_OG_Rep1 | GSE55347 | GPL17116 | Illumina HiSeq 2000 (Rattus norvegicus) | GSM1336194 | In Vivo Rat Liver Carcinogen (COH_WANG_AB0142ACXX_98800_L_BEZ_NU_OG_ACAGTG_s_6) | S011002-178 |
| GSE47792 | GSE47875 | GPL1355 | [Rat230_2] Affymetrix Rat Genome 230 2.0 Array | GSM1161471 | Rat_Liver_BEZAFIBRATE_617mg/kg_7d_NU_OG_Rep3 | GSE55347 | GPL17116 | Illumina HiSeq 2000 (Rattus norvegicus) | GSM1336190 | In Vivo Rat Liver Carcinogen (COH_WANG_AB0142ACXX_98576_L_BEZ_NU_OG_ACAGTG_s_7) | S011002-179 |
| GSE47792 | GSE47875 | GPL1355 | [Rat230_2] Affymetrix Rat Genome 230 2.0 Array | GSM1161458 | Rat_Liver_BEZAFIBRATE_617mg/kg_7d_NU_OG_Rep2 | GSE55347 | GPL17116 | Illumina HiSeq 2000 (Rattus norvegicus) | GSM1336191 | In Vivo Rat Liver Carcinogen (COH_WANG_AB0142ACXX_98658_L_BEZ_NU_OG_ACAGTG_s_8) | S011002-180 |
| GSE47792 | GSE47875 | GPL1355 | [Rat230_2] Affymetrix Rat Genome 230 2.0 Array | GSM1161509 | Rat_Liver_GEMFIBROZIL_700mg/kg_7d_NU_OG_Rep1 | GSE55347 | GPL17116 | Illumina HiSeq 2000 (Rattus norvegicus) | GSM1336257 | In Vivo Rat Liver Carcinogen (COH_WANG_AC0HK2ACXX_545_ACAGTG_s_3) | S011002-226 |
| GSE47792 | GSE47875 | GPL1355 | [Rat230_2] Affymetrix Rat Genome 230 2.0 Array | GSM1161513 | Rat_Liver_GEMFIBROZIL_700mg/kg_7d_NU_OG_Rep3 | GSE55347 | GPL17116 | Illumina HiSeq 2000 (Rattus norvegicus) | GSM1336267 | In Vivo Rat Liver Carcinogen (COH_WANG_AC0HK2ACXX_561_CGATGT_s_6) | S011002-227 |
| GSE47792 | GSE47875 | GPL1355 | [Rat230_2] Affymetrix Rat Genome 230 2.0 Array | GSM1161510 | Rat_Liver_GEMFIBROZIL_700mg/kg_7d_NU_OG_Rep2 | GSE55347 | GPL17116 | Illumina HiSeq 2000 (Rattus norvegicus) | GSM1336283 | In Vivo Rat Liver Carcinogen (COH_WANG_AC0HK2ACXX_577_CAGATC_s_8) | S011002-228 |
| GSE47792 | GSE47875 | GPL1355 | [Rat230_2] Affymetrix Rat Genome 230 2.0 Array | GSM1161483 | Rat_Liver_LOVASTATIN_450mg/kg_5d_NU_OG_Rep1 | GSE55347 | GPL17116 | Illumina HiSeq 2000 (Rattus norvegicus) | GSM1336252 | In Vivo Rat Liver Carcinogen (COH_WANG_AC0HK2ACXX_540_GCCAAT_s_2) | S011009-022 |
| GSE47792 | GSE47875 | GPL1355 | [Rat230_2] Affymetrix Rat Genome 230 2.0 Array | GSM1161491 | Rat_Liver_LOVASTATIN_450mg/kg_5d_NU_OG_Rep2 | GSE55347 | GPL17116 | Illumina HiSeq 2000 (Rattus norvegicus) | GSM1336262 | In Vivo Rat Liver Carcinogen (COH_WANG_AC0HK2ACXX_556_TGACCA_s_5) | S011009-023 |
| GSE47792 | GSE47875 | GPL1355 | [Rat230_2] Affymetrix Rat Genome 230 2.0 | GSM1161495 | Rat_Liver_LOVASTATIN_450mg/kg_5d_NU_OG_Re | GSE55347 | GPL17116 | Illumina HiSeq 2000 (Rattus norvegicus) | GSM1336278 | In Vivo Rat Liver Carcinogen (COH_WANG_AC0HK2 | S011009-024 |

| | | | | | | | | | | | |
|---|---|---|---|---|---|---|---|---|---|---|---|
| | | | Array | | p3 | | | | | ACXX_572_CTTGTA_s _7) | |
| GSE47792 | GSE47875 | GPL1355 | [Rat230_2] Affymetrix Rat Genome 230 2.0 Array | GSM1161461 | Rat_Liver_LEFLUN OMIDE_60mg/ kg_5d_NU_OG_Re p2 | GSE55347 | GPL17116 | Illumina HiSeq 2000 (Rattus norvegicus) | GSM1336232 | In Vivo Rat Liver Carcinogen (COH_WANG_AB029JA CXX_99414_L_LEF_NU _OG_TTAGGC_s_4) | S011009-130 |
| GSE47792 | GSE47875 | GPL1355 | [Rat230_2] Affymetrix Rat Genome 230 2.0 Array | GSM1161462 | Rat_Liver_LEFLUN OMIDE_60mg/ kg_5d_NU_OG_Re p3 | GSE55347 | GPL17116 | Illumina HiSeq 2000 (Rattus norvegicus) | GSM1336233 | In Vivo Rat Liver Carcinogen (COH_WANG_AB029JA CXX_99430_L_LEF_NU _OG_TTAGGC_s_5) | S011009-131 |
| GSE47792 | GSE47875 | GPL1355 | [Rat230_2] Affymetrix Rat Genome 230 2.0 Array | GSM1161457 | Rat_Liver_LEFLUN OMIDE_60mg/ kg_5d_NU_OG_Re p1 | GSE55347 | GPL17116 | Illumina HiSeq 2000 (Rattus norvegicus) | GSM1336231 | In Vivo Rat Liver Carcinogen (COH_WANG_AB029JA CXX_99358_L_LEF_NU _OG_TTAGGC_s_6) | S011009-132 |
| GSE47792 | GSE47875 | GPL1355 | [Rat230_2] Affymetrix Rat Genome 230 2.0 Array | GSM1161422 | Rat_Liver_ECONAZ OLE_334mg/ kg_5d_NU_OG_Re p3 | GSE55347 | GPL17116 | Illumina HiSeq 2000 (Rattus norvegicus) | GSM1336218 | In Vivo Rat Liver Carcinogen (COH_WANG_AB029JA CXX_97577_L_ECO_N U_OG_TGACCA_s_4) | S011016-154 |
| GSE47792 | GSE47875 | GPL1355 | [Rat230_2] Affymetrix Rat Genome 230 2.0 Array | GSM1161420 | Rat_Liver_ECONAZ OLE_334mg/ kg_5d_NU_OG_Re p1 | GSE55347 | GPL17116 | Illumina HiSeq 2000 (Rattus norvegicus) | GSM1336216 | In Vivo Rat Liver Carcinogen (COH_WANG_AB029JA CXX_97186_L_ECO_N U_OG_TGACCA_s_5) | S011016-155 |
| GSE47792 | GSE47875 | GPL1355 | [Rat230_2] Affymetrix Rat Genome 230 2.0 Array | GSM1161421 | Rat_Liver_ECONAZ OLE_334mg/ kg_5d_NU_OG_Re p2 | GSE55347 | GPL17116 | Illumina HiSeq 2000 (Rattus norvegicus) | GSM1336217 | In Vivo Rat Liver Carcinogen (COH_WANG_AB029JA CXX_97266_L_ECO_N U_OG_TGACCA_s_6) | S011016-156 |
| GSE47792 | GSE47875 | GPL1355 | [Rat230_2] Affymetrix Rat Genome 230 2.0 Array | GSM1161486 | Rat_Liver_MICONA ZOLE_920mg/ kg_5d_NU_OG_Re p1 | GSE55347 | GPL17116 | Illumina HiSeq 2000 (Rattus norvegicus) | GSM1336246 | In Vivo Rat Liver Carcinogen (COH_WANG_AC0HK2 ACXX_534_GCCAAT_s _1) | S011016-178 |
| GSE47792 | GSE47875 | GPL1355 | [Rat230_2] Affymetrix Rat Genome 230 2.0 Array | GSM1161514 | Rat_Liver_MICONA ZOLE_920mg/ kg_5d_NU_OG_Re p2 | GSE55347 | GPL17116 | Illumina HiSeq 2000 (Rattus norvegicus) | GSM1336238 | In Vivo Rat Liver Carcinogen (COH_WANG_AC098L ACXX_550_TGACCA_s _7) | S011016-179 |
| GSE47792 | GSE47875 | GPL1355 | [Rat230_2] Affymetrix Rat Genome 230 2.0 Array | GSM1161515 | Rat_Liver_MICONA ZOLE_920mg/ kg_5d_NU_OG_Re p3 | GSE55347 | GPL17116 | Illumina HiSeq 2000 (Rattus norvegicus) | GSM1336272 | In Vivo Rat Liver Carcinogen (COH_WANG_AC0HK2 ACXX_566_CTTGTA_s | S011016-180 |

| | | | | | | | | | | | |
|---|---|---|---|---|---|---|---|---|---|---|---|
| | | | | | | | | | | _6) | |
| GSE47792 | GSE47875 | GPL1355 | [Rat230_2] Affymetrix Rat Genome 230 2.0 Array | GSM1161516 | Rat_Liver_ETHINYLESTRADIOL_10mg/kg_5d_NU_OG_Rep1 | GSE55347 | GPL17116 | Illumina HiSeq 2000 (Rattus norvegicus) | GSM1336249 | In Vivo Rat Liver Carcinogen (COH_WANG_AC0HK2ACXX_537_CGATGT_s_2) | S011023-082 |
| GSE47792 | GSE47875 | GPL1355 | [Rat230_2] Affymetrix Rat Genome 230 2.0 Array | GSM1161522 | Rat_Liver_ETHINYLESTRADIOL_10mg/kg_5d_NU_OG_Rep2 | GSE55347 | GPL17116 | Illumina HiSeq 2000 (Rattus norvegicus) | GSM1336241 | In Vivo Rat Liver Carcinogen (COH_WANG_AC098LACXX_553_CAGATC_s_7) | S011023-083 |
| GSE47792 | GSE47875 | GPL1355 | [Rat230_2] Affymetrix Rat Genome 230 2.0 Array | GSM1161524 | Rat_Liver_ETHINYLESTRADIOL_10mg/kg_5d_NU_OG_Rep3 | GSE55347 | GPL17116 | Illumina HiSeq 2000 (Rattus norvegicus) | GSM1336275 | In Vivo Rat Liver Carcinogen (COH_WANG_AC0HK2ACXX_569_ACAGTG_s_7) | S011023-084 |
| GSE47792 | GSE47875 | GPL1355 | [Rat230_2] Affymetrix Rat Genome 230 2.0 Array | GSM1161488 | Rat_Liver_FLUCONAZOLE_394mg/kg_5d_NU_OG_Rep1 | GSE55347 | GPL17116 | Illumina HiSeq 2000 (Rattus norvegicus) | GSM1336245 | In Vivo Rat Liver Carcinogen (COH_WANG_AC0HK2ACXX_533_ACAGTG_s_1) | S011023-214 |
| GSE47792 | GSE47875 | GPL1355 | [Rat230_2] Affymetrix Rat Genome 230 2.0 Array | GSM1161493 | Rat_Liver_FLUCONAZOLE_394mg/kg_5d_NU_OG_Rep2 | GSE55347 | GPL17116 | Illumina HiSeq 2000 (Rattus norvegicus) | GSM1336237 | In Vivo Rat Liver Carcinogen (COH_WANG_AC098LACXX_549_CGATGT_s_7) | S011023-215 |
| GSE47792 | GSE47875 | GPL1355 | [Rat230_2] Affymetrix Rat Genome 230 2.0 Array | GSM1161507 | Rat_Liver_FLUCONAZOLE_394mg/kg_5d_NU_OG_Rep3 | GSE55347 | GPL17116 | Illumina HiSeq 2000 (Rattus norvegicus) | GSM1336271 | In Vivo Rat Liver Carcinogen (COH_WANG_AC0HK2ACXX_565_CAGATC_s_6) | S011023-216 |
| GSE47792 | GSE47875 | GPL1355 | [Rat230_2] Affymetrix Rat Genome 230 2.0 Array | GSM1161517 | Rat_Liver_ROSIGLITAZONE_1800mg/kg_5d_NU_OG_Rep1 | GSE55347 | GPL17116 | Illumina HiSeq 2000 (Rattus norvegicus) | GSM1336256 | In Vivo Rat Liver Carcinogen (COH_WANG_AC0HK2ACXX_544_TGACCA_s_3) | S011030-046 |
| GSE47792 | GSE47875 | GPL1355 | [Rat230_2] Affymetrix Rat Genome 230 2.0 Array | GSM1161518 | Rat_Liver_ROSIGLITAZONE_1800mg/kg_5d_NU_OG_Rep2 | GSE55347 | GPL17116 | Illumina HiSeq 2000 (Rattus norvegicus) | GSM1336266 | In Vivo Rat Liver Carcinogen (COH_WANG_AC0HK2ACXX_560_CTTGTA_s_5) | S011030-047 |
| GSE47792 | GSE47875 | GPL1355 | [Rat230_2] Affymetrix Rat Genome 230 2.0 Array | GSM1161519 | Rat_Liver_ROSIGLITAZONE_1800mg/kg_5d_NU_OG_Rep3 | GSE55347 | GPL17116 | Illumina HiSeq 2000 (Rattus norvegicus) | GSM1336282 | In Vivo Rat Liver Carcinogen (COH_WANG_AC0HK2ACXX_576_GCCAAT_s_8) | S011030-048 |

| | | | | | | | | | | |
|---|---|---|---|---|---|---|---|---|---|---|
| GSE47792 | GSE47875 | GPL1355 | [Rat230_2] Affymetrix Rat Genome 230 2.0 Array | GSM1161521 | Rat_Liver_SIMVASTATIN_1200mg/kg_3d_NU_OG_Rep2 | GSE55347 | GPL17116 | Illumina HiSeq 2000 (Rattus norvegicus) | GSM1336263 | In Vivo Rat Liver Carcinogen (COH_WANG_AC0HK2ACXX_557_ACAGTG_s_5) | S011030-115 |
| GSE47792 | GSE47875 | GPL1355 | [Rat230_2] Affymetrix Rat Genome 230 2.0 Array | GSM1161523 | Rat_Liver_SIMVASTATIN_1200mg/kg_3d_NU_OG_Rep3 | GSE55347 | GPL17116 | Illumina HiSeq 2000 (Rattus norvegicus) | GSM1336279 | In Vivo Rat Liver Carcinogen (COH_WANG_AC0HK2ACXX_573_CGATGT_s_8) | S011030-116 |
| GSE47792 | GSE47875 | GPL1355 | [Rat230_2] Affymetrix Rat Genome 230 2.0 Array | GSM1161520 | Rat_Liver_SIMVASTATIN_1200mg/kg_3d_NU_OG_Rep1 | GSE55347 | GPL17116 | Illumina HiSeq 2000 (Rattus norvegicus) | GSM1336253 | In Vivo Rat Liver Carcinogen (COH_WANG_AC0HK2ACXX_541_CAGATC_s_2) | S011030-117 |
| GSE47792 | GSE47875 | GPL1355 | [Rat230_2] Affymetrix Rat Genome 230 2.0 Array | GSM1161496 | Rat_Liver_NORETHINDRONE_375mg/kg_5d_NU_OG_Rep3 | GSE55347 | GPL17116 | Illumina HiSeq 2000 (Rattus norvegicus) | GSM1336276 | In Vivo Rat Liver Carcinogen (COH_WANG_AC0HK2ACXX_570_GCCAAT_s_7) | S011030-142 |
| GSE47792 | GSE47875 | GPL1355 | [Rat230_2] Affymetrix Rat Genome 230 2.0 Array | GSM1161484 | Rat_Liver_NORETHINDRONE_375mg/kg_5d_NU_OG_Rep1 | GSE55347 | GPL17116 | Illumina HiSeq 2000 (Rattus norvegicus) | GSM1336250 | In Vivo Rat Liver Carcinogen (COH_WANG_AC0HK2ACXX_538_TGACCA_s_2) | S011030-143 |
| GSE47792 | GSE47875 | GPL1355 | [Rat230_2] Affymetrix Rat Genome 230 2.0 Array | GSM1161489 | Rat_Liver_Vehicle__5d_NU_OG_Test_Rep1 | GSE55347 | GPL17116 | Illumina HiSeq 2000 (Rattus norvegicus) | GSM1336243 | In Vivo Rat Liver Carcinogen (COH_WANG_AC0HK2ACXX_531_CGATGT_s_1) | S011113-022 |
| GSE47792 | GSE47875 | GPL1355 | [Rat230_2] Affymetrix Rat Genome 230 2.0 Array | GSM1161498 | Rat_Liver_BETA-ESTRADIOL_150mg/kg_5d_NU_OG_Rep3 | GSE55347 | GPL17116 | Illumina HiSeq 2000 (Rattus norvegicus) | GSM1336273 | In Vivo Rat Liver Carcinogen (COH_WANG_AC0HK2ACXX_567_CGATGT_s_7) | S011113-046 |
| GSE47792 | GSE47875 | GPL1355 | [Rat230_2] Affymetrix Rat Genome 230 2.0 Array | GSM1161487 | Rat_Liver_BETA-ESTRADIOL_150mg/kg_5d_NU_OG_Rep1 | GSE55347 | GPL17116 | Illumina HiSeq 2000 (Rattus norvegicus) | GSM1336247 | In Vivo Rat Liver Carcinogen (COH_WANG_AC0HK2ACXX_535_CAGATC_s_1) | S011113-047 |
| GSE47792 | GSE47875 | GPL1355 | [Rat230_2] Affymetrix Rat Genome 230 2.0 Array | GSM1161494 | Rat_Liver_BETA-ESTRADIOL_150mg/kg_5d_NU_OG_Rep2 | GSE55347 | GPL17116 | Illumina HiSeq 2000 (Rattus norvegicus) | GSM1336239 | In Vivo Rat Liver Carcinogen (COH_WANG_AC098LACXX_551_ACAGTG_s_7) | S011113-048 |
| GSE47792 | GSE47875 | GPL1355 | [Rat230_2] | GSM1161497 | Rat_Liver_CLOTRI | GSE55347 | GPL17116 | Illumina HiSeq | GSM1336270 | In Vivo Rat Liver | S011127-130 |

| | | | | | | | | | | | |
|---|---|---|---|---|---|---|---|---|---|---|---|
| | | | Affymetrix Rat Genome 230 2.0 Array | | MAZOLE_89mg/kg_5d_NU_OG_Rep3 | | | 2000 (Rattus norvegicus) | | Carcinogen (COH_WANG_AC0HK2ACXX_564_GCCAAT_s_6) | |
| GSE47792 | GSE47875 | GPL1355 | [Rat230_2] Affymetrix Rat Genome 230 2.0 Array | GSM1161485 | Rat_Liver_CLOTRIMAZOLE_89mg/kg_5d_NU_OG_Rep1 | GSE55347 | GPL17116 | Illumina HiSeq 2000 (Rattus norvegicus) | GSM1336244 | In Vivo Rat Liver Carcinogen (COH_WANG_AC0HK2ACXX_532_TGACCA_s_1) | S011127-131 |
| GSE47792 | GSE47875 | GPL1355 | [Rat230_2] Affymetrix Rat Genome 230 2.0 Array | GSM1161490 | Rat_Liver_CLOTRIMAZOLE_89mg/kg_5d_NU_OG_Rep2 | GSE55347 | GPL17116 | Illumina HiSeq 2000 (Rattus norvegicus) | GSM1336260 | In Vivo Rat Liver Carcinogen (COH_WANG_AC0HK2ACXX_548_CTTGTA_s_3) | S011127-132 |
| GSE47792 | GSE47875 | GPL1355 | [Rat230_2] Affymetrix Rat Genome 230 2.0 Array | GSM1161425 | Rat_Liver_Vehicle__5d_NN_OG_Rep1 | GSE55347 | GPL14844 | Illumina HiScanSQ (Rattus norvegicus) | GSM1336286 | In Vivo Rat Liver Carcinogen (COH_WANG_BC064YACXX_98141_L_CTL_NN_OG_CGATGT_s_4) | S011127-214 |
| GSE47792 | GSE47875 | GPL1355 | [Rat230_2] Affymetrix Rat Genome 230 2.0 Array | GSM1161436 | Rat_Liver_Vehicle__3d_NU_OG_Rep3 | GSE55347 | GPL14844 | Illumina HiScanSQ (Rattus norvegicus) | GSM1336285 | In Vivo Rat Liver Carcinogen (COH_WANG_BC064YACXX_97426_L_CTL_NU_OG_GCCAAT_s_3) | S011211-018 |
| GSE47792 | GSE47875 | GPL1355 | [Rat230_2] Affymetrix Rat Genome 230 2.0 Array | GSM1161424 | Rat_Liver_PHENOBARBITAL_54mg/kg_5d_NN_OG_Rep2 | GSE55347 | GPL17116 | Illumina HiSeq 2000 (Rattus norvegicus) | GSM1336219 | In Vivo Rat Liver Carcinogen (COH_WANG_AB029JACXX_97849_L_PHE_NN_OG_ACAGTG_s_3) | S020115-238 |
| GSE47792 | GSE47875 | GPL1355 | [Rat230_2] Affymetrix Rat Genome 230 2.0 Array | GSM1161423 | Rat_Liver_PHENOBARBITAL_54mg/kg_5d_NN_OG_Rep1 | GSE55347 | GPL17116 | Illumina HiSeq 2000 (Rattus norvegicus) | GSM1336184 | In Vivo Rat Liver Carcinogen (COH_WANG_AB0142ACXX_97922_L_PHE_NN_OG_ACAGTG_s_1) | S020115-239 |
| GSE47792 | GSE47875 | GPL1355 | [Rat230_2] Affymetrix Rat Genome 230 2.0 Array | GSM1161429 | Rat_Liver_PHENOBARBITAL_54mg/kg_5d_NN_OG_Rep3 | GSE55347 | GPL17116 | Illumina HiSeq 2000 (Rattus norvegicus) | GSM1336185 | In Vivo Rat Liver Carcinogen (COH_WANG_AB0142ACXX_97998_L_PHE_NN_OG_ACAGTG_s_2) | S020115-240 |
| GSE47792 | GSE47875 | GPL1355 | [Rat230_2] Affymetrix Rat Genome 230 2.0 Array | GSM1161503 | Rat_Liver_Vehicle__5d_NU_OG_Test_Rep2 | GSE55347 | GPL17116 | Illumina HiSeq 2000 (Rattus norvegicus) | GSM1336259 | In Vivo Rat Liver Carcinogen (COH_WANG_AC0HK2ACXX_547_CAGATC_s_3) | S020205-020 |
| GSE47792 | GSE47875 | GPL1355 | [Rat230_2] Affymetrix Rat | GSM1161480 | Rat_Liver_METHIMAZOLE_100mg/ | GSE55347 | GPL17116 | Illumina HiSeq 2000 (Rattus | GSM1336213 | In Vivo Rat Liver Carcinogen | S020205-283 |

| | | | | | | | | | | | |
|---|---|---|---|---|---|---|---|---|---|---|---|
| | | | Genome 230 2.0 Array | | kg_3d_NN_OG_Rep1 | | | norvegicus) | | (COH_WANG_AB029JACXX_50944_L_MET_NN_OG_GCCAAT_s_4) | |
| GSE47792 | GSE47875 | GPL1355 | [Rat230_2] Affymetrix Rat Genome 230 2.0 Array | GSM1161481 | Rat_Liver_METHIMAZOLE_100mg/kg_3d_NN_OG_Rep2 | GSE55347 | GPL17116 | Illumina HiSeq 2000 (Rattus norvegicus) | GSM1336214 | In Vivo Rat Liver Carcinogen (COH_WANG_AB029JACXX_50980_L_MET_NN_OG_GCCAAT_s_5) | S020205-284 |
| GSE47792 | GSE47875 | GPL1355 | [Rat230_2] Affymetrix Rat Genome 230 2.0 Array | GSM1161482 | Rat_Liver_METHIMAZOLE_100mg/kg_3d_NN_OG_Rep3 | GSE55347 | GPL17116 | Illumina HiSeq 2000 (Rattus norvegicus) | GSM1336215 | In Vivo Rat Liver Carcinogen (COH_WANG_AB029JACXX_51024_L_MET_NN_OG_GCCAAT_s_6) | S020205-285 |
| GSE47792 | GSE47875 | GPL1355 | [Rat230_2] Affymetrix Rat Genome 230 2.0 Array | GSM1161502 | Rat_Liver_CERIVASTATIN_7mg/kg_5d_NU_OG_Rep1 | GSE55347 | GPL17116 | Illumina HiSeq 2000 (Rattus norvegicus) | GSM1336251 | In Vivo Rat Liver Carcinogen (COH_WANG_AC0HK2ACXX_539_ACAGTG_s_2) | S020211-118 |
| GSE47792 | GSE47875 | GPL1355 | [Rat230_2] Affymetrix Rat Genome 230 2.0 Array | GSM1161504 | Rat_Liver_CERIVASTATIN_7mg/kg_5d_NU_OG_Rep2 | GSE55347 | GPL17116 | Illumina HiSeq 2000 (Rattus norvegicus) | GSM1336261 | In Vivo Rat Liver Carcinogen (COH_WANG_AC0HK2ACXX_555_CGATGT_s_5) | S020211-119 |
| GSE47792 | GSE47875 | GPL1355 | [Rat230_2] Affymetrix Rat Genome 230 2.0 Array | GSM1161505 | Rat_Liver_CERIVASTATIN_7mg/kg_5d_NU_OG_Rep3 | GSE55347 | GPL17116 | Illumina HiSeq 2000 (Rattus norvegicus) | GSM1336277 | In Vivo Rat Liver Carcinogen (COH_WANG_AC0HK2ACXX_571_CAGATC_s_7) | S020211-120 |
| GSE47792 | GSE47875 | GPL1355 | [Rat230_2] Affymetrix Rat Genome 230 2.0 Array | GSM1161508 | Rat_Liver_Vehicle__5d_NU_OG_Test_Rep4 | GSE55347 | GPL17116 | Illumina HiSeq 2000 (Rattus norvegicus) | GSM1336269 | In Vivo Rat Liver Carcinogen (COH_WANG_AC0HK2ACXX_563_ACAGTG_s_6) | S020429-019 |
| GSE47792 | GSE47875 | GPL1355 | [Rat230_2] Affymetrix Rat Genome 230 2.0 Array | GSM1161506 | Rat_Liver_Vehicle__5d_NU_OG_Test_Rep3 | GSE55347 | GPL17116 | Illumina HiSeq 2000 (Rattus norvegicus) | GSM1336248 | In Vivo Rat Liver Carcinogen (COH_WANG_AC0HK2ACXX_536_CTTGTA_s_1) | S020429-022 |
| GSE47792 | GSE47875 | GPL1355 | [Rat230_2] Affymetrix Rat Genome 230 2.0 Array | GSM1161501 | Rat_Liver_CLOFIBRIC-ACID_448mg/kg_5d_NU_OG_Rep3 | GSE55347 | GPL17116 | Illumina HiSeq 2000 (Rattus norvegicus) | GSM1336255 | In Vivo Rat Liver Carcinogen (COH_WANG_AC0HK2ACXX_543_CGATGT_s_3) | S020429-094 |
| GSE47792 | GSE47875 | GPL1355 | [Rat230_2] Affymetrix Rat Genome 230 2.0 | GSM1161499 | Rat_Liver_CLOFIBRIC-ACID_448mg/kg_5d_NU_OG_Re | GSE55347 | GPL17116 | Illumina HiSeq 2000 (Rattus norvegicus) | GSM1336265 | In Vivo Rat Liver Carcinogen (COH_WANG_AC0HK2 | S020429-095 |

| | | | | | | | | | | | |
|---|---|---|---|---|---|---|---|---|---|---|---|
| | | | Array | | p1 | | | | | ACXX_559_CAGATC_s_5) | |
| GSE47792 | GSE47875 | GPL1355 | [Rat230_2] Affymetrix Rat Genome 230 2.0 Array | GSM1161500 | Rat_Liver_CLOFIBRIC-ACID_448mg/kg_5d_NU_OG_Rep2 | GSE55347 | GPL17116 | Illumina HiSeq 2000 (Rattus norvegicus) | GSM1336281 | In Vivo Rat Liver Carcinogen (COH_WANG_AC0HK2ACXX_575_ACAGTG_s_8) | S020429-096 |
| GSE47792 | GSE47875 | GPL1355 | [Rat230_2] Affymetrix Rat Genome 230 2.0 Array | GSM1161454 | Rat_Liver_AFLATOXIN-B1_.3mg/kg_5d_NN_OG_Rep1 | GSE55347 | GPL17116 | Illumina HiSeq 2000 (Rattus norvegicus) | GSM1336201 | In Vivo Rat Liver Carcinogen (COH_WANG_AB0142ACXX_99320_L_AFL_NN_OG_ATCACG_s_1) | S020923-118 |
| GSE47792 | GSE47875 | GPL1355 | [Rat230_2] Affymetrix Rat Genome 230 2.0 Array | GSM1161454 | Rat_Liver_AFLATOXIN-B1_.3mg/kg_5d_NN_OG_Rep1 | GSE55347 | GPL17116 | Illumina HiSeq 2000 (Rattus norvegicus) | GSM1336258 | In Vivo Rat Liver Carcinogen (COH_WANG_AC0HK2ACXX_546_GCCAAT_s_3) | S020923-118 |
| GSE47792 | GSE47875 | GPL1355 | [Rat230_2] Affymetrix Rat Genome 230 2.0 Array | GSM1161454 | Rat_Liver_AFLATOXIN-B1_.3mg/kg_5d_NN_OG_Rep1 | GSE55347 | GPL14844 | Illumina HiScanSQ (Rattus norvegicus) | GSM1336295 | In Vivo Rat Liver Carcinogen (COH_WANG_BC064YACXX_99320_L_AFL_NN_OG_ATCACG_s_5) | S020923-118 |
| GSE47792 | GSE47875 | GPL1355 | [Rat230_2] Affymetrix Rat Genome 230 2.0 Array | GSM1161455 | Rat_Liver_AFLATOXIN-B1_.3mg/kg_5d_NN_OG_Rep2 | GSE55347 | GPL17116 | Illumina HiSeq 2000 (Rattus norvegicus) | GSM1336202 | In Vivo Rat Liver Carcinogen (COH_WANG_AB0142ACXX_99328_L_AFL_NN_OG_ATCACG_s_7) | S020923-119 |
| GSE47792 | GSE47875 | GPL1355 | [Rat230_2] Affymetrix Rat Genome 230 2.0 Array | GSM1161455 | Rat_Liver_AFLATOXIN-B1_.3mg/kg_5d_NN_OG_Rep2 | GSE55347 | GPL17116 | Illumina HiSeq 2000 (Rattus norvegicus) | GSM1336268 | In Vivo Rat Liver Carcinogen (COH_WANG_AC0HK2ACXX_562_TGACCA_s_6) | S020923-119 |
| GSE47792 | GSE47875 | GPL1355 | [Rat230_2] Affymetrix Rat Genome 230 2.0 Array | GSM1161455 | Rat_Liver_AFLATOXIN-B1_.3mg/kg_5d_NN_OG_Rep2 | GSE55347 | GPL14844 | Illumina HiScanSQ (Rattus norvegicus) | GSM1336296 | In Vivo Rat Liver Carcinogen (COH_WANG_BC064YACXX_99328_L_AFL_NN_OG_ATCACG_s_4) | S020923-119 |
| GSE47792 | GSE47875 | GPL1355 | [Rat230_2] Affymetrix Rat Genome 230 2.0 Array | GSM1161456 | Rat_Liver_AFLATOXIN-B1_.3mg/kg_5d_NN_OG_Rep3 | GSE55347 | GPL17116 | Illumina HiSeq 2000 (Rattus norvegicus) | GSM1336203 | In Vivo Rat Liver Carcinogen (COH_WANG_AB0142ACXX_99336_L_AFL_NN_OG_ATCACG_s_8) | S020923-120 |
| GSE47792 | GSE47875 | GPL1355 | [Rat230_2] Affymetrix Rat Genome 230 2.0 Array | GSM1161456 | Rat_Liver_AFLATOXIN-B1_.3mg/kg_5d_NN_OG_Rep3 | GSE55347 | GPL17116 | Illumina HiSeq 2000 (Rattus norvegicus) | GSM1336284 | In Vivo Rat Liver Carcinogen (COH_WANG_AC0HK2ACXX_578_CTTGTA_s | S020923-120 |

| | | | | | | | | | | |
|---|---|---|---|---|---|---|---|---|---|---|
| | | | | | | | | | _8) | |
| GSE47792 | GSE47875 | GPL1355 | [Rat230_2] Affymetrix Rat Genome 230 2.0 Array | GSM1161456 | Rat_Liver_AFLATOXIN-B1_.3mg/kg_5d_NN_OG_Rep3 | GSE55347 | GPL14844 | Illumina HiScanSQ (Rattus norvegicus) | GSM1336297 | In Vivo Rat Liver Carcinogen (COH_WANG_BC064YACXX_99336_L_AFL_NN_OG_ATCACG_s_2) | S020923-120 |
| GSE47792 | GSE47875 | GPL1355 | [Rat230_2] Affymetrix Rat Genome 230 2.0 Array | GSM1161475 | Rat_Liver_CHLOROFORM_600mg/kg_5d_NU_OG_Rep1 | GSE55347 | GPL17116 | Illumina HiSeq 2000 (Rattus norvegicus) | GSM1336210 | In Vivo Rat Liver Carcinogen (COH_WANG_AB0142ACXX_99837_L_CHL_NU_OG_CGATGT_s_4) | S030120-112 |
| GSE47792 | GSE47875 | GPL1355 | [Rat230_2] Affymetrix Rat Genome 230 2.0 Array | GSM1161476 | Rat_Liver_CHLOROFORM_600mg/kg_5d_NU_OG_Rep2 | GSE55347 | GPL17116 | Illumina HiSeq 2000 (Rattus norvegicus) | GSM1336211 | In Vivo Rat Liver Carcinogen (COH_WANG_AB0142ACXX_99844_L_CHL_NU_OG_CGATGT_s_5) | S030120-113 |
| GSE47792 | GSE47875 | GPL1355 | [Rat230_2] Affymetrix Rat Genome 230 2.0 Array | GSM1161477 | Rat_Liver_CHLOROFORM_600mg/kg_5d_NU_OG_Rep3 | GSE55347 | GPL17116 | Illumina HiSeq 2000 (Rattus norvegicus) | GSM1336212 | In Vivo Rat Liver Carcinogen (COH_WANG_AB0142ACXX_99846_L_CHL_NU_OG_CGATGT_s_6) | S030120-114 |
| GSE47792 | GSE47875 | GPL1355 | [Rat230_2] Affymetrix Rat Genome 230 2.0 Array | GSM1161432 | Rat_Liver_PIRINIXIC-ACID_364mg/kg_5d_NN_OG_Rep2 | GSE55347 | GPL17116 | Illumina HiSeq 2000 (Rattus norvegicus) | GSM1336187 | In Vivo Rat Liver Carcinogen (COH_WANG_AB0142ACXX_98455_L_PIR_NN_OG_GCCAAT_s_2) | S030218-120 |
| GSE47792 | GSE47875 | GPL1355 | [Rat230_2] Affymetrix Rat Genome 230 2.0 Array | GSM1161433 | Rat_Liver_PIRINIXIC-ACID_364mg/kg_5d_NN_OG_Rep3 | GSE55347 | GPL17116 | Illumina HiSeq 2000 (Rattus norvegicus) | GSM1336188 | In Vivo Rat Liver Carcinogen (COH_WANG_AB0142ACXX_98518_L_PIR_NN_OG_GCCAAT_s_3) | S030218-121 |
| GSE47792 | GSE47875 | GPL1355 | [Rat230_2] Affymetrix Rat Genome 230 2.0 Array | GSM1161427 | Rat_Liver_PIRINIXIC-ACID_364mg/kg_5d_NN_OG_Rep1 | GSE55347 | GPL17116 | Illumina HiSeq 2000 (Rattus norvegicus) | GSM1336186 | In Vivo Rat Liver Carcinogen (COH_WANG_AB0142ACXX_98379_L_PIR_NN_OG_GCCAAT_s_4) | S030218-122 |
| GSE47792 | GSE47875 | GPL1355 | [Rat230_2] Affymetrix Rat Genome 230 2.0 Array | GSM1161473 | Rat_Liver_Vehicle__3d_NU_OG_Rep4 | GSE55347 | GPL14844 | Illumina HiScanSQ (Rattus norvegicus) | GSM1336298 | In Vivo Rat Liver Carcinogen (COH_WANG_BC064YACXX_99787_L_CTL_NU_OG_CAGATC_s_3) | S040311-015 |
| GSE47792 | GSE47875 | GPL1355 | [Rat230_2] Affymetrix Rat Genome 230 2.0 Array | GSM1161512 | Rat_Liver_Vehicle__5d_NU_OG_Test_Rep5 | GSE55347 | GPL17116 | Illumina HiSeq 2000 (Rattus norvegicus) | GSM1336240 | In Vivo Rat Liver Carcinogen (COH_WANG_AC098LACXX_552_GCCAAT_s_7) | S040311-022 |

| | | | | | | | | | | | |
|---|---|---|---|---|---|---|---|---|---|---|---|
| GSE47792 | GSE47875 | GPL1355 | [Rat230_2] Affymetrix Rat Genome 230 2.0 Array | GSM1161460 | Rat_Liver_NAFENOPIN_338mg/kg_5d_NU_OG_Rep3 | GSE55347 | GPL17116 | Illumina HiSeq 2000 (Rattus norvegicus) | GSM1336200 | In Vivo Rat Liver Carcinogen (COH_WANG_AB0142ACXX_98911_L_NAF_NU_OG_GCCAAT_s_5) | S040311-096 |
| GSE47792 | GSE47875 | GPL1355 | [Rat230_2] Affymetrix Rat Genome 230 2.0 Array | GSM1161446 | Rat_Liver_NAFENOPIN_338mg/kg_5d_NU_OG_Rep1 | GSE55347 | GPL17116 | Illumina HiSeq 2000 (Rattus norvegicus) | GSM1336195 | In Vivo Rat Liver Carcinogen (COH_WANG_AB0142ACXX_98842_L_NAF_NU_OG_GCCAAT_s_6) | S040311-097 |
| GSE47792 | GSE47875 | GPL1355 | [Rat230_2] Affymetrix Rat Genome 230 2.0 Array | GSM1161447 | Rat_Liver_NAFENOPIN_338mg/kg_5d_NU_OG_Rep2 | GSE55347 | GPL17116 | Illumina HiSeq 2000 (Rattus norvegicus) | GSM1336198 | In Vivo Rat Liver Carcinogen (COH_WANG_AB0142ACXX_98864_L_NAF_NU_OG_GCCAAT_s_7) | S040311-098 |
| GSE47792 | GSE47875 | GPL1355 | [Rat230_2] Affymetrix Rat Genome 230 2.0 Array | GSM1161465 | Rat_Liver_Vehicle__1d_NN_IP_Rep2 | GSE55347 | GPL17116 | Illumina HiSeq 2000 (Rattus norvegicus) | GSM1336234 | In Vivo Rat Liver Carcinogen (COH_WANG_AB029JACXX_99596_L_CTL_NN_IP_GATCAG_s_1) | S041007-004 |
| GSE47792 | GSE47875 | GPL1355 | [Rat230_2] Affymetrix Rat Genome 230 2.0 Array | GSM1161466 | Rat_Liver_Vehicle__1d_NN_IP_Rep3 | GSE55347 | GPL17116 | Illumina HiSeq 2000 (Rattus norvegicus) | GSM1336235 | In Vivo Rat Liver Carcinogen (COH_WANG_AB029JACXX_99612_L_CTL_NN_IP_GATCAG_s_2) | S041007-005 |
| GSE47792 | GSE47875 | GPL1355 | [Rat230_2] Affymetrix Rat Genome 230 2.0 Array | GSM1161474 | Rat_Liver_Vehicle__3d_NU_OG_Rep5 | GSE55347 | GPL14844 | Illumina HiScanSQ (Rattus norvegicus) | GSM1336299 | In Vivo Rat Liver Carcinogen (COH_WANG_BC064YACXX_99810_L_CTL_NU_OG_ACAGTG_s_3) | XXV040217-005 |
| GSE74728 | GSE74676 | GPL1355 | [Rat230_2] Affymetrix Rat Genome 230 2.0 Array | GSM1929638 | Hippocampus_Control_rep1 | GSE74726 | GPL17116 | Illumina HiScanSQ (Rattus norvegicus) | GSM1931261 | Hippocampus_Control_mRNA1 | |
| GSE74728 | GSE74676 | GPL1355 | [Rat230_2] Affymetrix Rat Genome 230 2.0 Array | GSM1929639 | Hippocampus_Control_rep2 | GSE74726 | GPL17116 | Illumina HiScanSQ (Rattus norvegicus) | GSM1931262 | Hippocampus_Control_mRNA2 | |
| GSE74728 | GSE74676 | GPL1355 | [Rat230_2] Affymetrix Rat Genome 230 2.0 Array | GSM1929640 | Hippocampus_Control_rep3 | GSE74726 | GPL17116 | Illumina HiScanSQ (Rattus norvegicus) | GSM1931263 | Hippocampus_Control_mRNA3 | |
| GSE74728 | GSE74676 | GPL1355 | [Rat230_2] Affymetrix Rat Genome 230 2.0 Array | GSM1929641 | Hippocampus_Control_rep4 | GSE74726 | GPL17116 | Illumina HiScanSQ (Rattus norvegicus) | GSM1931264 | Hippocampus_Control_mRNA4 | |

| | | | | | | | | | | |
|---|---|---|---|---|---|---|---|---|---|---|
| GSE74728 | GSE74676 | GPL1355 | [Rat230_2] Affymetrix Rat Genome 230 2.0 Array | GSM1929642 | Hippocampus_Control_rep5 | GSE74726 | GPL17116 | Illumina HiScanSQ (Rattus norvegicus) | GSM1931265 | Hippocampus_Control_mRNA5 |
| GSE74728 | GSE74676 | GPL1355 | [Rat230_2] Affymetrix Rat Genome 230 2.0 Array | GSM1929643 | Hippocampus_Control_rep6 | GSE74726 | GPL17116 | Illumina HiScanSQ (Rattus norvegicus) | GSM1931266 | Hippocampus_Control_mRNA6 |
| GSE74728 | GSE74676 | GPL1355 | [Rat230_2] Affymetrix Rat Genome 230 2.0 Array | GSM1929644 | Hippocampus_Control_rep7 | GSE74726 | GPL17116 | Illumina HiScanSQ (Rattus norvegicus) | GSM1931267 | Hippocampus_Control_mRNA7 |
| GSE74728 | GSE74676 | GPL1355 | [Rat230_2] Affymetrix Rat Genome 230 2.0 Array | GSM1929658 | Hippocampus_High-CPF_rep1 | GSE74726 | GPL17116 | Illumina HiScanSQ (Rattus norvegicus) | GSM1931268 | Hippocampus_High-CPF_mRNA1 |
| GSE74728 | GSE74676 | GPL1355 | [Rat230_2] Affymetrix Rat Genome 230 2.0 Array | GSM1929659 | Hippocampus_High-CPF_rep2 | GSE74726 | GPL17116 | Illumina HiScanSQ (Rattus norvegicus) | GSM1931269 | Hippocampus_High-CPF_mRNA2 |
| GSE74728 | GSE74676 | GPL1355 | [Rat230_2] Affymetrix Rat Genome 230 2.0 Array | GSM1929660 | Hippocampus_High-CPF_rep3 | GSE74726 | GPL17116 | Illumina HiScanSQ (Rattus norvegicus) | GSM1931270 | Hippocampus_High-CPF_mRNA3 |
| GSE74728 | GSE74676 | GPL1355 | [Rat230_2] Affymetrix Rat Genome 230 2.0 Array | GSM1929661 | Hippocampus_High-CPF_rep4 | GSE74726 | GPL17116 | Illumina HiScanSQ (Rattus norvegicus) | GSM1931271 | Hippocampus_High-CPF_mRNA4 |
| GSE74728 | GSE74676 | GPL1355 | [Rat230_2] Affymetrix Rat Genome 230 2.0 Array | GSM1929662 | Hippocampus_High-CPF_rep5 | GSE74726 | GPL17116 | Illumina HiScanSQ (Rattus norvegicus) | GSM1931272 | Hippocampus_High-CPF_mRNA5 |
| GSE74728 | GSE74676 | GPL1355 | [Rat230_2] Affymetrix Rat Genome 230 2.0 Array | GSM1929663 | Hippocampus_High-CPF_rep6 | GSE74726 | GPL17116 | Illumina HiScanSQ (Rattus norvegicus) | GSM1931273 | Hippocampus_High-CPF_mRNA6 |
| | GSE122184 | GPL1355 | [Rat230_2] Affymetrix Rat Genome 230 2.0 Array | GSM3457372 | CMET11-104_1001: corn oil_5ml/kg_1001 | GSE122315 | GPL20084 | Illumina NextSeq 500 (Rattus norvegicus) | GSM3464007 | corn oil_5ml/kg_1001__RNA Seq |
| | GSE122184 | GPL1355 | [Rat230_2] Affymetrix Rat Genome 230 2.0 Array | GSM3457373 | CMET11-104_1003: corn oil_5ml/kg_1001 | GSE122315 | GPL20084 | Illumina NextSeq 500 (Rattus norvegicus) | GSM3464008 | corn oil_5ml/kg_1003_RNASeq |
| | GSE122184 | GPL1355 | [Rat230_2] Affymetrix Rat | GSM3457374 | CMET11-104_1005: corn | GSE122315 | GPL20084 | Illumina NextSeq 500 | GSM3464009 | corn oil_5ml/kg_1005_RNAS |

| | | | | | | | | | |
|---|---|---|---|---|---|---|---|---|---|
| | | Genome 230 2.0 Array [Rat230_2] | | oil_5ml/kg_1001 | | | (Rattus norvegicus) | | eq |
| GSE122184 | GPL1355 | Affymetrix Rat Genome 230 2.0 Array [Rat230_2] | GSM3457375 | CMET11-104_3001: ANIT_100mg/kg_3001 | GSE122315 | GPL20084 | Illumina NextSeq 500 (Rattus norvegicus) | GSM3464010 | ANIT_100mg/kg_3001_RNASeq |
| GSE122184 | GPL1355 | Affymetrix Rat Genome 230 2.0 Array [Rat230_2] | GSM3457376 | CMET11-104_3003: ANIT_100mg/kg_3003 | GSE122315 | GPL20084 | Illumina NextSeq 500 (Rattus norvegicus) | GSM3464011 | ANIT_100mg/kg_3003_RNASeq |
| GSE122184 | GPL1355 | Affymetrix Rat Genome 230 2.0 Array [Rat230_2] | GSM3457377 | CMET11-104_3005: ANIT_100mg/kg_3005 | GSE122315 | GPL20084 | Illumina NextSeq 500 (Rattus norvegicus) | GSM3464012 | ANIT_100mg/kg_3005_RNASeq |
| GSE122184 | GPL1355 | Affymetrix Rat Genome 230 2.0 Array [Rat230_2] | GSM3457378 | CMET11-104_4001: CCl4_1582mg/kg_4001 | GSE122315 | GPL20084 | Illumina NextSeq 500 (Rattus norvegicus) | GSM3464013 | CCl4_1582mg/kg_4001_RNASeq |
| GSE122184 | GPL1355 | Affymetrix Rat Genome 230 2.0 Array [Rat230_2] | GSM3457379 | CMET11-104_4003: CCl4_1582mg/kg_4003 | GSE122315 | GPL20084 | Illumina NextSeq 500 (Rattus norvegicus) | GSM3464014 | CCl4_1582mg/kg_4003_RNASeq |
| GSE122184 | GPL1355 | Affymetrix Rat Genome 230 2.0 Array [Rat230_2] | GSM3457380 | CMET11-104_4005: CCl4_1582mg/kg_4005 | GSE122315 | GPL20084 | Illumina NextSeq 500 (Rattus norvegicus) | GSM3464015 | CCl4_1582mg/kg_4005_RNASeq |
| GSE122184 | GPL1355 | Affymetrix Rat Genome 230 2.0 Array [Rat230_2] | GSM3457381 | ITP16-041_1001: water_10ml/kg_1001 | GSE122315 | GPL20084 | Illumina NextSeq 500 (Rattus norvegicus) | GSM3464016 | water_10ml/kg_1001_RNASeq |
| GSE122184 | GPL1355 | Affymetrix Rat Genome 230 2.0 Array [Rat230_2] | GSM3457382 | ITP16-041_1003: water_10ml/kg_1003 | GSE122315 | GPL20084 | Illumina NextSeq 500 (Rattus norvegicus) | GSM3464017 | water_10ml/kg_1003_RNASeq |
| GSE122184 | GPL1355 | Affymetrix Rat Genome 230 2.0 Array [Rat230_2] | GSM3457383 | ITP16-041_2001: corn oil_2ml/kg_2001 | GSE122315 | GPL20084 | Illumina NextSeq 500 (Rattus norvegicus) | GSM3464018 | corn oil_2ml/kg_2001_RNASeq |
| GSE122184 | GPL1355 | Affymetrix Rat Genome 230 2.0 Array [Rat230_2] | GSM3457384 | ITP16-041_2003: corn oil_2ml/kg_2003 | GSE122315 | GPL20084 | Illumina NextSeq 500 (Rattus norvegicus) | GSM3464019 | corn oil_2ml/kg_2003_RNASeq |
| GSE122184 | GPL1355 | Affymetrix Rat Genome 230 2.0 Array | GSM3457385 | ITP16-041_2005: corn oil_2ml/kg_2005 | GSE122315 | GPL20084 | Illumina NextSeq 500 (Rattus norvegicus) | GSM3464020 | corn oil_2ml/kg_2005_RNASeq |

| | | | | | | | | | |
|---|---|---|---|---|---|---|---|---|---|
| | | [Rat230_2] | | | | | | | |
| GSE122184 | GPL1355 | Affymetrix Rat Genome 230 2.0 Array [Rat230_2] | GSM3457386 | ITP16-041_3001: acetaminophen_1000mg/kg_3001 | GSE122315 | GPL20084 | Illumina NextSeq 500 (Rattus norvegicus) | GSM3464021 | acetaminophen_1000mg/kg_3001_RNASeq |
| GSE122184 | GPL1355 | Affymetrix Rat Genome 230 2.0 Array [Rat230_2] | GSM3457387 | ITP16-041_3003: acetaminophen_1000mg/kg_3003 | GSE122315 | GPL20084 | Illumina NextSeq 500 (Rattus norvegicus) | GSM3464022 | acetaminophen_1000mg/kg_3003_RNASeq |
| GSE122184 | GPL1355 | Affymetrix Rat Genome 230 2.0 Array [Rat230_2] | GSM3457388 | ITP16-041_3005: acetaminophen_1000mg/kg_3005 | GSE122315 | GPL20084 | Illumina NextSeq 500 (Rattus norvegicus) | GSM3464023 | acetaminophen_1000mg/kg_3005_RNASeq |
| GSE122184 | GPL1355 | Affymetrix Rat Genome 230 2.0 Array [Rat230_2] | GSM3457389 | ITP16-041_4001: diclofenac_10mg/kg_4001 | GSE122315 | GPL20084 | Illumina NextSeq 500 (Rattus norvegicus) | GSM3464024 | diclofenac_10mg/kg_4001_RNASeq |
| GSE122184 | GPL1355 | Affymetrix Rat Genome 230 2.0 Array [Rat230_2] | GSM3457390 | ITP16-041_4003: diclofenac_10mg/kg_4003 | GSE122315 | GPL20084 | Illumina NextSeq 500 (Rattus norvegicus) | GSM3464025 | diclofenac_10mg/kg_4003_RNASeq |
| GSE122184 | GPL1355 | Affymetrix Rat Genome 230 2.0 Array [Rat230_2] | GSM3457391 | ITP16-041_4005: diclofenac_10mg/kg_4005 | GSE122315 | GPL20084 | Illumina NextSeq 500 (Rattus norvegicus) | GSM3464026 | diclofenac_10mg/kg_4005_RNASeq |
| GSE122184 | GPL1355 | Affymetrix Rat Genome 230 2.0 Array [Rat230_2] | GSM3457392 | ITP16-090_1001: 35% ethanol_2ml/kg_1001 | GSE122315 | GPL20084 | Illumina NextSeq 500 (Rattus norvegicus) | GSM3464027 | 35% ethanol_2ml/kg_1001_RNASeq |
| GSE122184 | GPL1355 | Affymetrix Rat Genome 230 2.0 Array [Rat230_2] | GSM3457393 | ITP16-090_1003: 35% ethanol_2ml/kg_1003 | GSE122315 | GPL20084 | Illumina NextSeq 500 (Rattus norvegicus) | GSM3464028 | 35% ethanol_2ml/kg_1003_RNASeq |
| GSE122184 | GPL1355 | Affymetrix Rat Genome 230 2.0 Array [Rat230_2] | GSM3457394 | ITP16-090_1005: 35% ethanol_2ml/kg_1005 | GSE122315 | GPL20084 | Illumina NextSeq 500 (Rattus norvegicus) | GSM3464029 | 35% ethanol_2ml/kg_1005_RNASeq |
| GSE122184 | GPL1355 | Affymetrix Rat Genome 230 2.0 Array [Rat230_2] | GSM3457395 | ITP16-090_4001: methylenedianiline_100mg/kg_4001 | GSE122315 | GPL20084 | Illumina NextSeq 500 (Rattus norvegicus) | GSM3464030 | methylenedianiline_100mg/kg_4001_RNASeq |
| GSE122184 | GPL1355 | Affymetrix Rat Genome 230 2.0 Array [Rat230_2] | GSM3457396 | ITP16-090_4003: methylenedianiline_100mg/kg_4003 | GSE122315 | GPL20084 | Illumina NextSeq 500 (Rattus norvegicus) | GSM3464031 | methylenedianiline_100mg/kg_4003_RNASeq |
| GSE122184 | GPL1355 | Affymetrix Rat | GSM3457397 | ITP16-090_4005: methylenedianiline_ | GSE122315 | GPL20084 | Illumina NextSeq 500 | GSM3464032 | methylenedianiline_100mg/kg_4005_RNASeq |

| | | | | | | | | |
|---|---|---|---|---|---|---|---|---|
| | | | Genome 230 2.0 Array | | 100mg/kg_4005 | | | (Rattus norvegicus) |

| B | array | | | | | seq | | | | | |
|---|---|---|---|---|---|---|---|---|---|---|---|
| superseries | series | platform | platform desc. | sample | sample desc. | series | platform | platform desc. | sample | sample desc. | sample code |
| | GSE66419 | GPL198 | [ATH1-121501] Affymetrix Arabidopsis ATH1 Genome Array | GSM1622277 | Ler_rep1 | GSE57690 | GPL13222 | Illumina HiSeq 2000 (Arabidopsis thaliana) | GSM1386775 | Ler - sample 1 - long fraction | |
| | GSE66419 | GPL198 | [ATH1-121501] Affymetrix Arabidopsis ATH1 Genome Array | GSM1622278 | Ler_rep2 | GSE57690 | GPL13222 | Illumina HiSeq 2000 (Arabidopsis thaliana) | GSM1386775 | Ler - sample 1 - long fraction | |
| | GSE66419 | GPL198 | [ATH1-121501] Affymetrix Arabidopsis ATH1 Genome Array | GSM1622279 | Ler_rep3 | GSE57690 | GPL13222 | Illumina HiSeq 2000 (Arabidopsis thaliana) | GSM1386775 | Ler - sample 1 - long fraction | |
| | GSE66419 | GPL198 | [ATH1-121501] Affymetrix Arabidopsis ATH1 Genome Array | GSM1622278 | Ler_rep2 | GSE57690 | GPL13222 | Illumina HiSeq 2000 (Arabidopsis thaliana) | GSM1386776 | Ler - sample 1 -short fraction | |
| | GSE66419 | GPL198 | [ATH1-121501] Affymetrix Arabidopsis ATH1 Genome Array | GSM1622279 | Ler_rep3 | GSE57690 | GPL13222 | Illumina HiSeq 2000 (Arabidopsis thaliana) | GSM1386776 | Ler - sample 1 -short fraction | |
| | GSE66419 | GPL198 | [ATH1-121501] Affymetrix Arabidopsis ATH1 Genome Array | GSM1622277 | Ler_rep1 | GSE57690 | GPL13222 | Illumina HiSeq 2000 (Arabidopsis thaliana) | GSM1386776 | Ler - sample 1 -short fraction | |
| | GSE66419 | GPL198 | [ATH1-121501] Affymetrix Arabidopsis ATH1 Genome Array | GSM1622277 | Ler_rep1 | GSE57690 | GPL13222 | Illumina HiSeq 2000 (Arabidopsis thaliana) | GSM1386777 | Ler - sample 2 - long fraction | |
| | GSE66419 | GPL198 | [ATH1-121501] | GSM1622278 | Ler_rep2 | GSE57690 | GPL13222 | Illumina HiSeq | GSM1386777 | Ler - sample 2 - long | |

| | | | | | | | | | |
|---|---|---|---|---|---|---|---|---|---|
| | | Affymetrix Arabidopsis ATH1 Genome Array [ATH1-121501] | | | | | 2000 (Arabidopsis thaliana) | | fraction |
| GSE66419 | GPL198 | Affymetrix Arabidopsis ATH1 Genome Array [ATH1-121501] | GSM1622279 | Ler_rep3 | GSE57690 | GPL13222 | Illumina HiSeq 2000 (Arabidopsis thaliana) | GSM1386777 | Ler - sample 2 - long fraction |
| GSE66419 | GPL198 | Affymetrix Arabidopsis ATH1 Genome Array [ATH1-121501] | GSM1622277 | Ler_rep1 | GSE57690 | GPL13222 | Illumina HiSeq 2000 (Arabidopsis thaliana) | GSM1386778 | Ler - sample 2 -short fraction |
| GSE66419 | GPL198 | Affymetrix Arabidopsis ATH1 Genome Array [ATH1-121501] | GSM1622278 | Ler_rep2 | GSE57690 | GPL13222 | Illumina HiSeq 2000 (Arabidopsis thaliana) | GSM1386778 | Ler - sample 2 -short fraction |
| GSE66419 | GPL198 | Affymetrix Arabidopsis ATH1 Genome Array [ATH1-121501] | GSM1622279 | Ler_rep3 | GSE57690 | GPL13222 | Illumina HiSeq 2000 (Arabidopsis thaliana) | GSM1386778 | Ler - sample 2 -short fraction |
| GSE66419 | GPL198 | Affymetrix Arabidopsis ATH1 Genome Array [ATH1-121501] | GSM1622277 | Ler_rep1 | GSE57690 | GPL13222 | Illumina HiSeq 2000 (Arabidopsis thaliana) | GSM1386779 | Ler - sample 3 - long fraction |
| GSE66419 | GPL198 | Affymetrix Arabidopsis ATH1 Genome Array [ATH1-121501] | GSM1622278 | Ler_rep2 | GSE57690 | GPL13222 | Illumina HiSeq 2000 (Arabidopsis thaliana) | GSM1386779 | Ler - sample 3 - long fraction |
| GSE66419 | GPL198 | Affymetrix Arabidopsis ATH1 Genome Array [ATH1-121501] | GSM1622279 | Ler_rep3 | GSE57690 | GPL13222 | Illumina HiSeq 2000 (Arabidopsis thaliana) | GSM1386779 | Ler - sample 3 - long fraction |
| GSE66419 | GPL198 | Affymetrix Arabidopsis ATH1 Genome Array [ATH1-121501] | GSM1622278 | Ler_rep2 | GSE57690 | GPL13222 | Illumina HiSeq 2000 (Arabidopsis thaliana) | GSM1386780 | Ler - sample 3 -short fraction |
| GSE66419 | GPL198 | Affymetrix | GSM1622279 | Ler_rep3 | GSE57690 | GPL13222 | Illumina HiSeq 2000 | GSM1386780 | Ler - sample 3 -short fraction |

| | | | | | | | | | | |
|---|---|---|---|---|---|---|---|---|---|---|
| | | | Arabidopsis ATH1 Genome Array [ATH1-121501] | | | | | (Arabidopsis thaliana) | | |
| | GSE66419 | GPL198 | Affymetrix Arabidopsis ATH1 Genome Array [ATH1-121501] | GSM1622277 | Ler_rep1 | GSE57690 | GPL13222 | Illumina HiSeq 2000 (Arabidopsis thaliana) | GSM1386780 | Ler - sample 3 -short fraction |
| | GSE66419 | GPL198 | Affymetrix Arabidopsis ATH1 Genome Array [ATH1-121501] | GSM1622277 | Ler_rep1 | GSE57690 | GPL13222 | Illumina HiSeq 2000 (Arabidopsis thaliana) | GSM1386781 | Ler - sample 4 - long fraction |
| | GSE66419 | GPL198 | Affymetrix Arabidopsis ATH1 Genome Array [ATH1-121501] | GSM1622278 | Ler_rep2 | GSE57690 | GPL13222 | Illumina HiSeq 2000 (Arabidopsis thaliana) | GSM1386781 | Ler - sample 4 - long fraction |
| | GSE66419 | GPL198 | Affymetrix Arabidopsis ATH1 Genome Array [ATH1-121501] | GSM1622279 | Ler_rep3 | GSE57690 | GPL13222 | Illumina HiSeq 2000 (Arabidopsis thaliana) | GSM1386781 | Ler - sample 4 - long fraction |
| | GSE66419 | GPL198 | Affymetrix Arabidopsis ATH1 Genome Array [ATH1-121501] | GSM1622277 | Ler_rep1 | GSE57690 | GPL13222 | Illumina HiSeq 2000 (Arabidopsis thaliana) | GSM1386782 | Ler - sample 4 -short fraction |
| | GSE66419 | GPL198 | Affymetrix Arabidopsis ATH1 Genome Array [ATH1-121501] | GSM1622278 | Ler_rep2 | GSE57690 | GPL13222 | Illumina HiSeq 2000 (Arabidopsis thaliana) | GSM1386782 | Ler - sample 4 -short fraction |
| | GSE66419 | GPL198 | Affymetrix Arabidopsis ATH1 Genome Array [ATH1-121501] | GSM1622279 | Ler_rep3 | GSE57690 | GPL13222 | Illumina HiSeq 2000 (Arabidopsis thaliana) | GSM1386782 | Ler - sample 4 -short fraction |
| GSE58857 | GSE58855 | GPL198 | Affymetrix Arabidopsis ATH1 Genome Array [ATH1-121501] | GSM1420938 | ATH1 - ML1p::YFP-RCI2A (ML1Y) - replicate 1 | GSE58856 | GPL13222 | Illumina HiSeq 2000 (Arabidopsis thaliana) | GSM1420953 | RNASeq - ML1p::YFP-RCI2A (ML1Y) - replicate 1 |
| GSE58857 | GSE58855 | GPL198 | Affymetrix Arabidopsis | GSM1420939 | ATH1 - ML1p::YFP-RCI2A (ML1Y) - replicate 2 | GSE58856 | GPL13222 | Illumina HiSeq 2000 (Arabidopsis | GSM1420953 | RNASeq - ML1p::YFP-RCI2A (ML1Y) - replicate 1 |

| | | | | | | | | | | |
|---|---|---|---|---|---|---|---|---|---|---|
| | | | ATH1 Genome Array [ATH1-121501] | | | | | thaliana) | | |
| GSE58857 | GSE58855 | GPL198 | Affymetrix Arabidopsis ATH1 Genome Array [ATH1-121501] | GSM1420940 | ATH1 - ML1p::YFP-RCI2A (ML1Y) - replicate 3 | GSE58856 | GPL13222 | Illumina HiSeq 2000 (Arabidopsis thaliana) | GSM1420953 | RNASeq - ML1p::YFP-RCI2A (ML1Y) - replicate 1 |
| GSE58857 | GSE58855 | GPL198 | Affymetrix Arabidopsis ATH1 Genome Array [ATH1-121501] | GSM1420939 | ATH1 - ML1p::YFP-RCI2A (ML1Y) - replicate 2 | GSE58856 | GPL13222 | Illumina HiSeq 2000 (Arabidopsis thaliana) | GSM1420954 | RNASeq - ML1p::YFP-RCI2A (ML1Y) - replicate 2 |
| GSE58857 | GSE58855 | GPL198 | Affymetrix Arabidopsis ATH1 Genome Array [ATH1-121501] | GSM1420940 | ATH1 - ML1p::YFP-RCI2A (ML1Y) - replicate 3 | GSE58856 | GPL13222 | Illumina HiSeq 2000 (Arabidopsis thaliana) | GSM1420954 | RNASeq - ML1p::YFP-RCI2A (ML1Y) - replicate 2 |
| GSE58857 | GSE58855 | GPL198 | Affymetrix Arabidopsis ATH1 Genome Array [ATH1-121501] | GSM1420938 | ATH1 - ML1p::YFP-RCI2A (ML1Y) - replicate 1 | GSE58856 | GPL13222 | Illumina HiSeq 2000 (Arabidopsis thaliana) | GSM1420954 | RNASeq - ML1p::YFP-RCI2A (ML1Y) - replicate 2 |
| GSE58857 | GSE58855 | GPL198 | Affymetrix Arabidopsis ATH1 Genome Array [ATH1-121501] | GSM1420941 | ATH1 - SPCHp::SPCH-YFP (SSY) - replicate 1 | GSE58856 | GPL13222 | Illumina HiSeq 2000 (Arabidopsis thaliana) | GSM1420955 | RNASeq - SPCHp::SPCH-YFP (SSY) - replicate 1 |
| GSE58857 | GSE58855 | GPL198 | Affymetrix Arabidopsis ATH1 Genome Array [ATH1-121501] | GSM1420942 | ATH1 - SPCHp::SPCH-YFP (SSY) - replicate 2 | GSE58856 | GPL13222 | Illumina HiSeq 2000 (Arabidopsis thaliana) | GSM1420955 | RNASeq - SPCHp::SPCH-YFP (SSY) - replicate 1 |
| GSE58857 | GSE58855 | GPL198 | Affymetrix Arabidopsis ATH1 Genome Array [ATH1-121501] | GSM1420943 | ATH1 - SPCHp::SPCH-YFP (SSY) - replicate 3 | GSE58856 | GPL13222 | Illumina HiSeq 2000 (Arabidopsis thaliana) | GSM1420955 | RNASeq - SPCHp::SPCH-YFP (SSY) - replicate 1 |
| GSE58857 | GSE58855 | GPL198 | Affymetrix Arabidopsis ATH1 Genome Array [ATH1-121501] | GSM1420941 | ATH1 - SPCHp::SPCH-YFP (SSY) - replicate 1 | GSE58856 | GPL13222 | Illumina HiSeq 2000 (Arabidopsis thaliana) | GSM1420956 | RNASeq - SPCHp::SPCH-YFP (SSY) - replicate 2 |
| GSE58857 | GSE58855 | GPL198 | Affymetrix Arabidopsis ATH1 Genome | GSM1420942 | ATH1 - SPCHp::SPCH-YFP (SSY) - replicate 2 | GSE58856 | GPL13222 | Illumina HiSeq 2000 (Arabidopsis thaliana) | GSM1420956 | RNASeq - SPCHp::SPCH-YFP (SSY) - replicate 2 |

| | | | | | | | | | | |
|---|---|---|---|---|---|---|---|---|---|---|
| GSE58857 | GSE58855 | GPL198 | Array [ATH1-121501] Affymetrix Arabidopsis ATH1 Genome Array | GSM1420943 | ATH1 - SPCHp::SPCH-YFP (SSY) - replicate 3 | GSE58856 | GPL13222 | Illumina HiSeq 2000 (Arabidopsis thaliana) | GSM1420956 | RNASeq - SPCHp::SPCH-YFP (SSY) - replicate 2 |
| GSE58857 | GSE58855 | GPL198 | [ATH1-121501] Affymetrix Arabidopsis ATH1 Genome Array | GSM1420941 | ATH1 - SPCHp::SPCH-YFP (SSY) - replicate 1 | GSE58856 | GPL13222 | Illumina HiSeq 2000 (Arabidopsis thaliana) | GSM1420957 | RNASeq - SPCHp::SPCH-YFP (SSY) - replicate 3 |
| GSE58857 | GSE58855 | GPL198 | [ATH1-121501] Affymetrix Arabidopsis ATH1 Genome Array | GSM1420942 | ATH1 - SPCHp::SPCH-YFP (SSY) - replicate 2 | GSE58856 | GPL13222 | Illumina HiSeq 2000 (Arabidopsis thaliana) | GSM1420957 | RNASeq - SPCHp::SPCH-YFP (SSY) - replicate 3 |
| GSE58857 | GSE58855 | GPL198 | [ATH1-121501] Affymetrix Arabidopsis ATH1 Genome Array | GSM1420943 | ATH1 - SPCHp::SPCH-YFP (SSY) - replicate 3 | GSE58856 | GPL13222 | Illumina HiSeq 2000 (Arabidopsis thaliana) | GSM1420957 | RNASeq - SPCHp::SPCH-YFP (SSY) - replicate 3 |
| GSE58857 | GSE58855 | GPL198 | [ATH1-121501] Affymetrix Arabidopsis ATH1 Genome Array | GSM1420945 | ATH1 - MUTEp::nucGFP (MG) - replicate 2 | GSE58856 | GPL13222 | Illumina HiSeq 2000 (Arabidopsis thaliana) | GSM1420958 | RNASeq - MUTEp::nucGFP (MG) - replicate 1 |
| GSE58857 | GSE58855 | GPL198 | [ATH1-121501] Affymetrix Arabidopsis ATH1 Genome Array | GSM1420946 | ATH1 - MUTEp::nucGFP (MG) - replicate 3 | GSE58856 | GPL13222 | Illumina HiSeq 2000 (Arabidopsis thaliana) | GSM1420958 | RNASeq - MUTEp::nucGFP (MG) - replicate 1 |
| GSE58857 | GSE58855 | GPL198 | [ATH1-121501] Affymetrix Arabidopsis ATH1 Genome Array | GSM1420944 | ATH1 - MUTEp::nucGFP (MG) - replicate 1 | GSE58856 | GPL13222 | Illumina HiSeq 2000 (Arabidopsis thaliana) | GSM1420958 | RNASeq - MUTEp::nucGFP (MG) - replicate 1 |
| GSE58857 | GSE58855 | GPL198 | [ATH1-121501] Affymetrix Arabidopsis ATH1 Genome Array | GSM1420944 | ATH1 - MUTEp::nucGFP (MG) - replicate 1 | GSE58856 | GPL13222 | Illumina HiSeq 2000 (Arabidopsis thaliana) | GSM1420959 | RNASeq - MUTEp::nucGFP (MG) - replicate 2 |
| GSE58857 | GSE58855 | GPL198 | [ATH1-121501] Affymetrix Arabidopsis ATH1 Genome Array | GSM1420945 | ATH1 - MUTEp::nucGFP (MG) - replicate 2 | GSE58856 | GPL13222 | Illumina HiSeq 2000 (Arabidopsis thaliana) | GSM1420959 | RNASeq - MUTEp::nucGFP (MG) - replicate 2 |

| | | | | | | | | | |
|---|---|---|---|---|---|---|---|---|---|
| GSE58857 | GSE58855 | GPL198 | [ATH1-121501] Affymetrix Arabidopsis ATH1 Genome Array | GSM1420946 | ATH1 - MUTEp::nucGFP (MG) - replicate 3 | GSE58856 | GPL13222 | Illumina HiSeq 2000 (Arabidopsis thaliana) | GSM1420959 | RNASeq - MUTEp::nucGFP (MG) - replicate 2 |
| GSE58857 | GSE58855 | GPL198 | [ATH1-121501] Affymetrix Arabidopsis ATH1 Genome Array | GSM1420947 | ATH1 - FAMAp::GFP-FAMA (FGF) - replicate 1 | GSE58856 | GPL13222 | Illumina HiSeq 2000 (Arabidopsis thaliana) | GSM1420960 | RNASeq - FAMAp::GFP-FAMA (FGF) - replicate 1 |
| GSE58857 | GSE58855 | GPL198 | [ATH1-121501] Affymetrix Arabidopsis ATH1 Genome Array | GSM1420948 | ATH1 - FAMAp::GFP-FAMA (FGF) - replicate 2 | GSE58856 | GPL13222 | Illumina HiSeq 2000 (Arabidopsis thaliana) | GSM1420960 | RNASeq - FAMAp::GFP-FAMA (FGF) - replicate 1 |
| GSE58857 | GSE58855 | GPL198 | [ATH1-121501] Affymetrix Arabidopsis ATH1 Genome Array | GSM1420949 | ATH1 - FAMAp::GFP-FAMA (FGF) - replicate 3 | GSE58856 | GPL13222 | Illumina HiSeq 2000 (Arabidopsis thaliana) | GSM1420960 | RNASeq - FAMAp::GFP-FAMA (FGF) - replicate 1 |
| GSE58857 | GSE58855 | GPL198 | [ATH1-121501] Affymetrix Arabidopsis ATH1 Genome Array | GSM1420948 | ATH1 - FAMAp::GFP-FAMA (FGF) - replicate 2 | GSE58856 | GPL13222 | Illumina HiSeq 2000 (Arabidopsis thaliana) | GSM1420961 | RNASeq - FAMAp::GFP-FAMA (FGF) - replicate 2 |
| GSE58857 | GSE58855 | GPL198 | [ATH1-121501] Affymetrix Arabidopsis ATH1 Genome Array | GSM1420949 | ATH1 - FAMAp::GFP-FAMA (FGF) - replicate 3 | GSE58856 | GPL13222 | Illumina HiSeq 2000 (Arabidopsis thaliana) | GSM1420961 | RNASeq - FAMAp::GFP-FAMA (FGF) - replicate 2 |
| GSE58857 | GSE58855 | GPL198 | [ATH1-121501] Affymetrix Arabidopsis ATH1 Genome Array | GSM1420947 | ATH1 - FAMAp::GFP-FAMA (FGF) - replicate 1 | GSE58856 | GPL13222 | Illumina HiSeq 2000 (Arabidopsis thaliana) | GSM1420961 | RNASeq - FAMAp::GFP-FAMA (FGF) - replicate 2 |
| GSE58857 | GSE58855 | GPL198 | [ATH1-121501] Affymetrix Arabidopsis ATH1 Genome Array | GSM1420950 | ATH1 - E1728::GFP (E1728G) - replicate 1 | GSE58856 | GPL13222 | Illumina HiSeq 2000 (Arabidopsis thaliana) | GSM1420962 | RNASeq - E1728::GFP (E1728G) - replicate 1 |
| GSE58857 | GSE58855 | GPL198 | [ATH1-121501] Affymetrix Arabidopsis ATH1 Genome Array | GSM1420951 | ATH1 - E1728::GFP (E1728G) - replicate 2 | GSE58856 | GPL13222 | Illumina HiSeq 2000 (Arabidopsis thaliana) | GSM1420962 | RNASeq - E1728::GFP (E1728G) - replicate 1 |
| GSE58857 | GSE58855 | GPL198 | [ATH1-121501] | GSM1420952 | ATH1 - | GSE58856 | GPL13222 | Illumina HiSeq | GSM1420962 | RNASeq - E1728::GFP |

| | | | | | | | | | | |
|---|---|---|---|---|---|---|---|---|---|---|
| | | | Affymetrix Arabidopsis ATH1 Genome Array [ATH1-121501] | | E1728::GFP (E1728G) - replicate 3 | | | 2000 (Arabidopsis thaliana) | | (E1728G) - replicate 1 |
| GSE58857 | GSE58855 | GPL198 | Affymetrix Arabidopsis ATH1 Genome Array [ATH1-121501] | GSM1420950 | ATH1 - E1728::GFP (E1728G) - replicate 1 | GSE58856 | GPL13222 | Illumina HiSeq 2000 (Arabidopsis thaliana) | GSM1420963 | RNASeq - E1728::GFP (E1728G) - replicate 2 |
| GSE58857 | GSE58855 | GPL198 | Affymetrix Arabidopsis ATH1 Genome Array [ATH1-121501] | GSM1420952 | ATH1 - E1728::GFP (E1728G) - replicate 3 | GSE58856 | GPL13222 | Illumina HiSeq 2000 (Arabidopsis thaliana) | GSM1420963 | RNASeq - E1728::GFP (E1728G) - replicate 2 |
| GSE58857 | GSE58855 | GPL198 | Affymetrix Arabidopsis ATH1 Genome Array [ATH1-121501] | GSM1420951 | ATH1 - E1728::GFP (E1728G) - replicate 2 | GSE58856 | GPL13222 | Illumina HiSeq 2000 (Arabidopsis thaliana) | GSM1420963 | RNASeq - E1728::GFP (E1728G) - replicate 2 |
| | GSE95396 | GPL198 | Affymetrix Arabidopsis ATH1 Genome Array [ATH1-121501] | GSM2509756 | Col0_time0_rep1 | GSE95473 | GPL13222 | Illumina HiSeq 2000 (Arabidopsis thaliana) | GSM2514737 | Col0_small_RNA_rep1 |
| | GSE95396 | GPL198 | Affymetrix Arabidopsis ATH1 Genome Array [ATH1-121501] | GSM2509757 | Col0_time0_rep2 | GSE95473 | GPL13222 | Illumina HiSeq 2000 (Arabidopsis thaliana) | GSM2514737 | Col0_small_RNA_rep1 |
| | GSE95396 | GPL198 | Affymetrix Arabidopsis ATH1 Genome Array [ATH1-121501] | GSM2509758 | Col0_time0_rep3 | GSE95473 | GPL13222 | Illumina HiSeq 2000 (Arabidopsis thaliana) | GSM2514737 | Col0_small_RNA_rep1 |
| | GSE95396 | GPL198 | Affymetrix Arabidopsis ATH1 Genome Array [ATH1-121501] | GSM2509756 | Col0_time0_rep1 | GSE95473 | GPL13222 | Illumina HiSeq 2000 (Arabidopsis thaliana) | GSM2514738 | Col0_small_RNA_rep2 |
| | GSE95396 | GPL198 | Affymetrix Arabidopsis ATH1 Genome Array [ATH1-121501] | GSM2509757 | Col0_time0_rep2 | GSE95473 | GPL13222 | Illumina HiSeq 2000 (Arabidopsis thaliana) | GSM2514738 | Col0_small_RNA_rep2 |
| | GSE95396 | GPL198 | Affymetrix | GSM2509758 | Col0_time0_rep3 | GSE95473 | GPL13222 | Illumina HiSeq 2000 | GSM2514738 | Col0_small_RNA_rep2 |

| | | | | | | | | | |
|---|---|---|---|---|---|---|---|---|---|
| | | Arabidopsis ATH1 Genome Array [ATH1-121501] | | | | | (Arabidopsis thaliana) | | |
| GSE95396 | GPL198 | Affymetrix Arabidopsis ATH1 Genome Array [ATH1-121501] | GSM2509756 | Col0_time0_rep1 | GSE95473 | GPL13222 | Illumina HiSeq 2000 (Arabidopsis thaliana) | GSM2514739 | Col0_small_RNA_rep3 |
| GSE95396 | GPL198 | Affymetrix Arabidopsis ATH1 Genome Array [ATH1-121501] | GSM2509757 | Col0_time0_rep2 | GSE95473 | GPL13222 | Illumina HiSeq 2000 (Arabidopsis thaliana) | GSM2514739 | Col0_small_RNA_rep3 |
| GSE95396 | GPL198 | Affymetrix Arabidopsis ATH1 Genome Array [ATH1-121501] | GSM2509758 | Col0_time0_rep3 | GSE95473 | GPL13222 | Illumina HiSeq 2000 (Arabidopsis thaliana) | GSM2514739 | Col0_small_RNA_rep3 |
| GSE95396 | GPL198 | Affymetrix Arabidopsis ATH1 Genome Array [ATH1-121501] | GSM2509762 | xrn3-8_time0_rep1 | GSE95473 | GPL13222 | Illumina HiSeq 2000 (Arabidopsis thaliana) | GSM2514740 | xrn3-8_small_RNA_rep1 |
| GSE95396 | GPL198 | Affymetrix Arabidopsis ATH1 Genome Array [ATH1-121501] | GSM2509763 | xrn3-8_time0_rep2 | GSE95473 | GPL13222 | Illumina HiSeq 2000 (Arabidopsis thaliana) | GSM2514740 | xrn3-8_small_RNA_rep1 |
| GSE95396 | GPL198 | Affymetrix Arabidopsis ATH1 Genome Array [ATH1-121501] | GSM2509764 | xrn3-8_time0_rep3 | GSE95473 | GPL13222 | Illumina HiSeq 2000 (Arabidopsis thaliana) | GSM2514740 | xrn3-8_small_RNA_rep1 |
| GSE95396 | GPL198 | Affymetrix Arabidopsis ATH1 Genome Array [ATH1-121501] | GSM2509762 | xrn3-8_time0_rep1 | GSE95473 | GPL13222 | Illumina HiSeq 2000 (Arabidopsis thaliana) | GSM2514741 | xrn3-8_small_RNA_rep2 |
| GSE95396 | GPL198 | Affymetrix Arabidopsis ATH1 Genome Array [ATH1-121501] | GSM2509763 | xrn3-8_time0_rep2 | GSE95473 | GPL13222 | Illumina HiSeq 2000 (Arabidopsis thaliana) | GSM2514741 | xrn3-8_small_RNA_rep2 |
| GSE95396 | GPL198 | Affymetrix Arabidopsis | GSM2509764 | xrn3-8_time0_rep3 | GSE95473 | GPL13222 | Illumina HiSeq 2000 (Arabidopsis | GSM2514741 | xrn3-8_small_RNA_rep2 |

| | | | | | | | | | | |
|---|---|---|---|---|---|---|---|---|---|---|
| | | | ATH1 Genome Array [ATH1-121501] | | | | | thaliana) | | |
| | GSE95396 | GPL198 | Affymetrix Arabidopsis ATH1 Genome Array [ATH1-121501] | GSM2509763 | xrn3-8_time0_rep2 | GSE95473 | GPL13222 | Illumina HiSeq 2000 (Arabidopsis thaliana) | GSM2514742 | xrn3-8_small_RNA_rep3 |
| | GSE95396 | GPL198 | Affymetrix Arabidopsis ATH1 Genome Array [ATH1-121501] | GSM2509762 | xrn3-8_time0_rep1 | GSE95473 | GPL13222 | Illumina HiSeq 2000 (Arabidopsis thaliana) | GSM2514742 | xrn3-8_small_RNA_rep3 |
| | GSE95396 | GPL198 | Affymetrix Arabidopsis ATH1 Genome Array [ATH1-121501] | GSM2509764 | xrn3-8_time0_rep3 | GSE95473 | GPL13222 | Illumina HiSeq 2000 (Arabidopsis thaliana) | GSM2514742 | xrn3-8_small_RNA_rep3 |
| GSE129249 | GSE129248 | GPL198 | Affymetrix Arabidopsis ATH1 Genome Array [ATH1-121501] | GSM3703534 | WT | GSE103924 | GPL13222 | Illumina HiSeq 2000 (Arabidopsis thaliana) | GSM2786261 | WTCol0-0-1 |
| GSE129249 | GSE129248 | GPL198 | Affymetrix Arabidopsis ATH1 Genome Array [ATH1-121501] | GSM3703535 | WT_1 | GSE103924 | GPL13222 | Illumina HiSeq 2000 (Arabidopsis thaliana) | GSM2786261 | WTCol0-0-1 |
| GSE129249 | GSE129248 | GPL198 | Affymetrix Arabidopsis ATH1 Genome Array [ATH1-121501] | GSM3703534 | WT | GSE103924 | GPL13222 | Illumina HiSeq 2000 (Arabidopsis thaliana) | GSM2786262 | WTCol0-0-2 |
| GSE129249 | GSE129248 | GPL198 | Affymetrix Arabidopsis ATH1 Genome Array [ATH1-121501] | GSM3703535 | WT_1 | GSE103924 | GPL13222 | Illumina HiSeq 2000 (Arabidopsis thaliana) | GSM2786262 | WTCol0-0-2 |
| GSE129249 | GSE129248 | GPL198 | Affymetrix Arabidopsis ATH1 Genome Array [ATH1-121501] | GSM3703534 | WT | GSE103924 | GPL13222 | Illumina HiSeq 2000 (Arabidopsis thaliana) | GSM2786263 | WTCol0-0-3 |
| GSE129249 | GSE129248 | GPL198 | Affymetrix Arabidopsis ATH1 Genome | GSM3703535 | WT_1 | GSE103924 | GPL13222 | Illumina HiSeq 2000 (Arabidopsis thaliana) | GSM2786263 | WTCol0-0-3 |

| | | | | | | | | | |
|---|---|---|---|---|---|---|---|---|---|
| | | Array [ATH1-121501] | | | | | | | |
| GSE97670 | GPL198 | Affymetrix Arabidopsis ATH1 Genome Array [ATH1-121501] | GSM2575031 | Col-0 replicate 1 | GSE106370 | GPL17639 | Illumina HiSeq 2500 (Arabidopsis thaliana) | GSM2836681 | Col-0_replicate1 |
| GSE97670 | GPL198 | Affymetrix Arabidopsis ATH1 Genome Array [ATH1-121501] | GSM2575032 | Col-0 replicate 2 | GSE106370 | GPL17639 | Illumina HiSeq 2500 (Arabidopsis thaliana) | GSM2836681 | Col-0_replicate1 |
| GSE97670 | GPL198 | Affymetrix Arabidopsis ATH1 Genome Array [ATH1-121501] | GSM2575033 | Col-0 replicate 3 | GSE106370 | GPL17639 | Illumina HiSeq 2500 (Arabidopsis thaliana) | GSM2836681 | Col-0_replicate1 |
| GSE97670 | GPL198 | Affymetrix Arabidopsis ATH1 Genome Array [ATH1-121501] | GSM2575031 | Col-0 replicate 1 | GSE106370 | GPL17639 | Illumina HiSeq 2500 (Arabidopsis thaliana) | GSM2836682 | Col-0_replicate2 |
| GSE97670 | GPL198 | Affymetrix Arabidopsis ATH1 Genome Array [ATH1-121501] | GSM2575032 | Col-0 replicate 2 | GSE106370 | GPL17639 | Illumina HiSeq 2500 (Arabidopsis thaliana) | GSM2836682 | Col-0_replicate2 |
| GSE97670 | GPL198 | Affymetrix Arabidopsis ATH1 Genome Array [ATH1-121501] | GSM2575033 | Col-0 replicate 3 | GSE106370 | GPL17639 | Illumina HiSeq 2500 (Arabidopsis thaliana) | GSM2836682 | Col-0_replicate2 |
| GSE97670 | GPL198 | Affymetrix Arabidopsis ATH1 Genome Array [ATH1-121501] | GSM2575036 | shr replicate 3 | GSE106370 | GPL17639 | Illumina HiSeq 2500 (Arabidopsis thaliana) | GSM2836683 | Shr-2_replicate1 |
| GSE97670 | GPL198 | Affymetrix Arabidopsis ATH1 Genome Array [ATH1-121501] | GSM2575034 | shr replicate 1 | GSE106370 | GPL17639 | Illumina HiSeq 2500 (Arabidopsis thaliana) | GSM2836683 | Shr-2_replicate1 |
| GSE97670 | GPL198 | Affymetrix Arabidopsis ATH1 Genome Array | GSM2575035 | shr replicate 2 | GSE106370 | GPL17639 | Illumina HiSeq 2500 (Arabidopsis thaliana) | GSM2836683 | Shr-2_replicate1 |

| | | | | | | | | | | |
|---|---|---|---|---|---|---|---|---|---|---|
| | GSE97670 | GPL198 | [ATH1-121501] Affymetrix Arabidopsis ATH1 Genome Array | GSM2575035 | shr replicate 2 | GSE106370 | GPL17639 | Illumina HiSeq 2500 (Arabidopsis thaliana) | GSM2836684 | Shr-2_replicate2 |
| | GSE97670 | GPL198 | [ATH1-121501] Affymetrix Arabidopsis ATH1 Genome Array | GSM2575034 | shr replicate 1 | GSE106370 | GPL17639 | Illumina HiSeq 2500 (Arabidopsis thaliana) | GSM2836684 | Shr-2_replicate2 |
| | GSE97670 | GPL198 | [ATH1-121501] Affymetrix Arabidopsis ATH1 Genome Array | GSM2575036 | shr replicate 3 | GSE106370 | GPL17639 | Illumina HiSeq 2500 (Arabidopsis thaliana) | GSM2836684 | Shr-2_replicate2 |
| GSE115555 | GSE115486 | GPL198 | [ATH1-121501] Affymetrix Arabidopsis ATH1 Genome Array | GSM3179200 | ROOT ZONE I (0.5mm) REP 1 | GSE115554 | GPL19580 | Illumina NextSeq 500 (Arabidopsis thaliana) | GSM3181995 | ROOT ZONE I (0.5mm) REP 1 [RNA-seq] |
| GSE115555 | GSE115486 | GPL198 | [ATH1-121501] Affymetrix Arabidopsis ATH1 Genome Array | GSM3179201 | ROOT ZONE I (0.5mm) REP 2 | GSE115554 | GPL19580 | Illumina NextSeq 500 (Arabidopsis thaliana) | GSM3181995 | ROOT ZONE I (0.5mm) REP 1 [RNA-seq] |
| GSE115555 | GSE115486 | GPL198 | [ATH1-121501] Affymetrix Arabidopsis ATH1 Genome Array | GSM3179202 | ROOT ZONE I (0.5mm) REP 3 | GSE115554 | GPL19580 | Illumina NextSeq 500 (Arabidopsis thaliana) | GSM3181995 | ROOT ZONE I (0.5mm) REP 1 [RNA-seq] |
| GSE115555 | GSE115486 | GPL198 | [ATH1-121501] Affymetrix Arabidopsis ATH1 Genome Array | GSM3179200 | ROOT ZONE I (0.5mm) REP 1 | GSE115554 | GPL19580 | Illumina NextSeq 500 (Arabidopsis thaliana) | GSM3181996 | ROOT ZONE I (0.5mm) REP 2 [RNA-seq] |
| GSE115555 | GSE115486 | GPL198 | [ATH1-121501] Affymetrix Arabidopsis ATH1 Genome Array | GSM3179201 | ROOT ZONE I (0.5mm) REP 2 | GSE115554 | GPL19580 | Illumina NextSeq 500 (Arabidopsis thaliana) | GSM3181996 | ROOT ZONE I (0.5mm) REP 2 [RNA-seq] |
| GSE115555 | GSE115486 | GPL198 | [ATH1-121501] Affymetrix Arabidopsis ATH1 Genome Array | GSM3179202 | ROOT ZONE I (0.5mm) REP 3 | GSE115554 | GPL19580 | Illumina NextSeq 500 (Arabidopsis thaliana) | GSM3181996 | ROOT ZONE I (0.5mm) REP 2 [RNA-seq] |
| GSE11555 | GSE11548 | GPL198 | [ATH1-121501] | GSM3179200 | ROOT ZONE I | GSE11555 | GPL19580 | Illumina | GSM3181997 | ROOT ZONE I (0.5mm) |

| | | | | | | | | | | |
|---|---|---|---|---|---|---|---|---|---|---|
| 5 | 6 | | Affymetrix Arabidopsis ATH1 Genome Array [ATH1-121501] | | (0.5mm) REP 1 | 4 | | NextSeq 500 (Arabidopsis thaliana) | | REP 3 [RNA-seq] |
| GSE115555 | GSE115486 | GPL198 | Affymetrix Arabidopsis ATH1 Genome Array [ATH1-121501] | GSM3179201 | ROOT ZONE I (0.5mm) REP 2 | GSE115554 | GPL19580 | Illumina NextSeq 500 (Arabidopsis thaliana) | GSM3181997 | ROOT ZONE I (0.5mm) REP 3 [RNA-seq] |
| GSE115555 | GSE115486 | GPL198 | Affymetrix Arabidopsis ATH1 Genome Array [ATH1-121501] | GSM3179202 | ROOT ZONE I (0.5mm) REP 3 | GSE115554 | GPL19580 | Illumina NextSeq 500 (Arabidopsis thaliana) | GSM3181997 | ROOT ZONE I (0.5mm) REP 3 [RNA-seq] |
| GSE115555 | GSE115486 | GPL198 | Affymetrix Arabidopsis ATH1 Genome Array [ATH1-121501] | GSM3179204 | ROOT ZONE II (1.5mm) REP 2 | GSE115554 | GPL19580 | Illumina NextSeq 500 (Arabidopsis thaliana) | GSM3181998 | ROOT ZONE II (1.5mm) REP 1 [RNA-seq] |
| GSE115555 | GSE115486 | GPL198 | Affymetrix Arabidopsis ATH1 Genome Array [ATH1-121501] | GSM3179203 | ROOT ZONE II (1.5mm) REP 1 | GSE115554 | GPL19580 | Illumina NextSeq 500 (Arabidopsis thaliana) | GSM3181998 | ROOT ZONE II (1.5mm) REP 1 [RNA-seq] |
| GSE115555 | GSE115486 | GPL198 | Affymetrix Arabidopsis ATH1 Genome Array [ATH1-121501] | GSM3179205 | ROOT ZONE II (1.5mm) REP 3 | GSE115554 | GPL19580 | Illumina NextSeq 500 (Arabidopsis thaliana) | GSM3181998 | ROOT ZONE II (1.5mm) REP 1 [RNA-seq] |
| GSE115555 | GSE115486 | GPL198 | Affymetrix Arabidopsis ATH1 Genome Array [ATH1-121501] | GSM3179203 | ROOT ZONE II (1.5mm) REP 1 | GSE115554 | GPL19580 | Illumina NextSeq 500 (Arabidopsis thaliana) | GSM3181999 | ROOT ZONE II (1.5mm) REP 2 [RNA-seq] |
| GSE115555 | GSE115486 | GPL198 | Affymetrix Arabidopsis ATH1 Genome Array [ATH1-121501] | GSM3179205 | ROOT ZONE II (1.5mm) REP 3 | GSE115554 | GPL19580 | Illumina NextSeq 500 (Arabidopsis thaliana) | GSM3181999 | ROOT ZONE II (1.5mm) REP 2 [RNA-seq] |
| GSE115555 | GSE115486 | GPL198 | Affymetrix Arabidopsis ATH1 Genome Array [ATH1-121501] | GSM3179204 | ROOT ZONE II (1.5mm) REP 2 | GSE115554 | GPL19580 | Illumina NextSeq 500 (Arabidopsis thaliana) | GSM3181999 | ROOT ZONE II (1.5mm) REP 2 [RNA-seq] |
| GSE115555 | GSE115486 | GPL198 | Affymetrix | GSM3179203 | ROOT ZONE II (1.5mm) REP 1 | GSE115554 | GPL19580 | Illumina NextSeq 500 | GSM3182000 | ROOT ZONE II (1.5mm) REP 3 [RNA-seq] |

| | | | | | | | | | | |
|---|---|---|---|---|---|---|---|---|---|---|
| | | | Arabidopsis ATH1 Genome Array | | | | | (Arabidopsis thaliana) | | |
| GSE115555 | GSE115486 | GPL198 | [ATH1-121501] Affymetrix Arabidopsis ATH1 Genome Array | GSM3179204 | ROOT ZONE II (1.5mm) REP 2 | GSE115554 | GPL19580 | Illumina NextSeq 500 (Arabidopsis thaliana) | GSM3182000 | ROOT ZONE II (1.5mm) REP 3 [RNA-seq] |
| GSE115555 | GSE115486 | GPL198 | [ATH1-121501] Affymetrix Arabidopsis ATH1 Genome Array | GSM3179205 | ROOT ZONE II (1.5mm) REP 3 | GSE115554 | GPL19580 | Illumina NextSeq 500 (Arabidopsis thaliana) | GSM3182000 | ROOT ZONE II (1.5mm) REP 3 [RNA-seq] |

| C | array | | | | | seq | | | | | |
|---|---|---|---|---|---|---|---|---|---|---|---|
| superseries | series | platform | platform desc. | sample | sample desc. | series | platform | platform desc. | sample | sample desc. | sample code |
| NA | GSE43358 | GPL570 | [HG-U133_Plus_2] Affymetrix Human Genome U133 Plus 2.0 Array | GSM1060915 | HER2-02 | EGAD00001000627 | GPL11154 | Illumina HiSeq 2000 | HER2.02 | HER2-02 | HER2.02 |
| NA | GSE43358 | GPL570 | [HG-U133_Plus_2] Affymetrix Human Genome U133 Plus 2.0 Array | GSM1060921 | HER2-03 | EGAD00001000627 | GPL11154 | Illumina HiSeq 2000 | HER2.03 | HER2-03 | HER2.03 |
| NA | GSE43358 | GPL570 | [HG-U133_Plus_2] Affymetrix Human Genome U133 Plus 2.0 Array | GSM1060909 | HER2-13 | EGAD00001000627 | GPL11154 | Illumina HiSeq 2000 | HER2.13 | HER2-13 | HER2.13 |
| NA | GSE43358 | GPL570 | [HG-U133_Plus_2] Affymetrix Human Genome U133 Plus 2.0 Array | GSM1060910 | HER2-14 | EGAD00001000627 | GPL11154 | Illumina HiSeq 2000 | HER2.14 | HER2-14 | HER2.14 |
| NA | GSE43358 | GPL570 | [HG-U133_Plus_2] Affymetrix Human Genome U133 Plus 2.0 Array | GSM1060910 | HER2-14 | EGAD00001000627 | GPL11154 | Illumina HiSeq 2000 | HER2.14.N | HER2-14 | HER2.14.N |
| NA | GSE43358 | GPL570 | [HG-U133_Plus_2] Affymetrix Human Genome U133 | GSM1060911 | HER2-15 | EGAD0000 | GPL11154 | Illumina HiSeq 2000 | HER2.15 | HER2-15 | HER2.15 |

| | | | | | | | | | | | |
|---|---|---|---|---|---|---|---|---|---|---|---|
| | | | Plus 2.0 Array [HG-U133_Plus_2] | | | 1000627 | | | | | |
| NA | GSE43358 | GPL570 | Affymetrix Human Genome U133 Plus 2.0 Array [HG-U133_Plus_2] | GSM1060912 | HER2-16 | EGAD0000 1000627 | GPL11154 | Illumina HiSeq 2000 | HER2.16 | HER2-16 | HER2.16 |
| NA | GSE43358 | GPL570 | Affymetrix Human Genome U133 Plus 2.0 Array [HG-U133_Plus_2] | GSM1060913 | HER2-18 | EGAD0000 1000627 | GPL11154 | Illumina HiSeq 2000 | HER2.18 | HER2-18 | HER2.18 |
| NA | GSE43358 | GPL570 | Affymetrix Human Genome U133 Plus 2.0 Array [HG-U133_Plus_2] | GSM1060914 | HER2-19 | EGAD0000 1000627 | GPL11154 | Illumina HiSeq 2000 | HER2.19 | HER2-19 | HER2.19 |
| NA | GSE43358 | GPL570 | Affymetrix Human Genome U133 Plus 2.0 Array [HG-U133_Plus_2] | GSM1060916 | HER2-20 | EGAD0000 1000627 | GPL11154 | Illumina HiSeq 2000 | HER2.20 | HER2-20 | HER2.20 |
| NA | GSE43358 | GPL570 | Affymetrix Human Genome U133 Plus 2.0 Array [HG-U133_Plus_2] | GSM1060917 | HER2-21 | EGAD0000 1000627 | GPL11154 | Illumina HiSeq 2000 | HER2.21 | HER2-21 | HER2.21 |
| NA | GSE43358 | GPL570 | Affymetrix Human Genome U133 Plus 2.0 Array [HG-U133_Plus_2] | GSM1060918 | HER2-22 | EGAD0000 1000627 | GPL11154 | Illumina HiSeq 2000 | HER2.22 | HER2-22 | HER2.22 |
| NA | GSE43358 | GPL570 | Affymetrix Human Genome U133 Plus 2.0 Array [HG-U133_Plus_2] | GSM1060919 | HER2-23 | EGAD0000 1000627 | GPL11154 | Illumina HiSeq 2000 | HER2.23 | HER2-23 | HER2.23 |
| NA | GSE43358 | GPL570 | Affymetrix Human Genome U133 Plus 2.0 Array [HG-U133_Plus_2] | GSM1060920 | HER2-24 | EGAD0000 1000627 | GPL11154 | Illumina HiSeq 2000 | HER2.24 | HER2-24 | HER2.24 |
| NA | GSE43358 | GPL570 | Affymetrix Human Genome U133 Plus 2.0 Array | GSM1060922 | HER2-25 | EGAD0000 1000627 | GPL11154 | Illumina HiSeq 2000 | HER2.25 | HER2-25 | HER2.25 |

| | | | | | | | | | | | |
|---|---|---|---|---|---|---|---|---|---|---|---|
| NA | GSE43358 | GPL570 | [HG-U133_Plus_2] Affymetrix Human Genome U133 Plus 2.0 Array | GSM1060939 | LUMA-04 | EGAD00001000627 | GPL11154 | Illumina HiSeq 2000 | LUMA.04 | LUMA-04 | LUMA.04 |
| NA | GSE43358 | GPL570 | [HG-U133_Plus_2] Affymetrix Human Genome U133 Plus 2.0 Array | GSM1060923 | LUMA-18 | EGAD00001000627 | GPL11154 | Illumina HiSeq 2000 | LUMA.18 | LUMA-18 | LUMA.18 |
| NA | GSE43358 | GPL570 | [HG-U133_Plus_2] Affymetrix Human Genome U133 Plus 2.0 Array | GSM1060923 | LUMA-18 | EGAD00001000627 | GPL11154 | Illumina HiSeq 2000 | LUMA.18.N | LUMA-18 | LUMA.18.N |
| NA | GSE43358 | GPL570 | [HG-U133_Plus_2] Affymetrix Human Genome U133 Plus 2.0 Array | GSM1060924 | LUMA-19 | EGAD00001000627 | GPL11154 | Illumina HiSeq 2000 | LUMA.19 | LUMA-19 | LUMA.19 |
| NA | GSE43358 | GPL570 | [HG-U133_Plus_2] Affymetrix Human Genome U133 Plus 2.0 Array | GSM1060924 | LUMA-19 | EGAD00001000627 | GPL11154 | Illumina HiSeq 2000 | LUMA.19.N | LUMA-19 | LUMA.19.N |
| NA | GSE43358 | GPL570 | [HG-U133_Plus_2] Affymetrix Human Genome U133 Plus 2.0 Array | GSM1060925 | LUMA-20 | EGAD00001000627 | GPL11154 | Illumina HiSeq 2000 | LUMA.20 | LUMA-20 | LUMA.20 |
| NA | GSE43358 | GPL570 | [HG-U133_Plus_2] Affymetrix Human Genome U133 Plus 2.0 Array | GSM1060925 | LUMA-20 | EGAD00001000627 | GPL11154 | Illumina HiSeq 2000 | LUMA.20.N | LUMA-20 | LUMA.20.N |
| NA | GSE43358 | GPL570 | [HG-U133_Plus_2] Affymetrix Human Genome U133 Plus 2.0 Array | GSM1060926 | LUMA-21 | EGAD00001000627 | GPL11154 | Illumina HiSeq 2000 | LUMA.21 | LUMA-21 | LUMA.21 |
| NA | GSE43358 | GPL570 | [HG-U133_Plus_2] Affymetrix Human Genome U133 Plus 2.0 Array | GSM1060927 | LUMA-22 | EGAD00001000627 | GPL11154 | Illumina HiSeq 2000 | LUMA.22 | LUMA-22 | LUMA.22 |
| NA | GSE43358 | GPL570 | [HG- | GSM1060928 | LUMA-23 | EGAD0000 | GPL11154 | Illumina HiSeq | LUMA.23 | LUMA-23 | LUMA.23 |

| | | | | | | | | | | | |
|---|---|---|---|---|---|---|---|---|---|---|---|
| | | | U133_Plus_2] | | | | | | | | |
| | | | Affymetrix Human Genome U133 Plus 2.0 Array [HG-U133_Plus_2] | | | 1000627 | | 2000 | | | |
| NA | GSE43358 | GPL570 | Affymetrix Human Genome U133 Plus 2.0 Array [HG-U133_Plus_2] | GSM1060929 | LUMA-24 | EGAD00001000627 | GPL11154 | Illumina HiSeq 2000 | LUMA.24.N | LUMA-24 | LUMA.24.N |
| NA | GSE43358 | GPL570 | Affymetrix Human Genome U133 Plus 2.0 Array [HG-U133_Plus_2] | GSM1060930 | LUMA-25 | EGAD00001000627 | GPL11154 | Illumina HiSeq 2000 | LUMA.25 | LUMA-25 | LUMA.25 |
| NA | GSE43358 | GPL570 | Affymetrix Human Genome U133 Plus 2.0 Array [HG-U133_Plus_2] | GSM1060931 | LUMA-26 | EGAD00001000627 | GPL11154 | Illumina HiSeq 2000 | LUMA.26 | LUMA-26 | LUMA.26 |
| NA | GSE43358 | GPL570 | Affymetrix Human Genome U133 Plus 2.0 Array [HG-U133_Plus_2] | GSM1060932 | LUMA-27 | EGAD00001000627 | GPL11154 | Illumina HiSeq 2000 | LUMA.27 | LUMA-27 | LUMA.27 |
| NA | GSE43358 | GPL570 | Affymetrix Human Genome U133 Plus 2.0 Array [HG-U133_Plus_2] | GSM1060936 | LUMA-31 | EGAD00001000627 | GPL11154 | Illumina HiSeq 2000 | LUMA.31 | LUMA-31 | LUMA.31 |
| NA | GSE43358 | GPL570 | Affymetrix Human Genome U133 Plus 2.0 Array [HG-U133_Plus_2] | GSM1060937 | LUMA-32 | EGAD00001000627 | GPL11154 | Illumina HiSeq 2000 | LUMA.32 | LUMA-32 | LUMA.32 |
| NA | GSE43358 | GPL570 | Affymetrix Human Genome U133 Plus 2.0 Array [HG-U133_Plus_2] | GSM1060938 | LUMA-33 | EGAD00001000627 | GPL11154 | Illumina HiSeq 2000 | LUMA.33 | LUMA-33 | LUMA.33 |
| NA | GSE43358 | GPL570 | Affymetrix Human Genome U133 Plus 2.0 Array [HG- | GSM1060940 | LUMB-01 | EGAD00001000627 | GPL11154 | Illumina HiSeq 2000 | LUMB.01 | LUMB-01 | LUMB.01 |
| NA | GSE43358 | GPL570 | U133_Plus_2] | GSM1060948 | LUMB-03 | EGAD00001000627 | GPL11154 | Illumina HiSeq 2000 | LUMB.03 | LUMB-03 | LUMB.03 |

| | | | | | | | | | | | |
|---|---|---|---|---|---|---|---|---|---|---|---|
| | | | Affymetrix Human Genome U133 Plus 2.0 Array [HG-U133_Plus_2] | | | | | | | | |
| NA | GSE43358 | GPL570 | Affymetrix Human Genome U133 Plus 2.0 Array [HG-U133_Plus_2] | GSM1060949 | LUMB-05 | EGAD00001000627 | GPL11154 | Illumina HiSeq 2000 | LUMB.05 | LUMB-05 | LUMB.05 |
| NA | GSE43358 | GPL570 | Affymetrix Human Genome U133 Plus 2.0 Array [HG-U133_Plus_2] | GSM1060943 | LUMB-18 | EGAD00001000627 | GPL11154 | Illumina HiSeq 2000 | LUMB.18 | LUMB-18 | LUMB.18 |
| NA | GSE43358 | GPL570 | Affymetrix Human Genome U133 Plus 2.0 Array [HG-U133_Plus_2] | GSM1060944 | LUMB-19 | EGAD00001000627 | GPL11154 | Illumina HiSeq 2000 | LUMB.19 | LUMB-19 | LUMB.19 |
| NA | GSE43358 | GPL570 | Affymetrix Human Genome U133 Plus 2.0 Array [HG-U133_Plus_2] | GSM1060945 | LUMB-20 | EGAD00001000627 | GPL11154 | Illumina HiSeq 2000 | LUMB.20 | LUMB-20 | LUMB.20 |
| NA | GSE43358 | GPL570 | Affymetrix Human Genome U133 Plus 2.0 Array [HG-U133_Plus_2] | GSM1060946 | LUMB-21 | EGAD00001000627 | GPL11154 | Illumina HiSeq 2000 | LUMB.21 | LUMB-21 | LUMB.21 |
| NA | GSE43358 | GPL570 | Affymetrix Human Genome U133 Plus 2.0 Array [HG-U133_Plus_2] | GSM1060946 | LUMB-21 | EGAD00001000627 | GPL11154 | Illumina HiSeq 2000 | LUMB.22 | LUMB-21 | LUMB.22 |
| NA | GSE43358 | GPL570 | Affymetrix Human Genome U133 Plus 2.0 Array [HG-U133_Plus_2] | GSM1060947 | LUMB-23 | EGAD00001000627 | GPL11154 | Illumina HiSeq 2000 | LUMB.23 | LUMB-23 | LUMB.23 |
| NA | GSE43358 | GPL570 | Affymetrix Human Genome U133 Plus 2.0 Array [HG-U133_Plus_2] | GSM1060947 | LUMB-23 | EGAD00001000627 | GPL11154 | Illumina HiSeq 2000 | LUMB.23.N | LUMB-23 | LUMB.23.N |
| NA | GSE43358 | GPL570 | Affymetrix Human | GSM1060950 | TN-01 | EGAD00001000627 | GPL11154 | Illumina HiSeq 2000 | TN.01 | TN-01 | TN.01 |

| | | | | | | | | | | | | | |
|---|---|---|---|---|---|---|---|---|---|---|---|---|---|
| | | | Genome U133 Plus 2.0 Array [HG-U133_Plus_2] | | | | | | | | | | |
| NA | GSE43358 | GPL570 | Affymetrix Human Genome U133 Plus 2.0 Array [HG-U133_Plus_2] | GSM1060956 | TN-02 | | EGAD00001000627 | GPL11154 | Illumina HiSeq 2000 | TN.02 | TN-02 | | TN.02 |
| NA | GSE43358 | GPL570 | Affymetrix Human Genome U133 Plus 2.0 Array [HG-U133_Plus_2] | GSM1060965 | TN-03 | | EGAD00001000627 | GPL11154 | Illumina HiSeq 2000 | TN.03 | TN-03 | | TN.03 |
| NA | GSE43358 | GPL570 | Affymetrix Human Genome U133 Plus 2.0 Array [HG-U133_Plus_2] | GSM1060966 | TN-05 | | EGAD00001000627 | GPL11154 | Illumina HiSeq 2000 | TN.05 | TN-05 | | TN.05 |
| NA | GSE43358 | GPL570 | Affymetrix Human Genome U133 Plus 2.0 Array [HG-U133_Plus_2] | GSM1060951 | TN-15 | | EGAD00001000627 | GPL11154 | Illumina HiSeq 2000 | TN.15 | TN-15 | | TN.15 |
| NA | GSE43358 | GPL570 | Affymetrix Human Genome U133 Plus 2.0 Array [HG-U133_Plus_2] | GSM1060951 | TN-15 | | EGAD00001000627 | GPL11154 | Illumina HiSeq 2000 | TN.15.N | TN-15 | | TN.15.N |
| NA | GSE43358 | GPL570 | Affymetrix Human Genome U133 Plus 2.0 Array [HG-U133_Plus_2] | GSM1060952 | TN-16 | | EGAD00001000627 | GPL11154 | Illumina HiSeq 2000 | TN.16 | TN-16 | | TN.16 |
| NA | GSE43358 | GPL570 | Affymetrix Human Genome U133 Plus 2.0 Array [HG-U133_Plus_2] | GSM1060953 | TN-17 | | EGAD00001000627 | GPL11154 | Illumina HiSeq 2000 | TN.17 | TN-17 | | TN.17 |
| NA | GSE43358 | GPL570 | Affymetrix Human Genome U133 Plus 2.0 Array [HG-U133_Plus_2] | GSM1060953 | TN-17 | | EGAD00001000627 | GPL11154 | Illumina HiSeq 2000 | TN.17.N | TN-17 | | TN.17.N |
| NA | GSE43358 | GPL570 | Affymetrix Human Genome U133 | GSM1060954 | TN-18 | | EGAD00001000627 | GPL11154 | Illumina HiSeq 2000 | TN.18 | TN-18 | | TN.18 |

| | | | | | | | | | | | | | |
|---|---|---|---|---|---|---|---|---|---|---|---|---|---|
| | | | Plus 2.0 Array [HG-U133_Plus_2] | | | | | | | | | | |
| NA | GSE43358 | GPL570 | Affymetrix Human Genome U133 Plus 2.0 Array [HG-U133_Plus_2] | GSM1060954 | TN-18 | | EGAD00001000627 | GPL11154 | Illumina HiSeq 2000 | TN.18.N | TN-18 | | TN.18.N |
| NA | GSE43358 | GPL570 | Affymetrix Human Genome U133 Plus 2.0 Array [HG-U133_Plus_2] | GSM1060955 | TN-19 | | EGAD00001000627 | GPL11154 | Illumina HiSeq 2000 | TN.19 | TN-19 | | TN.19 |
| NA | GSE43358 | GPL570 | Affymetrix Human Genome U133 Plus 2.0 Array [HG-U133_Plus_2] | GSM1060957 | TN-20 | | EGAD00001000627 | GPL11154 | Illumina HiSeq 2000 | TN.20 | TN-20 | | TN.20 |
| NA | GSE43358 | GPL570 | Affymetrix Human Genome U133 Plus 2.0 Array [HG-U133_Plus_2] | GSM1060958 | TN-21 | | EGAD00001000627 | GPL11154 | Illumina HiSeq 2000 | TN.21 | TN-21 | | TN.21 |
| NA | GSE43358 | GPL570 | Affymetrix Human Genome U133 Plus 2.0 Array [HG-U133_Plus_2] | GSM1060959 | TN-22 | | EGAD00001000627 | GPL11154 | Illumina HiSeq 2000 | TN.22 | TN-22 | | TN.22 |
| NA | GSE43358 | GPL570 | Affymetrix Human Genome U133 Plus 2.0 Array [HG-U133_Plus_2] | GSM1060960 | TN-23 | | EGAD00001000627 | GPL11154 | Illumina HiSeq 2000 | TN.23 | TN-23 | | TN.23 |
| NA | GSE43358 | GPL570 | Affymetrix Human Genome U133 Plus 2.0 Array [HG-U133_Plus_2] | GSM1060961 | TN-24 | | EGAD00001000627 | GPL11154 | Illumina HiSeq 2000 | TN.24 | TN-24 | | TN.24 |
| NA | GSE43358 | GPL570 | Affymetrix Human Genome U133 Plus 2.0 Array [HG-U133_Plus_2] | GSM1060962 | TN-25 | | EGAD00001000627 | GPL11154 | Illumina HiSeq 2000 | TN.25 | TN-25 | | TN.25 |
| NA | GSE43358 | GPL570 | Affymetrix Human Genome U133 Plus 2.0 Array | GSM1060963 | TN-26 | | EGAD00001000627 | GPL11154 | Illumina HiSeq 2000 | TN.26 | TN-26 | | TN.26 |

| | | | | | | | | | | | |
|---|---|---|---|---|---|---|---|---|---|---|---|
| NA | GSE43358 | GPL570 | [HG-U133_Plus_2] Affymetrix Human Genome U133 Plus 2.0 Array | GSM1060964 | TN-28 | EGAD00001000627 | GPL11154 | Illumina HiSeq 2000 | TN.28 | TN-28 | TN.28 |
| D superseries | array series | platform | platform desc. | sample | sample desc. | seq series | platform | platform desc. | sample | sample desc. | sample code |
| GSE46401 | GSE46394 | GPL13534 | Illumina HumanMethylation450 BeadChip (HumanMethylation450_15017482) | GSM1129686 | Atherosclerotic lesion 93 | GSE46327 | GPL11154 | Illumina HiSeq 2000 (Homo sapiens) | GSM1128658 | 93A | |
| GSE46401 | GSE46394 | GPL13534 | Illumina HumanMethylation450 BeadChip (HumanMethylation450_15017482) | GSM1129701 | Aortic tissue 93 | GSE46327 | GPL11154 | Illumina HiSeq 2000 (Homo sapiens) | GSM1128659 | 93N | |
| GSE52980 | GSE51954 | GPL13534 | Illumina HumanMethylation450 BeadChip (HumanMethylation450_15017482) | GSM1255809 | genomic DNA from young sun protected epidermis 1 | GSE52972 | GPL11154 | Illumina HiSeq 2000 (Homo sapiens) | GSM1279668 | bisfulfite treated genomic DNA from young sun protected epidermis 1 | P41 |
| GSE52980 | GSE51954 | GPL13534 | Illumina HumanMethylation450 BeadChip (HumanMethylation450_15017482) | GSM1255810 | genomic DNA from young sun protected epidermis 2 | GSE52972 | GPL11154 | Illumina HiSeq 2000 (Homo sapiens) | GSM1279671 | bisulfite treated genomic DNA from young sun protected epidermis 2 | E43 |
| GSE52980 | GSE51954 | GPL13534 | Illumina HumanMethylation450 BeadChip (HumanMethylation450_15017482) | GSM1255811 | genomic DNA from young sun protected epidermis 3 | GSE52972 | GPL11154 | Illumina HiSeq 2000 (Homo sapiens) | GSM1279672 | bisulfite treated genomic DNA from young sun protected epidermis 3 | P45 |
| GSE52980 | GSE51954 | GPL13534 | Illumina HumanMethylation450 BeadChip (HumanMethylation450_15017482) | GSM1255818 | genomic DNA from young sun exposed epidermis 1 | GSE52972 | GPL11154 | Illumina HiSeq 2000 (Homo sapiens) | GSM1279669 | bisfulfite treated genomic DNA from young sun exposed epidermis 1 | E41 |
| GSE52980 | GSE51954 | GPL13534 | Illumina HumanMethylatio | GSM1255819 | genomic DNA from young sun exposed | GSE52972 | GPL11154 | Illumina HiSeq 2000 (Homo | GSM1279670 | bisfulfite treated genomic DNA from | P43 |

| | | | | | | | | | | | |
|---|---|---|---|---|---|---|---|---|---|---|---|
| | | | n450 BeadChip (HumanMethylation450_15017482) | | epidermis 2 | | | sapiens) | | young sun exposed epidermis 2 | |
| GSE52980 | GSE51954 | GPL13534 | Illumina HumanMethylation450 BeadChip (HumanMethylation450_15017482) | GSM1255820 | genomic DNA from young sun exposed epidermis 3 | GSE52972 | GPL11154 | Illumina HiSeq 2000 (Homo sapiens) | GSM1279673 | bisulfite treated genomic DNA from young sun exposed epidermis 3 | E45 |
| GSE52980 | GSE51954 | GPL13534 | Illumina HumanMethylation450 BeadChip (HumanMethylation450_15017482) | GSM1255827 | genomic DNA from old sun protected epidermis 1 | GSE52972 | GPL11154 | Illumina HiSeq 2000 (Homo sapiens) | GSM1279674 | bisfulfite treated genomic DNA from old sun protected epidermis 1 | P47 |
| GSE52980 | GSE51954 | GPL13534 | Illumina HumanMethylation450 BeadChip (HumanMethylation450_15017482) | GSM1255828 | genomic DNA from old sun protected epidermis 2 | GSE52972 | GPL11154 | Illumina HiSeq 2000 (Homo sapiens) | GSM1279676 | bisfulfite treated genomic DNA from old sun protected epidermis 2 | P49 |
| GSE52980 | GSE51954 | GPL13534 | Illumina HumanMethylation450 BeadChip (HumanMethylation450_15017482) | GSM1255829 | genomic DNA from old sun protected epidermis 3 | GSE52972 | GPL11154 | Illumina HiSeq 2000 (Homo sapiens) | GSM1279678 | bisulfite treated genomic DNA from old sun protected epidermis 3 | P51 |
| GSE52980 | GSE51954 | GPL13534 | Illumina HumanMethylation450 BeadChip (HumanMethylation450_15017482) | GSM1255837 | genomic DNA from old sun exposed epidermis 1 | GSE52972 | GPL11154 | Illumina HiSeq 2000 (Homo sapiens) | GSM1279675 | bisfulfite treated genomic DNA from old sun exposed epidermis 1 | E47 |
| GSE52980 | GSE51954 | GPL13534 | Illumina HumanMethylation450 BeadChip (HumanMethylation450_15017482) | GSM1255838 | genomic DNA from old sun exposed epidermis 2 | GSE52972 | GPL11154 | Illumina HiSeq 2000 (Homo sapiens) | GSM1279677 | bisulfite treated genomic DNA from old sun exposed epidermis 2 | E49 |
| GSE52980 | GSE51954 | GPL13534 | Illumina HumanMethylation450 BeadChip (HumanMethylation450_15017482) | GSM1255839 | genomic DNA from old sun exposed epidermis 3 | GSE52972 | GPL11154 | Illumina HiSeq 2000 (Homo sapiens) | GSM1279679 | bisulfite treated genomic DNA from old sun exposed epidermis 3 | E51 |

| | | | | | | | | | | |
|---|---|---|---|---|---|---|---|---|---|---|
| GSE57361 | GSE57360 | GPL13534 | Illumina HumanMethylation450 BeadChip (HumanMethylation450_15017482) | GSM1381002 | Alzheimer's disease A09_151 | GSE57359 | GPL11154 | Illumina HiSeq 2000 (Homo sapiens) | GSM1380998 | Alzheimer's disease |
| GSE57361 | GSE57360 | GPL13534 | Illumina HumanMethylation450 BeadChip (HumanMethylation450_15017482) | GSM1381003 | Alzheimer's disease A08_127 | GSE57359 | GPL11154 | Illumina HiSeq 2000 (Homo sapiens) | GSM1380998 | Alzheimer's disease |
| GSE57361 | GSE57360 | GPL13534 | Illumina HumanMethylation450 BeadChip (HumanMethylation450_15017482) | GSM1381004 | Alzheimer's disease A09_26 | GSE57359 | GPL11154 | Illumina HiSeq 2000 (Homo sapiens) | GSM1380998 | Alzheimer's disease |
| GSE57361 | GSE57360 | GPL13534 | Illumina HumanMethylation450 BeadChip (HumanMethylation450_15017482) | GSM1381005 | Alzheimer's disease A10_51 | GSE57359 | GPL11154 | Illumina HiSeq 2000 (Homo sapiens) | GSM1380998 | Alzheimer's disease |
| GSE57361 | GSE57360 | GPL13534 | Illumina HumanMethylation450 BeadChip (HumanMethylation450_15017482) | GSM1381006 | Alzheimer's disease 31_05 | GSE57359 | GPL11154 | Illumina HiSeq 2000 (Homo sapiens) | GSM1380998 | Alzheimer's disease |
| GSE57361 | GSE57360 | GPL13534 | Illumina HumanMethylation450 BeadChip (HumanMethylation450_15017482) | GSM1381007 | Alzheimer's disease 17_08 | GSE57359 | GPL11154 | Illumina HiSeq 2000 (Homo sapiens) | GSM1380998 | Alzheimer's disease |
| GSE57361 | GSE57360 | GPL13534 | Illumina HumanMethylation450 BeadChip (HumanMethylation450_15017482) | GSM1381008 | Down Syndrome 35_06 | GSE57359 | GPL11154 | Illumina HiSeq 2000 (Homo sapiens) | GSM1381001 | Down’s syndrome |
| GSE57361 | GSE57360 | GPL13534 | Illumina HumanMethylation450 BeadChip (HumanMethylati | GSM1381009 | Down Syndrome 32_08 | GSE57359 | GPL11154 | Illumina HiSeq 2000 (Homo sapiens) | GSM1381001 | Down’s syndrome |

| | | | | | | | | | | |
|---|---|---|---|---|---|---|---|---|---|---|
| | | | on450_15017482 ) | | | | | | | |
| GSE57361 | GSE57360 | GPL13534 | Illumina HumanMethylation450 BeadChip (HumanMethylation450_15017482 ) | GSM1381010 | Down Syndrome with Alzheimer's disease 31_08 | GSE57359 | GPL11154 | Illumina HiSeq 2000 (Homo sapiens) | GSM1381001 | Down’s syndrome |
| GSE57361 | GSE57360 | GPL13534 | Illumina HumanMethylation450 BeadChip (HumanMethylation450_15017482 ) | GSM1381011 | Down Syndrome with Alzheimer's disease 31_07 | GSE57359 | GPL11154 | Illumina HiSeq 2000 (Homo sapiens) | GSM1381001 | Down’s syndrome |
| GSE57361 | GSE57360 | GPL13534 | Illumina HumanMethylation450 BeadChip (HumanMethylation450_15017482 ) | GSM1381012 | Down Syndrome with Alzheimer's disease 38_08 | GSE57359 | GPL11154 | Illumina HiSeq 2000 (Homo sapiens) | GSM1381001 | Down’s syndrome |
| GSE57361 | GSE57360 | GPL13534 | Illumina HumanMethylation450 BeadChip (HumanMethylation450_15017482 ) | GSM1381013 | Lewy Body disease DLB2 | GSE57359 | GPL11154 | Illumina HiSeq 2000 (Homo sapiens) | GSM1380999 | Dementia of Lewy bodies |
| GSE57361 | GSE57360 | GPL13534 | Illumina HumanMethylation450 BeadChip (HumanMethylation450_15017482 ) | GSM1381014 | Lewy Body disease 11_052 | GSE57359 | GPL11154 | Illumina HiSeq 2000 (Homo sapiens) | GSM1380999 | Dementia of Lewy bodies |
| GSE57361 | GSE57360 | GPL13534 | Illumina HumanMethylation450 BeadChip (HumanMethylation450_15017482 ) | GSM1381015 | Lewy Body disease BK-458 | GSE57359 | GPL11154 | Illumina HiSeq 2000 (Homo sapiens) | GSM1380999 | Dementia of Lewy bodies |
| GSE57361 | GSE57360 | GPL13534 | Illumina HumanMethylation450 BeadChip (HumanMethylation450_15017482 ) | GSM1381016 | Lewy Body disease BK-1066 | GSE57359 | GPL11154 | Illumina HiSeq 2000 (Homo sapiens) | GSM1380999 | Dementia of Lewy bodies |
| GSE57361 | GSE57360 | GPL13534 | Illumina HumanMethylatio | GSM1381017 | Lewy Body disease BK-424 | GSE57359 | GPL11154 | Illumina HiSeq 2000 (Homo | GSM1380999 | Dementia of Lewy bodies |

| | | | | | | | | | | |
|---|---|---|---|---|---|---|---|---|---|---|
| | | | n450 BeadChip (HumanMethylation450_15017482) | | | | | sapiens) | | |
| GSE57361 | GSE57360 | GPL13534 | Illumina HumanMethylation450 BeadChip (HumanMethylation450_15017482) | GSM1381018 | Lewy Body disease BK-1159 | GSE57359 | GPL11154 | Illumina HiSeq 2000 (Homo sapiens) | GSM1380999 | Dementia of Lewy bodies |
| GSE57361 | GSE57360 | GPL13534 | Illumina HumanMethylation450 BeadChip (HumanMethylation450_15017482) | GSM1381019 | Parkinson's disease BK_1027 | GSE57359 | GPL11154 | Illumina HiSeq 2000 (Homo sapiens) | GSM1381000 | Parkinson's disease |
| GSE57361 | GSE57360 | GPL13534 | Illumina HumanMethylation450 BeadChip (HumanMethylation450_15017482) | GSM1381020 | Parkinson's disease BK-680 | GSE57359 | GPL11154 | Illumina HiSeq 2000 (Homo sapiens) | GSM1381000 | Parkinson's disease |
| GSE57361 | GSE57360 | GPL13534 | Illumina HumanMethylation450 BeadChip (HumanMethylation450_15017482) | GSM1381021 | Parkinson's disease BK-574 | GSE57359 | GPL11154 | Illumina HiSeq 2000 (Homo sapiens) | GSM1381000 | Parkinson's disease |
| GSE57361 | GSE57360 | GPL13534 | Illumina HumanMethylation450 BeadChip (HumanMethylation450_15017482) | GSM1381022 | Parkinson's disease BK-1182 | GSE57359 | GPL11154 | Illumina HiSeq 2000 (Homo sapiens) | GSM1381000 | Parkinson's disease |
| GSE57361 | GSE57360 | GPL13534 | Illumina HumanMethylation450 BeadChip (HumanMethylation450_15017482) | GSM1381023 | Parkinson's disease BK-476 | GSE57359 | GPL11154 | Illumina HiSeq 2000 (Homo sapiens) | GSM1381000 | Parkinson's disease |
| GSE89474 | GSE89472 | GPL13534 | Illumina HumanMethylation450 BeadChip (HumanMethylation450_15017482) | GSM2373076 | TwinA_ALS | GSE89473 | GPL11154 | Illumina HiSeq 2000 (Homo sapiens) | GSM2373086 | ALS_Male_blood_A |

| | | | | | | | | | | |
|---|---|---|---|---|---|---|---|---|---|---|
| GSE89474 | GSE89472 | GPL13534 | Illumina HumanMethylation450 BeadChip (HumanMethylation450_15017482) | GSM2373077 | TwinB_Control | GSE89473 | GPL11154 | Illumina HiSeq 2000 (Homo sapiens) | GSM2373087 | control_Male_blood_B |
| GSE89474 | GSE89472 | GPL13534 | Illumina HumanMethylation450 BeadChip (HumanMethylation450_15017482) | GSM2373078 | TwinC_ALS | GSE89473 | GPL11154 | Illumina HiSeq 2000 (Homo sapiens) | GSM2373088 | ALS_Male_blood_C |
| GSE89474 | GSE89472 | GPL13534 | Illumina HumanMethylation450 BeadChip (HumanMethylation450_15017482) | GSM2373079 | TwinD_Control | GSE89473 | GPL11154 | Illumina HiSeq 2000 (Homo sapiens) | GSM2373089 | control_Male_blood_D |
| GSE89474 | GSE89472 | GPL13534 | Illumina HumanMethylation450 BeadChip (HumanMethylation450_15017482) | GSM2373080 | TwinE_ALS | GSE89473 | GPL11154 | Illumina HiSeq 2000 (Homo sapiens) | GSM2373090 | ALS_Female_blood_E |
| GSE89474 | GSE89472 | GPL13534 | Illumina HumanMethylation450 BeadChip (HumanMethylation450_15017482) | GSM2373081 | TwinF_Control | GSE89473 | GPL11154 | Illumina HiSeq 2000 (Homo sapiens) | GSM2373091 | control_Female_blood_F |
| GSE89474 | GSE89472 | GPL13534 | Illumina HumanMethylation450 BeadChip (HumanMethylation450_15017482) | GSM2373082 | TwinG_ALS | GSE89473 | GPL11154 | Illumina HiSeq 2000 (Homo sapiens) | GSM2373092 | ALS_Female_blood_G |
| GSE89474 | GSE89472 | GPL13534 | Illumina HumanMethylation450 BeadChip (HumanMethylation450_15017482) | GSM2373083 | TwinH_Control | GSE89473 | GPL11154 | Illumina HiSeq 2000 (Homo sapiens) | GSM2373093 | control_Female_blood_H |
| GSE89474 | GSE89472 | GPL13534 | Illumina HumanMethylation450 BeadChip (HumanMethylati | GSM2373084 | TwinI_ALS | GSE89473 | GPL11154 | Illumina HiSeq 2000 (Homo sapiens) | GSM2373094 | ALS_Female_blood_I |

| | | | | | | | | | | |
|---|---|---|---|---|---|---|---|---|---|---|
| | | | on450_15017482 ) | | | | | | | |
| GSE89474 | GSE89472 | GPL1353 4 | Illumina HumanMethylatio n450 BeadChip (HumanMethylati on450_15017482 ) | GSM2373085 | TwinJ_Control | GSE89473 | GPL11154 | Illumina HiSeq 2000 (Homo sapiens) | GSM2373095 | control_Female_blood_J |
| GSE92469 | GSE92462 | GPL1353 4 | Illumina HumanMethylatio n450 BeadChip (HumanMethylati on450_15017482 ) | GSM2430095 | 450K_GSC80 | GSE92460 | GPL16791 | Illumina HiSeq 2500 (Homo sapiens) | GSM2430057 | RRBS_GSC80 Sample 13 |
| GSE92469 | GSE92462 | GPL1353 4 | Illumina HumanMethylatio n450 BeadChip (HumanMethylati on450_15017482 ) | GSM2430095 | 450K_GSC80 | GSE92460 | GPL16791 | Illumina HiSeq 2500 (Homo sapiens) | GSM2430067 | TAB-RRBS_GSC80 Sample 23 |
| GSE92469 | GSE92462 | GPL1353 4 | Illumina HumanMethylatio n450 BeadChip (HumanMethylati on450_15017482 ) | GSM2430095 | 450K_GSC80 | GSE92460 | GPL16791 | Illumina HiSeq 2500 (Homo sapiens) | GSM2430079 | MAB-RRBS_GSC80 |
| GSE92469 | GSE92462 | GPL1353 4 | Illumina HumanMethylatio n450 BeadChip (HumanMethylati on450_15017482 ) | GSM2430097 | 450K_GSC64 | GSE92460 | GPL16791 | Illumina HiSeq 2500 (Homo sapiens) | GSM2430056 | RRBS_GSC64 Sample 12 |
| GSE92469 | GSE92462 | GPL1353 4 | Illumina HumanMethylatio n450 BeadChip (HumanMethylati on450_15017482 ) | GSM2430097 | 450K_GSC64 | GSE92460 | GPL16791 | Illumina HiSeq 2500 (Homo sapiens) | GSM2430066 | TAB-RRBS_GSC64 Sample 22 |
| GSE92469 | GSE92462 | GPL1353 4 | Illumina HumanMethylatio n450 BeadChip (HumanMethylati on450_15017482 ) | GSM2430097 | 450K_GSC64 | GSE92460 | GPL16791 | Illumina HiSeq 2500 (Homo sapiens) | GSM2430078 | MAB-RRBS_GSC64 |
| GSE92469 | GSE92462 | GPL1353 4 | Illumina HumanMethylatio | GSM2430097 | 450K_GSC64 | GSE92460 | GPL16791 | Illumina HiSeq 2500 (Homo | GSM2430082 | RRBS_GSC64FBS |

| | | | | | | | | | | |
|---|---|---|---|---|---|---|---|---|---|---|
| | | | n450 BeadChip (HumanMethylation450_15017482) | | | | | sapiens) | | |
| GSE92469 | GSE92462 | GPL13534 | Illumina HumanMethylation450 BeadChip (HumanMethylation450_15017482) | GSM2430097 | 450K_GSC64 | GSE92460 | GPL16791 | Illumina HiSeq 2500 (Homo sapiens) | GSM2430087 | TAB-RRBS_GSC64FBS |
| GSE92469 | GSE92462 | GPL13534 | Illumina HumanMethylation450 BeadChip (HumanMethylation450_15017482) | GSM2430099 | 450K_GSC102 | GSE92460 | GPL16791 | Illumina HiSeq 2500 (Homo sapiens) | GSM2430059 | RRBS_GSC102 Sample 15 |
| GSE92469 | GSE92462 | GPL13534 | Illumina HumanMethylation450 BeadChip (HumanMethylation450_15017482) | GSM2430099 | 450K_GSC102 | GSE92460 | GPL16791 | Illumina HiSeq 2500 (Homo sapiens) | GSM2430069 | TAB-RRBS_GSC102 Sample 25 |
| GSE92469 | GSE92462 | GPL13534 | Illumina HumanMethylation450 BeadChip (HumanMethylation450_15017482) | GSM2430099 | 450K_GSC102 | GSE92460 | GPL16791 | Illumina HiSeq 2500 (Homo sapiens) | GSM2430076 | MAB-RRBS_GSC102 Sample 35 |
| GSE92469 | GSE92462 | GPL13534 | Illumina HumanMethylation450 BeadChip (HumanMethylation450_15017482) | GSM2430100 | 450K_GSC6 | GSE92460 | GPL16791 | Illumina HiSeq 2500 (Homo sapiens) | GSM2430051 | RRBS_GSC6 Sample 7 |
| GSE92469 | GSE92462 | GPL13534 | Illumina HumanMethylation450 BeadChip (HumanMethylation450_15017482) | GSM2430100 | 450K_GSC6 | GSE92460 | GPL16791 | Illumina HiSeq 2500 (Homo sapiens) | GSM2430061 | TAB-RRBS_GSC6 Sample 17 |
| GSE92469 | GSE92462 | GPL13534 | Illumina HumanMethylation450 BeadChip (HumanMethylation450_15017482) | GSM2430100 | 450K_GSC6 | GSE92460 | GPL16791 | Illumina HiSeq 2500 (Homo sapiens) | GSM2430077 | MAB-RRBS_GSC6 |

| | | | | | | | | | | |
|---|---|---|---|---|---|---|---|---|---|---|
| GSE92469 | GSE92462 | GPL13534 | Illumina HumanMethylation450 BeadChip (HumanMethylation450_15017482) | GSM2430100 | 450K_GSC6 | | GSE92460 | GPL16791 | Illumina HiSeq 2500 (Homo sapiens) | GSM2430081 | RRBS_GSC6FBS |
| GSE92469 | GSE92462 | GPL13534 | Illumina HumanMethylation450 BeadChip (HumanMethylation450_15017482) | GSM2430100 | 450K_GSC6 | | GSE92460 | GPL16791 | Illumina HiSeq 2500 (Homo sapiens) | GSM2430086 | TAB-RRBS_GSC6FBS |
| GSE92469 | GSE92462 | GPL13534 | Illumina HumanMethylation450 BeadChip (HumanMethylation450_15017482) | GSM2430101 | 450K_GSC14 | | GSE92460 | GPL16791 | Illumina HiSeq 2500 (Homo sapiens) | GSM2430054 | RRBS_GSC14 Sample 10 |
| GSE92469 | GSE92462 | GPL13534 | Illumina HumanMethylation450 BeadChip (HumanMethylation450_15017482) | GSM2430101 | 450K_GSC14 | | GSE92460 | GPL16791 | Illumina HiSeq 2500 (Homo sapiens) | GSM2430064 | TAB-RRBS_GSC14 Sample 20 |
| GSE92469 | GSE92462 | GPL13534 | Illumina HumanMethylation450 BeadChip (HumanMethylation450_15017482) | GSM2430101 | 450K_GSC14 | | GSE92460 | GPL16791 | Illumina HiSeq 2500 (Homo sapiens) | GSM2430073 | MAB-RRBS_GSC14 Sample 30 |
| GSE92469 | GSE92462 | GPL13534 | Illumina HumanMethylation450 BeadChip (HumanMethylation450_15017482) | GSM2430102 | 450K_GSC10 | | GSE92460 | GPL16791 | Illumina HiSeq 2500 (Homo sapiens) | GSM2430052 | RRBS_GSC10 Sample 8 |
| GSE92469 | GSE92462 | GPL13534 | Illumina HumanMethylation450 BeadChip (HumanMethylation450_15017482) | GSM2430102 | 450K_GSC10 | | GSE92460 | GPL16791 | Illumina HiSeq 2500 (Homo sapiens) | GSM2430062 | TAB-RRBS_GSC10 Sample 18 |
| GSE92469 | GSE92462 | GPL13534 | Illumina HumanMethylation450 BeadChip (HumanMethylati | GSM2430102 | 450K_GSC10 | | GSE92460 | GPL16791 | Illumina HiSeq 2500 (Homo sapiens) | GSM2430071 | MAB-RRBS_GSC10 Sample 28 |

| | | | | | | | | | | |
|---|---|---|---|---|---|---|---|---|---|---|
| | | | on450_15017482 ) | | | | | | | |
| GSE92469 | GSE92462 | GPL13534 | Illumina HumanMethylation450 BeadChip (HumanMethylation450_15017482 ) | GSM2430103 | 450K_GSC39 | GSE92460 | GPL16791 | Illumina HiSeq 2500 (Homo sapiens) | GSM2430055 | RRBS_GSC39 Sample 11 |
| GSE92469 | GSE92462 | GPL13534 | Illumina HumanMethylation450 BeadChip (HumanMethylation450_15017482 ) | GSM2430103 | 450K_GSC39 | GSE92460 | GPL16791 | Illumina HiSeq 2500 (Homo sapiens) | GSM2430065 | TAB-RRBS_GSC39 Sample 21 |
| GSE92469 | GSE92462 | GPL13534 | Illumina HumanMethylation450 BeadChip (HumanMethylation450_15017482 ) | GSM2430103 | 450K_GSC39 | GSE92460 | GPL16791 | Illumina HiSeq 2500 (Homo sapiens) | GSM2430074 | MAB-RRBS_GSC39 Sample 31 |
| GSE92469 | GSE92462 | GPL13534 | Illumina HumanMethylation450 BeadChip (HumanMethylation450_15017482 ) | GSM2430111 | 450K_GSC12 | GSE92460 | GPL16791 | Illumina HiSeq 2500 (Homo sapiens) | GSM2430053 | RRBS_GSC12 Sample 9 |
| GSE92469 | GSE92462 | GPL13534 | Illumina HumanMethylation450 BeadChip (HumanMethylation450_15017482 ) | GSM2430111 | 450K_GSC12 | GSE92460 | GPL16791 | Illumina HiSeq 2500 (Homo sapiens) | GSM2430063 | TAB-RRBS_GSC12 Sample 19 |
| GSE92469 | GSE92462 | GPL13534 | Illumina HumanMethylation450 BeadChip (HumanMethylation450_15017482 ) | GSM2430111 | 450K_GSC12 | GSE92460 | GPL16791 | Illumina HiSeq 2500 (Homo sapiens) | GSM2430072 | MAB-RRBS_GSC12 Sample 29 |
| GSE92469 | GSE92462 | GPL13534 | Illumina HumanMethylation450 BeadChip (HumanMethylation450_15017482 ) | GSM2430113 | 450K_GSC84 | GSE92460 | GPL16791 | Illumina HiSeq 2500 (Homo sapiens) | GSM2430058 | RRBS_GSC84 Sample 14 |
| GSE92469 | GSE92462 | GPL13534 | Illumina HumanMethylatio | GSM2430113 | 450K_GSC84 | GSE92460 | GPL16791 | Illumina HiSeq 2500 (Homo | GSM2430068 | TAB-RRBS_GSC84 Sample 24 |

| | | | | | | | | | | |
|---|---|---|---|---|---|---|---|---|---|---|
| | | | n450 BeadChip (HumanMethylation450_15017482) | | | | | sapiens) | | |
| GSE92469 | GSE92462 | GPL13534 | Illumina HumanMethylation450 BeadChip (HumanMethylation450_15017482) | GSM2430113 | 450K_GSC84 | GSE92460 | GPL16791 | Illumina HiSeq 2500 (Homo sapiens) | GSM2430075 | MAB-RRBS_GSC84 Sample 34 |
| GSE92469 | GSE92462 | GPL13534 | Illumina HumanMethylation450 BeadChip (HumanMethylation450_15017482) | GSM2430113 | 450K_GSC84 | GSE92460 | GPL16791 | Illumina HiSeq 2500 (Homo sapiens) | GSM2430083 | RRBS_GSC84FBS |
| GSE92469 | GSE92462 | GPL13534 | Illumina HumanMethylation450 BeadChip (HumanMethylation450_15017482) | GSM2430113 | 450K_GSC84 | GSE92460 | GPL16791 | Illumina HiSeq 2500 (Homo sapiens) | GSM2430088 | TAB-RRBS_GSC84FBS |
| GSE92469 | GSE92462 | GPL13534 | Illumina HumanMethylation450 BeadChip (HumanMethylation450_15017482) | GSM2430129 | 450K_GSC143 | GSE92460 | GPL16791 | Illumina HiSeq 2500 (Homo sapiens) | GSM2430060 | RRBS_GSC143 Sample 16 |
| GSE92469 | GSE92462 | GPL13534 | Illumina HumanMethylation450 BeadChip (HumanMethylation450_15017482) | GSM2430129 | 450K_GSC143 | GSE92460 | GPL16791 | Illumina HiSeq 2500 (Homo sapiens) | GSM2430070 | TAB-RRBS_GSC143 Sample 26 |
| GSE92469 | GSE92462 | GPL13534 | Illumina HumanMethylation450 BeadChip (HumanMethylation450_15017482) | GSM2430129 | 450K_GSC143 | GSE92460 | GPL16791 | Illumina HiSeq 2500 (Homo sapiens) | GSM2430080 | MAB-RRBS_GSC143 |
| GSE92469 | GSE92462 | GPL13534 | Illumina HumanMethylation450 BeadChip (HumanMethylation450_15017482) | GSM2430133 | 450K_NSC23 | GSE92460 | GPL16791 | Illumina HiSeq 2500 (Homo sapiens) | GSM2430045 | RRBS_NSC23 Sample 1 |

| | | | | | | | | | | |
|---|---|---|---|---|---|---|---|---|---|---|
| GSE92469 | GSE92462 | GPL13534 | Illumina HumanMethylation450 BeadChip (HumanMethylation450_15017482) | GSM2430133 | 450K_NSC23 | GSE92460 | GPL16791 | Illumina HiSeq 2500 (Homo sapiens) | GSM2430047 | TAB-RRBS_NSC23 Sample 3 |
| GSE92469 | GSE92462 | GPL13534 | Illumina HumanMethylation450 BeadChip (HumanMethylation450_15017482) | GSM2430133 | 450K_NSC23 | GSE92460 | GPL16791 | Illumina HiSeq 2500 (Homo sapiens) | GSM2430049 | MAB-RRBS_NSC23 Sample 5 |
| GSE57361 | GSE57360 | GPL13534 | Illumina HumanMethylation450 BeadChip (HumanMethylation450_15017482) | GSM2742407 | AD_1 | GSE57359 | GPL11154 | Illumina HiSeq 2000 (Homo sapiens) | GSM1380998 | Alzheimer's disease |
| GSE57361 | GSE57360 | GPL13534 | Illumina HumanMethylation450 BeadChip (HumanMethylation450_15017482) | GSM2742408 | DLB_01 | GSE57359 | GPL11154 | Illumina HiSeq 2000 (Homo sapiens) | GSM1380999 | Dementia of Lewy bodies |
| GSE103505 | GSE103502 | GPL16304 | Illumina HumanMethylation450 BeadChip [UBC enhanced annotation v1.0] | GSM2772516 | IMR-90 | GSE103503 | GPL20795 | HiSeq X Ten (Homo sapiens) | GSM2772525 | IMR90 |
| GSE103505 | GSE103502 | GPL16304 | Illumina HumanMethylation450 BeadChip [UBC enhanced annotation v1.0] | GSM2772517 | NA12878 | GSE103503 | GPL20795 | HiSeq X Ten (Homo sapiens) | GSM2772524 | NA12878 |
| GSE103505 | GSE103502 | GPL16304 | Illumina HumanMethylation450 BeadChip [UBC enhanced annotation v1.0] | GSM2772518 | IMR-90-2 | GSE103503 | GPL20795 | HiSeq X Ten (Homo sapiens) | GSM2772525 | IMR90 |
| GSE103505 | GSE103502 | GPL16304 | Illumina HumanMethylation450 BeadChip [UBC enhanced annotation v1.0] | GSM2772519 | NA12878-2 | GSE103503 | GPL20795 | HiSeq X Ten (Homo sapiens) | GSM2772524 | NA12878 |
| GSE103505 | GSE103502 | GPL16304 | Illumina HumanMethylatio | GSM2772520 | IMR-90-3 | GSE103503 | GPL20795 | HiSeq X Ten (Homo sapiens) | GSM2772525 | IMR90 |

| | | | | | | | | | | |
|---|---|---|---|---|---|---|---|---|---|---|
| | | | n450 BeadChip [UBC enhanced annotation v1.0] | | | | | | | |
| GSE103505 | GSE103502 | GPL16304 | Illumina HumanMethylation450 BeadChip [UBC enhanced annotation v1.0] | GSM2772521 | NA12878-3 | GSE103503 | GPL20795 | HiSeq X Ten (Homo sapiens) | GSM2772524 | NA12878 |
| GSE103505 | GSE103502 | GPL16304 | Illumina HumanMethylation450 BeadChip [UBC enhanced annotation v1.0] | GSM2772522 | IMR-90-4 | GSE103503 | GPL20795 | HiSeq X Ten (Homo sapiens) | GSM2772525 | IMR90 |
| GSE103505 | GSE103502 | GPL16304 | Illumina HumanMethylation450 BeadChip [UBC enhanced annotation v1.0] | GSM2772523 | NA12878-4 | GSE103503 | GPL20795 | HiSeq X Ten (Homo sapiens) | GSM2772524 | NA12878 |
| GSE211122 | GSE211120 | GPL13534 | Illumina HumanMethylation450 BeadChip (HumanMethylation450_15017482) | GSM6450945 | TC29-2 | GSE211121 | GPL20301 | Illumina HiSeq 4000 (Homo sapiens) | GSM6450983 | Sperm, Testicular Cancer, 12month [TC29-2] |
| GSE211122 | GSE211120 | GPL13534 | Illumina HumanMethylation450 BeadChip (HumanMethylation450_15017482) | GSM6450947 | HD15-3 | GSE211121 | GPL20301 | Illumina HiSeq 4000 (Homo sapiens) | GSM6450982 | Sperm, Hodgkin disease, 18month [HD15-3] |
| GSE211122 | GSE211120 | GPL13534 | Illumina HumanMethylation450 BeadChip (HumanMethylation450_15017482) | GSM6450951 | TC29-3 | GSE211121 | GPL20301 | Illumina HiSeq 4000 (Homo sapiens) | GSM6450984 | Sperm, Testicular Cancer, 18month [TC29-3] |
| GSE211122 | GSE211120 | GPL13534 | Illumina HumanMethylation450 BeadChip (HumanMethylation450_15017482) | GSM6450970 | CC33-2 | GSE211121 | GPL20301 | Illumina HiSeq 4000 (Homo sapiens) | GSM6450977 | Sperm, Control1, 12month [CC33-2] |

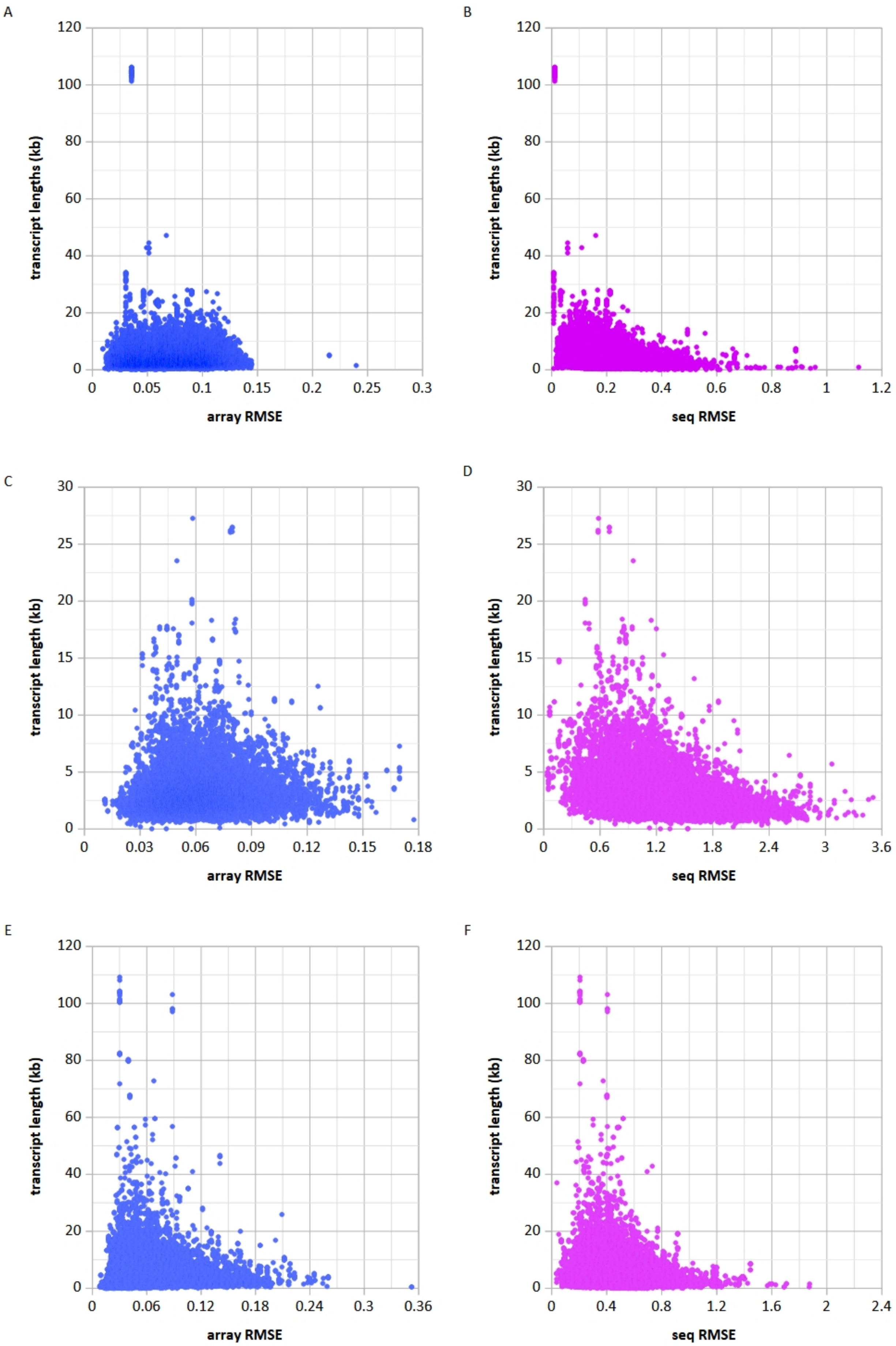
A
120
100
80
60
40
20
0
transcript lengths (kb)
0
0.05
0.1
0.15
0.2
0.25
0.3
array RMSE
B
120
100
80
60
40
20
0
transcript lengths (kb)
0
0.2
0.4
0.6
0.8
1
1.2
seq RMSE
C
30
25
20
15
10
5
0
transcript length (kb)
0
0.03
0.06
0.09
0.12
0.15
0.18
array RMSE
D
30
25
20
15
10
5
0
transcript length (kb)
0
0.6
1.2
1.8
2.4
3
3.6
seq RMSE
E
120
100
80
60
40
20
0
transcript length (kb)
0
0.06
0.12
0.18
0.24
0.3
0.36
array RMSE
F
120
100
80
60
40
20
0
transcript length (kb)
0
0.4
0.8
1.2
1.6
2
2.4
seq RMSE

**Figure S1**: Root mean square errors (RMSE) for X-Plat-predicted microarray and RNA-seq expression values (array RMSE and seq RMSE) in rat (A–B), Arabidopsis (C–D), and human (E–F) visualized as scatterplots of array RMSE vs transcript length (A, C, E) and seq RMSE vs transcript length (B, D, F).

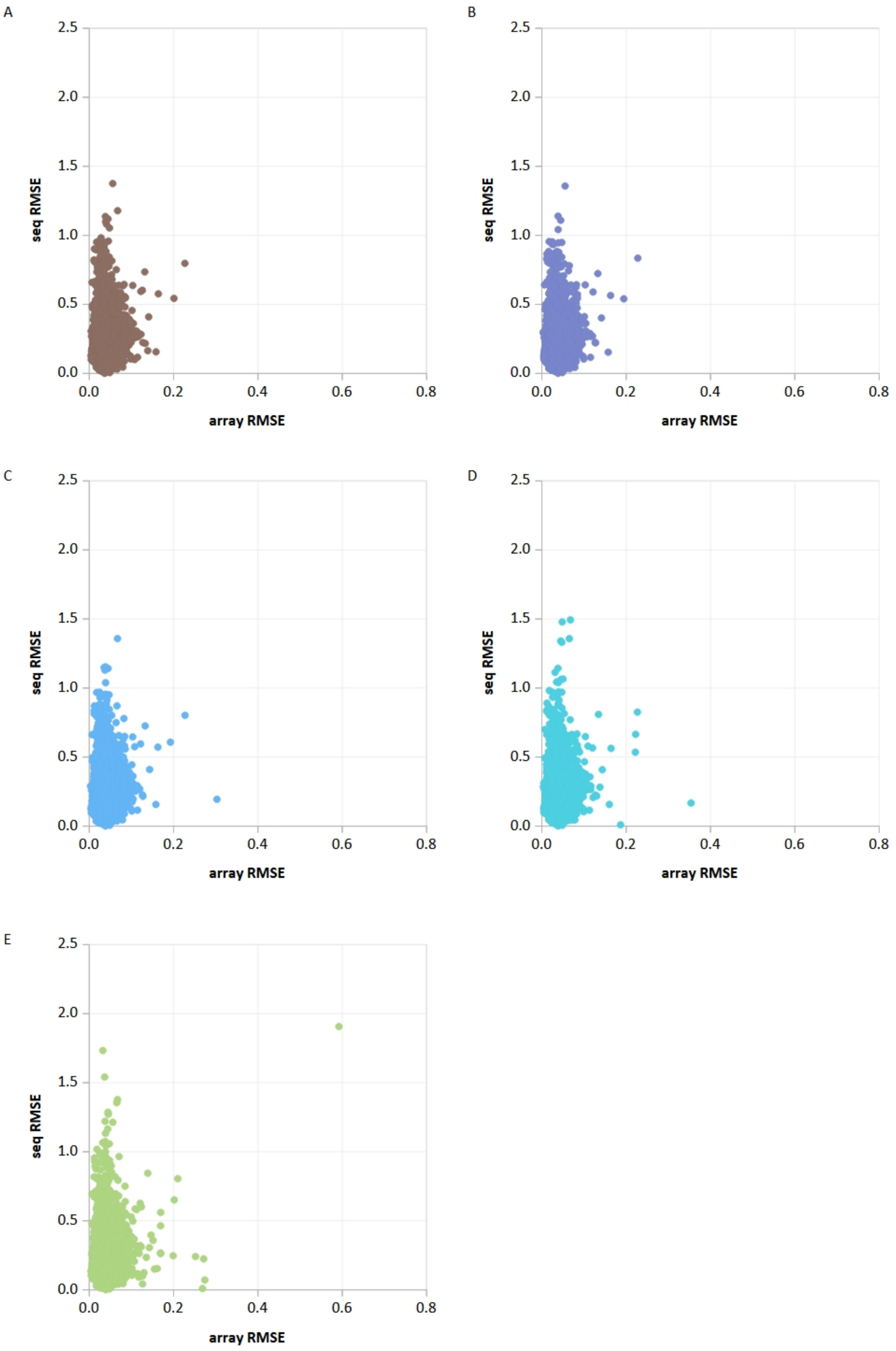
A
B
C
D
E
seq RMSE
array RMSE
0.0
0.5
1.0
1.5
2.0
2.5
0.2
0.4
0.6
0.8

**Figure S2**: Effect of training-set downsampling on X-Plat performance for rat expression data. Scatterplots of array RMSE vs seq RMSE for X-Plat-predicted values in the rat dataset when the model was trained on (A) the full training set (no downsampling), or on random subsets comprising (B) 90%, (C), 70% (D), 50%, and (E) 30% of the original training samples, with the same held-out test samples used in all panels.
